# Arsenal of Nanobodies for Broad-Spectrum Countermeasures against Current and Future SARS-CoV-2 Variants of Concerns

**DOI:** 10.1101/2021.12.20.473401

**Authors:** M. A. Rossotti, H. van Faassen, A. Tran, J. Sheff, J. K. Sandhu, D. Duque, M. Hewitt, S. Wen, R. Bavananthasivam, S. Beitari, K. Matte, G. Laroche, P. M. Giguère, C. Gervais, M. Stuible, J. Guimond, S. Perret, G. Hussack, M.-A. Langlois, Y. Durocher, J. Tanha

## Abstract

Nanobodies offer several potential advantages over mAbs for the control of SARS-CoV-2. Their ability to access cryptic epitopes conserved across SARS-CoV-2 variants of concern (VoCs) and feasibility to engineer modular, multimeric designs, make these antibody fragments ideal candidates for developing broad-spectrum therapeutics against current and continually emerging SARS-CoV-2 VoCs. Here we describe a diverse collection of 37 anti-SARS-CoV-2 spike glycoprotein nanobodies extensively characterized as both monovalent and IgG Fc-fused bivalent modalities. The panel of nanobodies were shown to have high intrinsic affinity; high thermal, thermodynamic and aerosolization stability; broad subunit/domain specificity and cross-reactivity across many VoCs; wide-ranging epitopic and mechanistic diversity; high and broad *in vitro* neutralization potencies; and high neutralization efficacies in hamster models of SARS-CoV-2 infection, reducing viral burden by up to six orders of magnitude to below detectable levels. *In vivo* protection was demonstrated with anti-RBD and previously unreported anti-NTD and anti-S2 nanobodies. This collection of nanobodies provides a therapeutic toolbox from which various cocktails or multi-paratopic formats could be built to tackle current and future SARS-CoV-2 variants and SARS-related viruses. Furthermore, the high aerosol-ability of nanobodies provides the option for effective needle-free delivery through inhalation.

## INTRODUCTION

Declared a pandemic in March 2020 by the World Health Organization (covid19.who.int), coronavirus disease 2019 (COVID-19), caused by severe acute respiratory syndrome coronavirus 2 (SARS-CoV-2), remains a severe global health and economic burden. As of 20 December, 2021, 275 million individuals have been infected world-wide, of which ∼5.5 million have died (coronavirus.jhu.edu). The toll on public health has been exacerbated with the continual emergence of SARS-CoV-2 variants of concern (VoCs) ^1, 2^. These VoCs, which include Alpha (B.1.1.7), Beta (B.1.351), Gamma (P.1), Delta (B.1.617.2) and Omicron (B.1.1.529), can evade COVID-19 vaccines and therapeutics to different extents, and the evolutionary trajectory of the virus variants predicts newer VoC escape mutants to emerge in the future ^1–8^.

Key to SARS-CoV-2 infection is its surface-displayed spike glycoprotein (S) ^9–13^, a homotrimeric protein where each protomer ectodomain format consists of S1 and S2 subunits. S1 is further delineated by an N-terminal domain (NTD), a receptor-binding domain (RBD) and subdomains SD1 and SD2. The spike glycoprotein mediates cell entry, a critical first phase in the infection process, involving two discrete but concerted steps. In the first, virus-cell binding step, the RBD, essentially through its receptor-binding motif (RBM), binds to its host receptor angiotensin-converting enzyme II (ACE2). This is followed by the second, virus-cell fusion step, which is mediated by the S2 subunit and concludes the viral cell entry event. Spike glycoprotein is the primary target for COVID-19 therapeutic antibodies, which operate by stopping virus cell entry *via* blocking the cell binding and/or fusion step. In particular, the mechanism of action of most potent neutralizing antibodies involves binding to the RBD, although neutralizing antibodies targeting the NTD domain ^14–19^ and the S2 subunit ^20, 21^ have also been reported.

While many COVID-19 immunotherapies are based on monoclonal antibodies (mAbs), single-domain antibodies (mostly V_H_Hs) are also being pursued as alternative therapeutics ^22–48^. V_H_Hs (nanobodies) are the variable domains of camelid heavy-chain antibodies responsible for antigen recognition. One nanobody (VHH-72/XVR011) has already entered clinical trials for COVID-19 therapy ^29, 39, 49^. V_H_Hs offer potential advantages over mAbs as COVID-19 immunotherapeutics, most notably because of their stability against aerosolization that allows for convenient, low-cost and effective needle-free delivery of V_H_Hs into the key site of infection (lungs) by inhalation ^40, 41, 50–52^. Importantly, V_H_Hs permit modular assembly of multimeric/multi-paratopic nanobody constructs with drastically improved efficacy and cross-reactivity/neutralization breadth across VoCs ^34, 36^. Multispecific V_H_H constructs can also be designed to target confined geometric spaces on the surface of the target antigen without nanobody clash, a feature not achievable with larger mAbs. Critically, with small size and frequently extended CDR3s, V_H_Hs can reach cryptic epitopes that are hidden from mAbs and conserved across SARS-CoV-2 VoCs, allowing for the development of broad-spectrum nanobody therapeutics against current and future VoCs ^34, 39, 40^.

Here we report the isolation, extensive characterization and *in vivo* efficacy of a large panel of SARS-CoV-2-targeting nanobodies. Monovalent V_H_H and bivalent V_H_H-Fc formats were assessed for binding affinity; thermal, thermodynamic and aerosol stability; epitopic diversity; S subunit/domain specificity; cross-reactivity to multiple betacoronavirus subgenera and VoCs; *in vitro* cross-neutralization potencies against several VoCs; and *in vivo* neutralization efficacies using a hamster model of infection. Multiple neutralization mechanisms of action are possible through V_H_H binding to RBD, NTD, and S2, including inhibiting the virus-cell binding and/or fusion steps. This robust collection of nanobodies provides a foundation for development of effective broad-spectrum therapeutics (monotherapy, cocktails or multimerics/multi-paratopics) that could tackle current and future SARS-CoV-2 variants and SARS-related viruses.

## 2. RESULTS

### 2.1 Llama immunization and serum analyses

Prior to immunization, serology and panning experiments, purified SARS-CoV-2 spike glycoprotein (S) antigens were validated for functionality in adsorbed/captured states on microtiter wells (**Fig. S1; Table S1**). Two llamas (Green & Red) were immunized with SARS-CoV-2 Wuhan-Hu-1 (Wuhan) S fragments. Specifically, Green was primed with S and boosted with three doses of RBD fragment, while Red received four doses of S. Both llamas produced a strong and specific immune response to S, S1, S2 and RBD with Green consistently outperforming Red (up to 10-fold) across all four target proteins (**Fig. S2A**). Analyses of total polyclonal sera by flow cytometry-based surrogate neutralization assays (SVNA) showed a more potent neutralizing antibody response generated by Green (**Fig. S2B**).

### 2.2 Phage display library construction, selection and screening

Two phage display libraries, Green and Red, were constructed using day 28 peripheral blood mononuclear cells (PBMCs) and separately subjected to two rounds of panning against S fragments. To further maximize for V_H_H diversity, panning was performed under multiple selection conditions (P1 – P6; see Methods) to direct selection towards S-, S1-, S2-, RBD-, NTD- and RBM-specific binders. Monoclonal phage ELISA combined with DNA sequencing identified 37 unique V_H_Hs across all screens (**Fig. 1A; Fig. 1B**). Most V_H_Hs originated from llama Green, 26 *vs* 11 from llama Red (**Fig. 1A**). Llama Green, which was predominantly immunized with RBD, yielded a higher proportion of RBD-specific V_H_Hs (15 RBD, six NTD, five S2) compared to llama Red which was immunized with only S and yielded mostly S2-specific V_H_Hs (two RBD, three NTD, six S2).

**Fig. 1.**
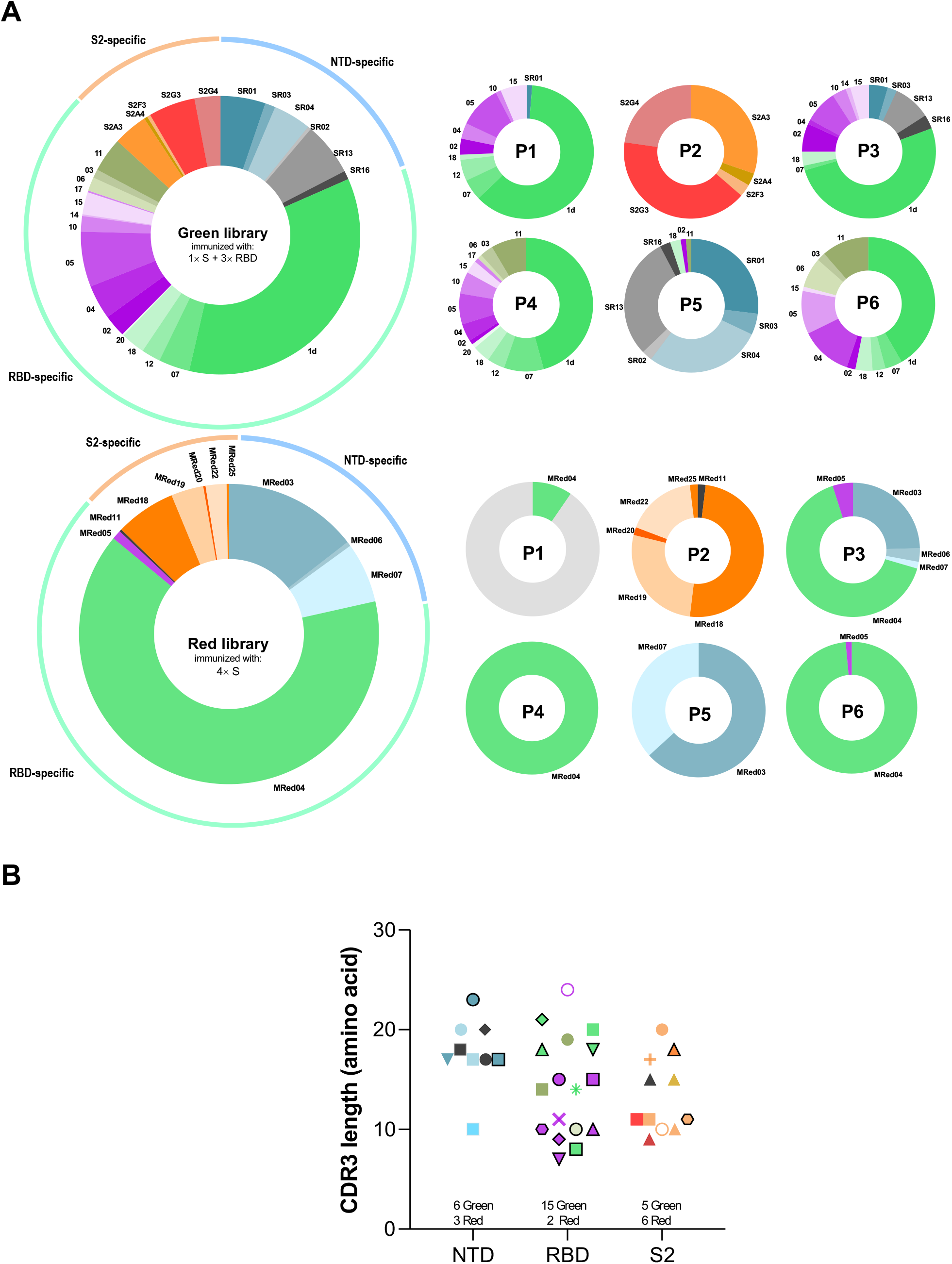
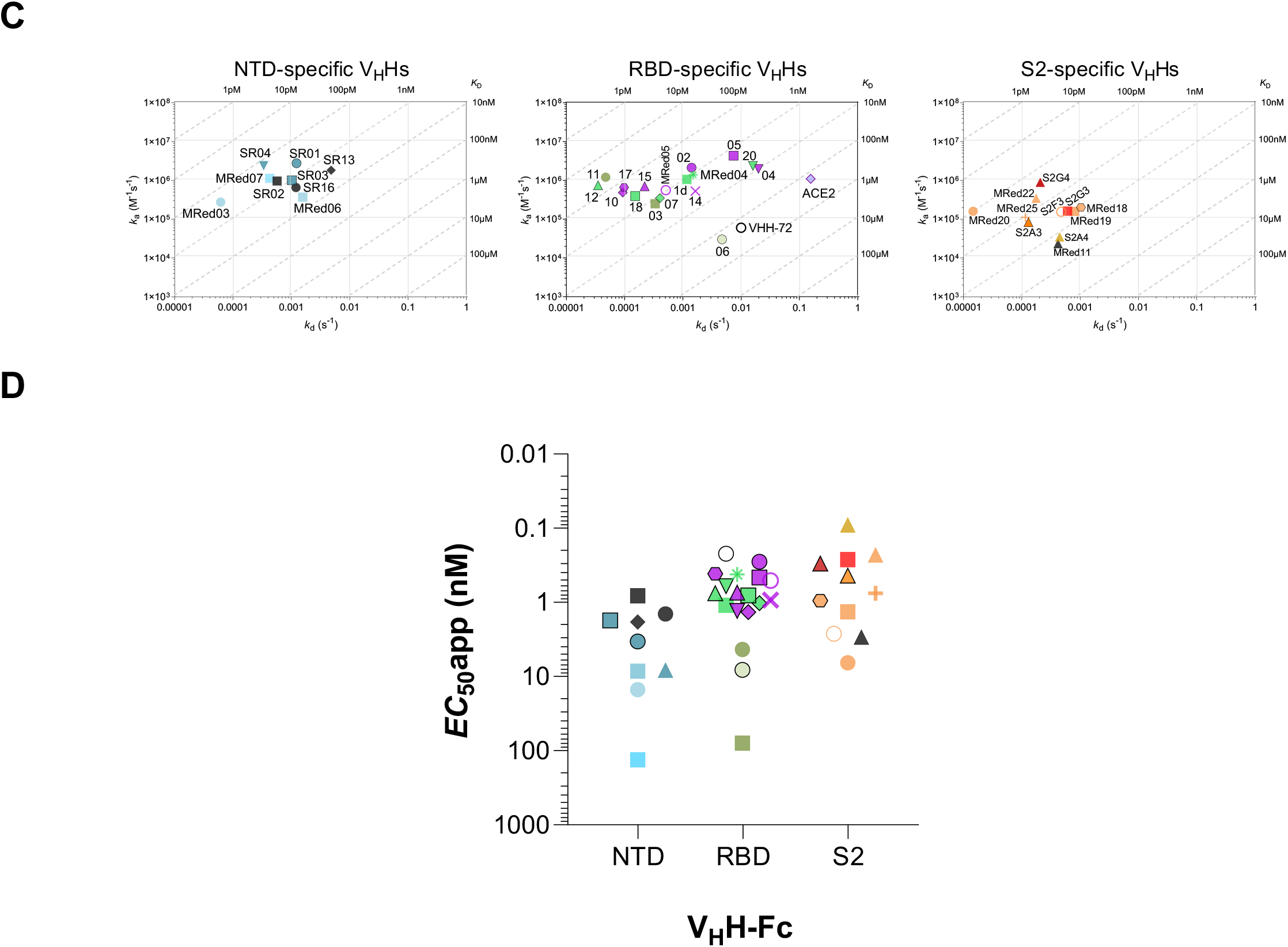
Selection, binding affinity and subunit/domain specificity of anti-SARS-CoV-2 S V_H_Hs. V_H_Hs are grouped based on their specificity for NTD, RBD or S2 and color-coded based on their epitope bin designation (*see* **Fig. 5A**). **A**) Library origin, relative proportion and subunit/domain specificity of the 37 anti-SARS-CoV-2 S V_H_Hs selected from the Green and Red libraries by employing six panning strategies P1 – P6. Binding specificity patterns in P1 – P6 reflect the selection strategies employed. For the Red library, P1 also selected for a V_H_H (ARes09, grey, unlabeled) that was specific to the resistin trimerization domain of the S used for immunization and panning. Green library V_H_Hs were isolated from the llama immunized once with S and three times with RBD. Red library V_H_Hs were isolated from the llama immunized four times with S. **B**) Diversity of the 37 SARS-CoV-2 S V_H_Hs shown in terms of CDR3 length. The number of V_H_Hs derived from each of the two libraries are shown. **C**) On-/off-rate maps summarizing V_H_H kinetic rate constants, *k*_a_s and k_d_s, determined by SPR. Diagonal lines represent equilibrium dissociation constants, *K*_D_s (*see* also **Table 1**). Maps were constructed using the V_H_H binding data (**Fig. S3; Table S2**) against SARS-CoV-2 Wuhan S (all except 12 and MRed05) or RBD/SD1 (12 and MRed05). V_H_H subunit/domain specificities were determined by SPR and ELISA (**Fig. S3**; **Table S2**; **Table S3**). Anti-SARS-CoV S VHH-72 which cross-reacts with SARS-CoV-2 RBD ^29^ and the monomeric ACE2 (ACE2-H_6_) are included as benchmark/reference binders. **D**) Binding of SARS-CoV-2 S V_H_H-Fcs to S- expressing CHO cells (CHO-SPK) obtained by flow cytometry. Apparent *EC*_50_s (*EC*_50_apps) were obtained from graphs in **Fig. S4** and are included in **Table 1**. Black open circle, VHH-72 benchmark.

Selected V_H_Hs were then (i) cloned as fusions to the biotinylation acceptor peptide (BAP) and His_6_ tags and produced in *E. coli*; and (ii) cloned in fusion with human IgG1 hinge-Fc domain (V_H_H-Fc) and produced in HEK293-6E cells.

### 2.3. Binding characteristics of V_H_Hs and V_H_H-Fcs

V_H_Hs/V_H_H-Fcs were tested by SPR and ELISA against recombinant Wuhan SARS-CoV-2 S, S1, RBD, NTD and S2 proteins to determine affinities and subunit/domain specificities (**Fig. 1C; Fig. S3; Table S2; Table S3**). V_H_Hs bound with high affinity, with the majority of *K*_D_s in the single-digit-nM to pM range. Three clusters of V_H_Hs were identified: 17 RBD-specific V_H_Hs, nine NTD-specific V_H_Hs (no reactivity to RBD, bound S and S1) and 11 S2-specific V_H_Hs (**Fig. 1C**). The domain specificity of the NTD binders was confirmed in subsequent ELISAs (**Fig S3B; Table S3**). By flow cytometry, V_H_H-Fcs bound SARS-CoV-2 S (Wuhan) in a more natural context on the cell membrane of CHO cells (CHO^55E1™^) stably transfected with the S protein (**Fig. 1D; Fig. S4; Table 1**). High apparent affinities (*EC*_50_apps) in the single-digit-nM to pM range were observed for the majority of V_H_H-Fcs.

**Table 1.** Binding characteristics of SARS-CoV-2 V_H_Hs

| V<sub>H</sub>H/ACE2 | Epitope |  | SARS-CoV-2 Wuhan S | SARS-CoV-2 Alpha S |  | SARS-CoV-2 Beta S |  | SARS-CoV S |  | SARS-CoV-2 S expressing cells<sup>9</sup> |
| --- | --- | --- | --- | --- | --- | --- | --- | --- | --- | --- |
|  | bin | type | SPR (K<sub>D</sub>, nM) | ELISA<sup>6</sup> | SPR (K<sub>D</sub>, nM) | ELISA<sup>e</sup> | SPR (K<sub>D</sub>, nM) | ELISA<sup>6</sup> | SPR (K<sub>D</sub>, nM) | FC (EC<sub>50</sub>app nM) |
| **RBD-specific V<sub>H</sub>H** |  |  |  |  |  |  |  |  |  |  |
| 1d | 1 | Conf.<sup>c</sup> | 0.75 | + | 0.91 | + | 1.2 | + | - | 1.1 |
| 02 | 2/3 | Conf.<sup>c</sup> | 0.62 | + | 13.6 | - | - | - | - | 0.3 |
| 03 | 6 | Conf.<sup>c</sup> | 1.56 | + | 1.49 | + | 4.08 | - | - | 79 |
| 04 | 4 | Conf.<sup>c</sup> | 10.2 | + | 11.7 | + | - | - | - | 1.3 |
| 05 | 2/3 | Conf.<sup>c</sup> | 2.6 | + | 11.4 | - | - | - | - | 0.5 |
| 06 | 5 | Linr.<sup>d</sup> | 223 | + | 229 | + | 248 | - | - | 8.1 |
| 07 | 1 | Conf.<sup>c</sup> | 0.94 | + | 1.1 | + | 1.1 | + | 12.2 | 1 |
| 10 | 2/3/4 | Conf.<sup>c</sup> | 0.2 | + | 0.21 | + | 9.73 | - | - | 1.3 |
| 11 | 6 | Conf.<sup>c</sup> | 0.018 | + | 0.017 | + | 0.023 | + | 0.014 | 4.3 |
| 12<sup>a</sup> | 1 | Conf.<sup>c</sup> | 0.047 | + | 0.046 | + | 0.04 | + | nd | 0.8 |
| 14 | 2/4 | Conf.<sup>c</sup> | 2.6 | + | 2.44 | + | wb<sup>f</sup> | - | - | 0.9 |
| 15 | 2/3/4 | Conf.<sup>c</sup> | 0.32 | + | 0.31 | + | 22.2 | - | - | 0.7 |
| 17 | 3/4 | Conf.<sup>c</sup> | 0.15 | + | 0.13 | + | 5.1 | - | - | 0.4 |
| 18 | 1 | Linr.<sup>d</sup> | 0.32 | + | 0.35 | + | 0.37 | - | - | 0.8 |
| 20 | 1 | Conf.<sup>c</sup> | 4.39 | + | 4.97 | + | 5.47 | + | - | 0.6 |
| MRed04 | 1 | Conf.<sup>c</sup> | 0.86 | + | 0.91 | + | 1.07 | + | 300 | 0.4 |
| MRed05<sup>a</sup> | 2/3/4 | Conf.<sup>c</sup> | 0.91 | + | 0.31 | + | 0.89 | - | nd | 0.5 |
| VHH-72<sup>b</sup> | 1 | Conf.<sup>c</sup> | 86.2 | + | 96 | + | 124 | + | 6.52 | 0.2 |
| ACE2<sup>b</sup> | na<sup>d</sup> | na | 153 | + | 18.3 | + | 131 | + | 351 | 1.2 |
| **NTD-specific V<sub>H</sub>H** |  |  |  |  |  |  |  |  |  |  |
| SR01 | 7 | Linr.<sup>d</sup> | 0.56 | + | 0.59 | + | 0.2 | + | 0.154 | 3.4 |
| SR02 | nd | Conf.<sup>c</sup> | 0.14 | + | 0.06 | + | 0.15 | + | - | 0.8 |
| SR03 | 7 | Conf.<sup>c</sup> | 1.69 | + | 1.72 | + | 2.49 | - | - | 1.7 |
| SR04 | 7 | Linr.<sup>d</sup> | 0.14 | + | 0.27 | + | 0.32 | - | - | 8.2 |
| SR13 | nd | Conf.<sup>c</sup> | 3.6 | + | 5.8 | + | 7 | - | - | 1.8 |
| SR16 | nd | Conf.<sup>c</sup> | 2 | + | 1.6 | + | 2.6 | - | - | 1.4 |
| MRed03 | 8 | Conf.<sup>c</sup> | 0.51 | + | 0.36 | + | 0.67 | - | - | 15 |
| MRed06 | 8 | Conf.<sup>c</sup> | 5.2 | + | 5.72 | + | 7.24 | - | - | 8.5 |
| MRed07 | 9 | Conf.<sup>c</sup> | 0.11 | + | 0.26 | + | 0.23 | - | - | 132 |
| **S2-specific V<sub>H</sub>H** |  |  |  |  |  |  |  |  |  |  |
| S2A3 | 10 | Linr.<sup>d</sup> | 0.56 | + | 2.18 | + | 0.85 | - | - | 0.4 |
| S2A4 | 11 | Linr.<sup>d</sup> | nd | + | nd | + | nd | - | - | 0.1 |
| S2F3 | 12 | Linr.<sup>d</sup> | nd | + | nd | + | nd | + | 4.9 | 2.7 |
| S2G3 | 13 | Linr.<sup>d</sup> | 1.87 | + | 1.78 | + | 1.85 | + | 4.27 | 0.3 |
| S2G4 | 14 | Linr.<sup>d</sup> | 0.23 | + | nd | + | nd | + | 0.8 | 0.3 |
| MRed11 | nd | Linr.<sup>d</sup> | 6.2 | + | 13.7 | + | 6.2 | - | - | 2.9 |
| MRed18 | 12 | Linr.<sup>d</sup> | 6.03 | + | 12.9 | + | 6.48 | + | 22.5 | 0.9 |
| MRed19 | 12 | Linr.<sup>d</sup> | 9.07 | + | 20.2 | + | 8.07 | + | 24.6 | 1.3 |
| MRed20 | 12 | Linr.<sup>d</sup> | nd | + | nd | + | nd | + | 10.7 | 4.7 |
| MRed22 | 12 | Conf.<sup>c</sup> | nd | + | nd | + | nd | - | - | 0.3 |
| MRed25 | 12 | Conf.<sup>c</sup> | nd | + | nd | + | nd | + | 2.29 | 0.8 |

<sup>a</sup>K<sub>DS</sub> were determined by flowing monomeric V<sub>HH</sub>s over sensorchip surfaces immobilized with S, except for V<sub>HH</sub> 12 and MRed05, which were determined by flowing monomeric RBDs over V<sub>HH</sub>-Fc-captured surfaces (see **Table S4**); <sup>b</sup>V<sub>HH</sub>-72 benchmark is a SARS-CoV S-specific V<sub>HH</sub> that cross-reacts with SARS-CoV-2 S[<sup>1</sup>]; <sup>c</sup>His<sub>6</sub>-tagged monomeric ACE2 (ACE2-H<sub>6</sub>) was used for SPR assays, human Ig Fc-fused dimeric ACE2 (ACE2-Fc) for ELISA and cell binding assays by flow cytometry; <sup>d</sup>Conf., conformational epitope; <sup>e</sup>Linr., linear epitope; <sup>f</sup>ELISA was performed at 125 nM (1 µg/mL) V<sub>HH</sub>-Fc concentration (see **Fig.2A**); <sup>g</sup>wb, weak binding; <sup>h</sup>cell binding was performed by flow cytometry (FC) using V<sub>HH</sub>-Fc; na, not applicable; nd, not determined. <sup>1</sup>Wrapp D, *et al.* Structural Basis for Potent Neutralization of Betacoronaviruses by Single-Domain Camelid Antibodies. *Cell* **181**, 1004-1015 e1015 (2020).

V_H_Hs were examined for cross-reactivity to a collection of spike glycoprotein fragments from various coronavirus genera and SARS-CoV-2 variants by ELISA and SPR. In ELISA (**Fig. 2A**; **Table 1**), many V_H_H-Fcs cross-reacted with the S protein from VoCs Alpha, Beta, Gamma, Delta and Kappa (B.1.617.1; Variant Being Monitored [VBM]). The exceptions were: 1) RBD-specific V_H_Hs 02/05 and 04/14/15 did not cross-react with Beta and Gamma, and Kappa, respectively, and 2) S2-specific V_H_Hs MRed18 and MRed19 did not cross-react with Kappa. All nine NTD-specific V_H_Hs cross-reacted with all variants tested. Additionally, many V_H_Hs cross-reacted with pangolin CoV, with fewer cross-reacting to SARS-CoV, SARS-like CoV WIV1, bat SARS-like CoV and civet SARS CoV. These viruses, including variants, are all of the Betacoronavirus Sarbecovirus subgenus. None of the antibodies tested cross-reacted with the remaining 11 non-Sarbecovirus Betacoronavirus, Alphacoronavirus, Deltacoronavirus or Gammacoronavirus. The broadly cross-reactive antibodies included V_H_Hs targeting all three regions of the S protein (RBD, NTD, S2). The most broadly cross-reactive V_H_Hs recognizing 10 – 11 viruses, including SARS-CoV-2 variants, were two NTD binders (SR01, SR02), six RBD binders (1d, 07, 11, 12, 20, MRed04) and six S2 binders (S2F3, S2G3, S2G4, MRed18, MRed19, MRed20). The VHH-72 benchmark was also broadly cross-reactive. The panel of V_H_Hs had similar cross-reactivity profiles to human ACE2, except that ACE2 did not bind civet SARS-CoV S and, unsurprisingly, bound HCoV-NL63 S (**Fig. 2A**) ^53, 54^.

**Fig. 2.**
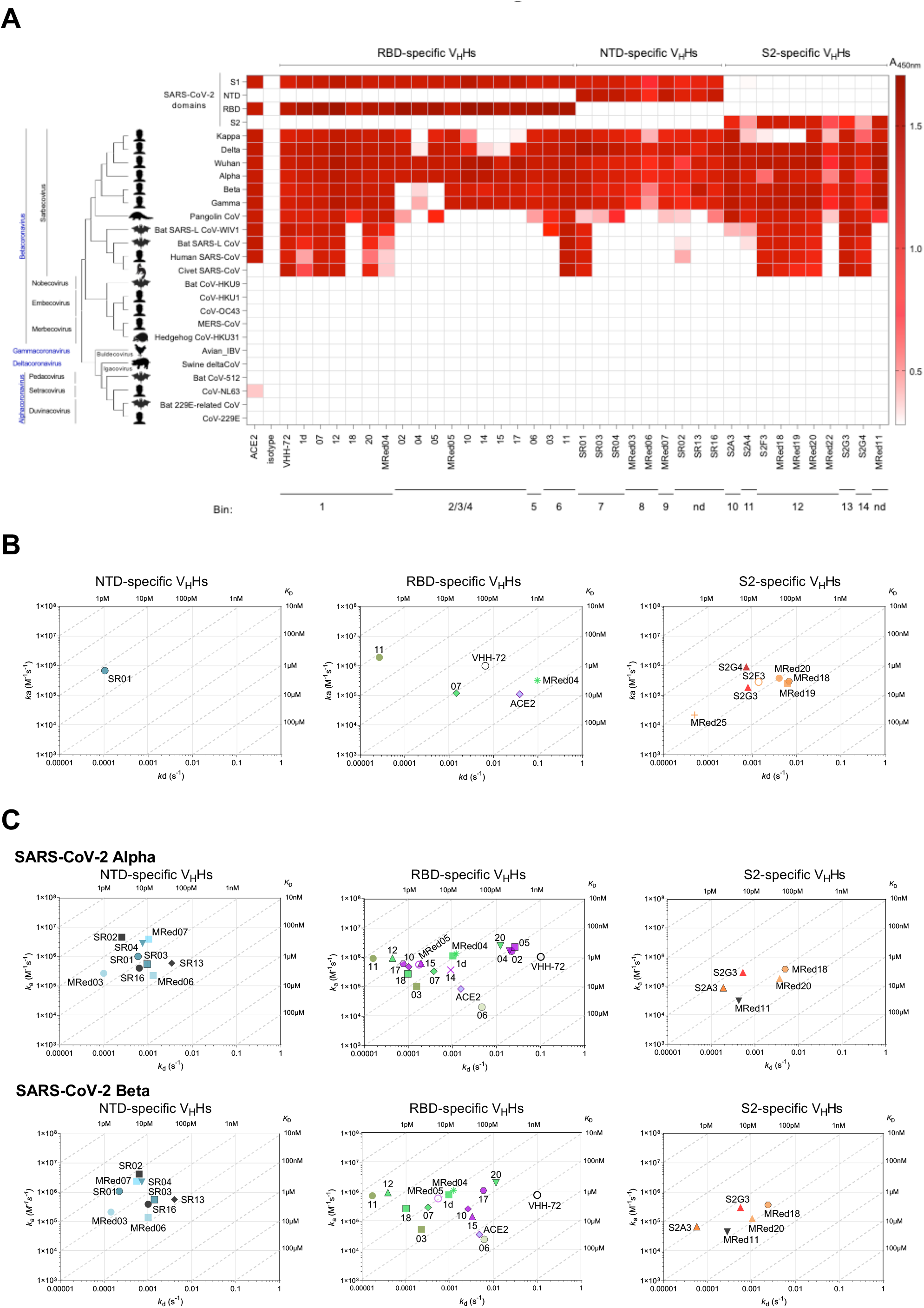
Cross-reactivity of anti-SARS-CoV-2 S V_H_Hs. Data are organized based on V_H_H subunit/domain specificity and epitope bin designation (*see* **Fig. 5A**). **A**) ELISA showing the cross-reactivity of V_H_Hs against various coronavirus spike glycoprotein fragments, S, S1, S2, RBD and NTD. Shades of red represent binding, colorless boxes represent no binding. Assays were performed at a single V_H_H-Fc concentration. The “isotype” control (A20.1 V_H_H-Fc) shows no binding to S. Anti-SARS-CoV VHH-72 and ACE2-Fc were included as references. The phylogenetic tree of spike glycoproteins was constructed using MEGA11^95^. **B, C**) On-/off-rate maps summarizing V_H_H kinetic rate constants, *k*_a_s and *k*_d_s determined by SPR for the binding of V_H_Hs to SARS-CoV S (**B**) and SARS-CoV-2 Alpha and Beta S (**C**). Diagonal lines represent equilibrium dissociation constants, *K*_D_s (*see* also **Table 1**). Maps were constructed using the V_H_H binding data from **Fig. S5** and **Table S4**. Anti-SARS-CoV VHH-72 and the monomeric ACE2 (ACE2-H_6_) are included as benchmark/reference binders.

When tested by SPR against SARS-CoV, 11 out of 14 ELISA-positive V_H_Hs cross-reacted with SARS-CoV S, most with comparably high affinities (**Fig. 2B; Table 1; Fig. S5A; Table S4**). Seven of these V_H_Hs were S2-specific, three RBD-specific and one NTD-specific. Against the Alpha and Beta variants, the SPR cross-reactivity data, performed with all RBD, all NTD and five S2 V_H_Hs (31 V_H_Hs), were consistent with ELISA, except for 04 and 14 which were negative or very weak for binding to the Beta variant by SPR. All 31 V_H_Hs bound the Alpha variant S protein, 28 of which were also cross-reactive to the Beta variant S protein (**Fig. 2C; Fig S5B**; **Table 1; Table S4**). Thirteen out of 17 RBD-specific V_H_Hs bound all three variants with similar affinities, except for V_H_Hs 10, 15 and 17 which bound to the Beta variant with 40 – 50-fold weaker affinity; the remaining four that did not bind the Beta variant showed cross-reactivity with the Alpha variant with similar (04, 14) or reduced (∼5-fold [05] and ∼20-fold [02]) affinity relative to the Wuhan variant. All nine NTD-specific and five S2-specific V_H_Hs cross-reacted with the three variants with essentially the same affinities. The loss of binding for some of the ELISA-positive nanobodies in SPR assays could be due to the loss of binding avidity (V_H_H-Fc was used in ELISA *vs* V_H_H in SPR) and/or epitope hindrance on the sensorchip. In agreement with this, V_H_H 20 (ELISA^+^/SPR^-^) neutralizes SARS-CoV in surrogate virus neutralization assays (*see* below) as a V_H_H-Fc format; so do V­H-Fc 18 and SR13 which were false negatives by both ELISA and SPR, additionally demonstrating the number of SARS-CoV cross-reacting V_H_Hs exceeds that gleaned from the ELISA and SPR data.

### 2.4. Stability characteristics of V_H_Hs

By size exclusion chromatography (SEC), all tested V_H_Hs were aggregation-resistant (**Fig. 3A; Fig. S6**). V_H_Hs were highly thermostable: with the exception of V_H_H 11, MRed25 and S2A3 and which had *T*_m_s of ∼ 60 – 61°C, the remaining 30 V_H_Hs had higher *T*_m_s of ∼65 – 80°C (median: 70.4°C) (**Fig. 3B; Fig. S7; Table S5**). Conformational stability of V_H_Hs was determined by measuring free energy of unfolding (Δ*G*^0^) in GdnHCl equilibrium denaturation experiments, with Δ*G*^0^ ranging from 21.4 – 53.4 kJ/mol (median: 30.7 kJ/mol), and an *m* value range of 10.3 – 19.8 kJ/M*mol (median: 14.6 kJ/M*mol) observed (**Fig. 3C**; **Fig. S8; Table S5**)^55–59^.

**Fig. 3.**
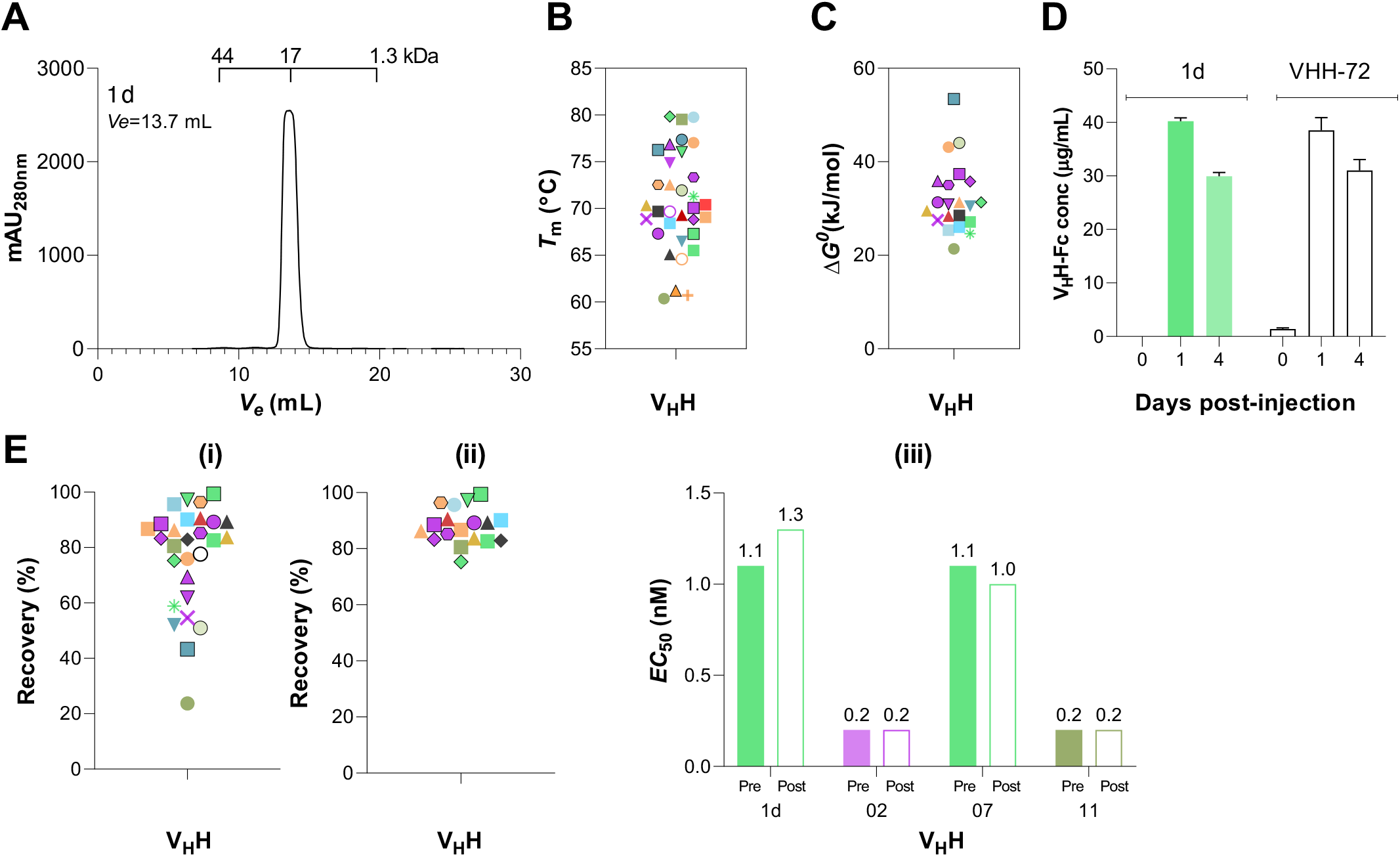
Stability of anti-SARS-CoV-2 S V_H_Hs. **A**) Representative SEC profile demonstrating the aggregation resistance of V_H_Hs. See **Fig. S6** for the full dataset. **B**) Summary of V_H_H *T*_m_ data. *T*_m_s were obtained from plots of % folded *vs* temperature (**Fig. S7; Table S5**). **C**) Summary of V_H_H Δ*G*^0^ data. Δ*G*^0^ (as well as other thermodynamic parameters, *C*_m_ and *m* values) are reported in **Table S5**. **D**) *In vivo* stability and persistence of V_H_Hs. Stability and persistence were determined by monitoring the concentration of a representative V_H_H-Fc (1d) in hamster blood at various days post-injection by ELISA. VHH72-Fc was used as the benchmark. **E**) Stability of V_H_Hs against aerosolization. Summary of % recovery of all (**i**) and lead (**ii**) V_H_Hs are shown. Percent recovery represents the proportion of a V_H_H that remained soluble monomer following aerosolization. Graphs were generated based on the data in **Fig. S9A** and **Table S6**. Open circle in (**i**) represents benchmark VHH-72. (**iii**) Activity of pre- *vs* post-aerosolized V_H_Hs expressed in terms of antigen binding (*EC*_50_). *EC*_50_s were determined by ELISA. V_H_Hs in “**B**”, “**C**”, “**E(i)**” and “**E(ii)**”are color-coded based on their epitope bin designation (*see* **Fig. 5A**).

Since we planned to test V_H_H-Fcs in hamsters for *in vivo* efficacy, we pre-emptively assessed their *in vivo* stability and persistence. We chose 1d V_H_H-Fc as a representative and included VHH-72 V_H_H-Fc, whose modified/enhanced version is currently in a phase 1 clinical trial, as a reference. Hamsters were injected intraperitoneally (IP) with 1 mg of each antibody and serum antibody concentration was monitored for up to four days by ELISA. Significant and comparable V_H_H-Fc concentrations were present in the hamster sera for both 1d and VHH-72 V_H_H-Fcs on days 1 and 4 post injection (**Fig. 3D**), indicating V_H_H-Fcs would have the required serum stability and persistence *in vivo* for the duration of the animal studies.

V_H_Hs were also examined for their aggregation resistance and stability upon aerosolization. For a few V_H_Hs, aerosolization induced soluble aggregate formation as determined by SEC, while for others it led to the formation of visible aggregates (**Fig. S9; Table S6**). This resulted in reduced % recoveries, measures of V_H_H stability against aerosolization, and corresponding to the proportion of V_H_Hs that remained as soluble monomer following aerosolization (**Fig. 3E; Fig. S9; Table S6**). The majority of V_H_Hs (18 out of 28 V_H_Hs tested), however, were stable against aerosolization with high % recoveries (**Fig. 3E**). Additionally, several V_H_Hs still showed a high % recovery upon aerosolization (50 – 70 %) despite the formation of some visible aggregates. Comparison of ELISA-derived *EC*_50_s of select pre-aerosolized *vs* post-aerosolized V_H_Hs clearly demonstrated aerosolization did not compromise the binding activities of V_H_Hs (**Fig. 3E**; **Fig. S9**).

### 2.5. Screening for neutralizing V_H_Hs by surrogate virus neutralization assays (SVNAs)

A preliminary screen of 25 V_H_Hs (14 RBD-specific, six NTD-specific and five S2-specfic) by ELISA- and SPR-based SVNAs identified several potential neutralizers, predominantly from the RBD-binding cohort (**Fig. S10; Table S7**). A more relevant SVNA, which assessed the ability of antibodies to block binding of S to Vero E6 cells displaying ACE2, was then used as a screen to identify neutralizing V_H_Hs and V_H_H-Fcs. Neutralizing V_H_Hs displayed similar potencies (*IC*_50_: 5 – 21 nM) and outperformed the benchmark VHH-72 (*IC*_50_: 59 nM) by as much as 12-fold (**Fig. S11; Table 2**). Compared to V_H_Hs, a larger number of V_H_H-Fcs demonstrated neutralization capabilities (**Fig. 4A; Fig. S12; Table 2**). While neutralizing monomer V_H_Hs did not benefit from reformatting (except for the VHH-72 benchmark), several non-neutralizing V_H_Hs (three RBD-specific and three NTD-specific) benefitted profoundly from reformatting and were transformed into neutralizers that had potencies similar to other RBD-specific V_H_H-Fcs. All S2-specific V_H_Hs remained non-neutralizing as V_H_H-Fcs.

**Table 2.**
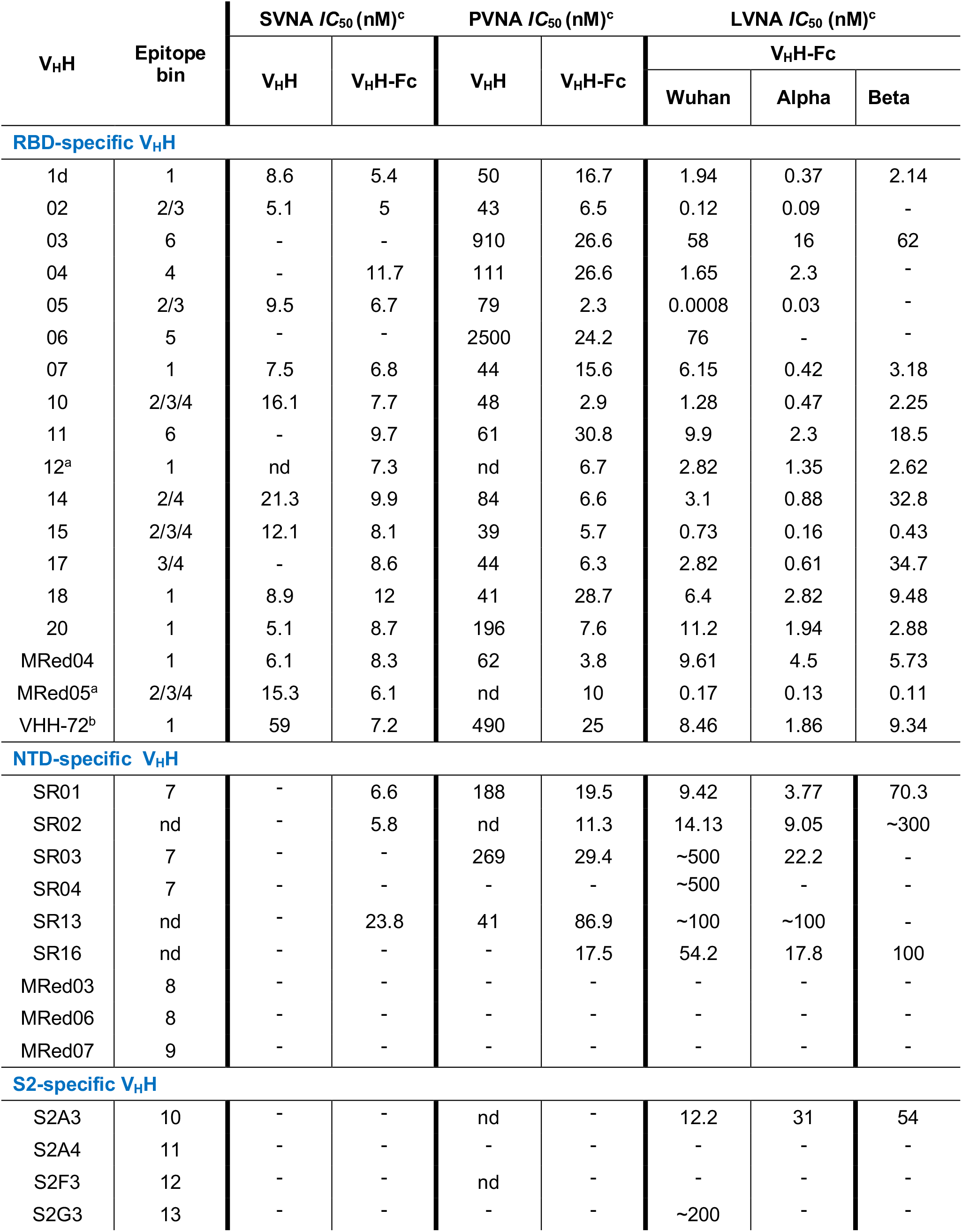
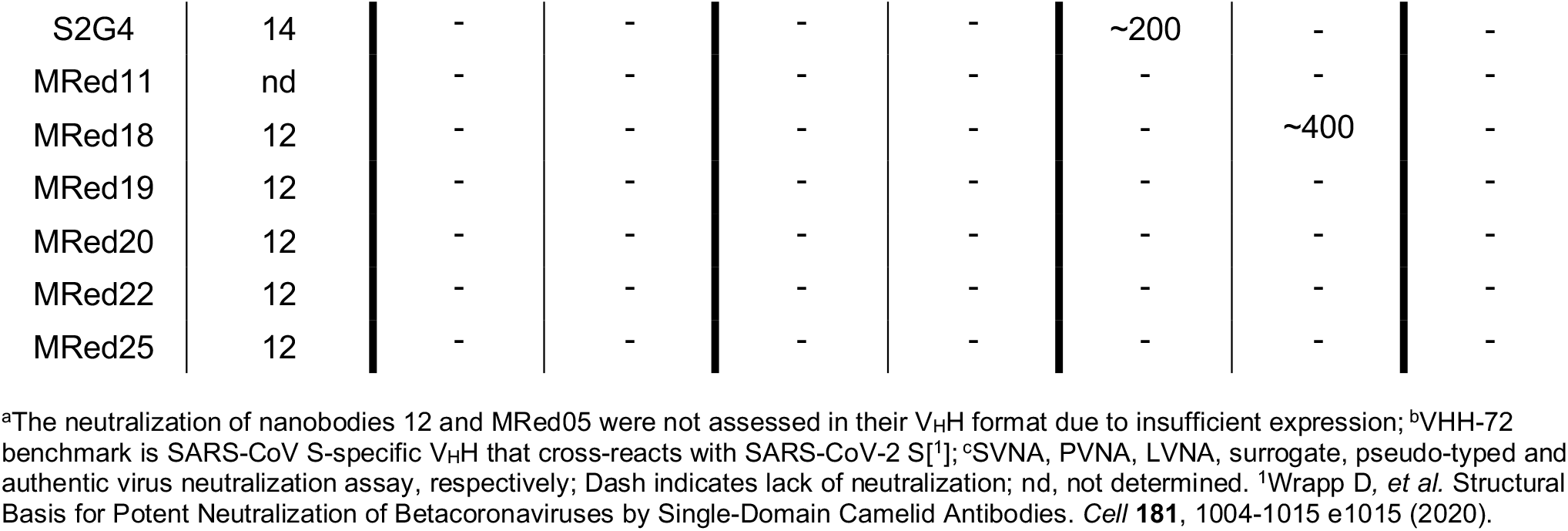
Neutralization potencies of V_H_Hs against SARS-CoV-2 (Wuhan) and variants

**Fig. 4.**
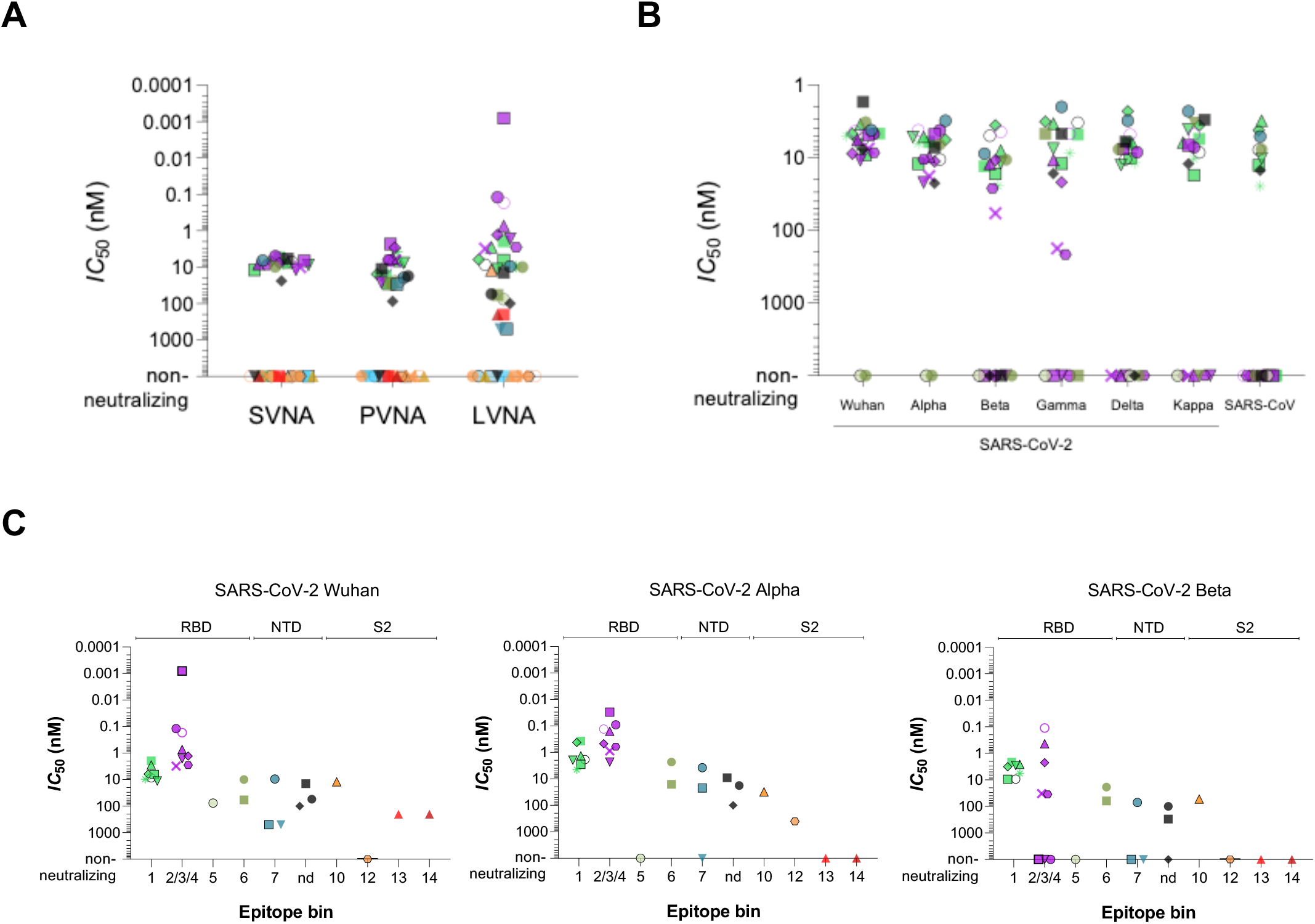
*In vitro* neutralization potency of anti-SARS-CoV-2 S V_H_H-Fcs. **A**) Summary of *IC*_50_s obtained by surrogate (SVNA), pseudotyped (PVNA) and live (LVNA) virus neutralization assays against SARS-CoV-2 Wuhan. **B**) Summary of *IC*_50_s obtained by SVNAs against SARS-CoV-2 variants Wuhan, Alpha, Beta, Gamma, Delta and Kappa and SARS-CoV. **C**) Summary of *IC*_50_s obtained by LVNAs for V_H_H-Fcs against Wuhan, Alpha and Beta SARS-CoV-2 variants. See also **Table 2** for *IC*_50_ values. Graphs were generated based on the data in **Fig. S12 – Fig. S15** and **Table S8**. Data are organized based on subunit/domain specificity and epitope bin. Data are mean ± s.e.m. of technical duplicates. Black open circle, VHH-72 benchmark. V_H_Hs are color-coded based on their epitope bin designation (*see* **Fig. 5A**).

Extending our SVNAs to variants Alpha, Beta, Gamma, Delta and Kappa using all of the RBD-specific and a subset of NTD-specific V_H_H-Fcs (**Fig. 4B; Table S8**), several observations were made. First, for cross-neutralizing V_H_Hs the *IC*_50_s across variants did not change significantly. Second, while all Wuhan neutralizers also remained Alpha neutralizers, some lost their capability to inhibit Beta, Gamma, Delta and Kappa with variable cross-neutralizing patterns. In particular, with respect to the RBD-specific V_H_Hs, the cross-neutralization profiles for Beta *vs* Gamma and Delta *vs* Kappa were identical, similarly reflective of the key escape mutations in these variants (K417N, E484K and N501Y for Beta *vs* K417T, E484K and N501Y for Gamma; L452R and T478K for Delta *vs* L452R and E484Q for Kappa). Third, and importantly, 12 out of 20 V_H_H-Fcs (10 RBD-specific, two NTD-specific) were Delta neutralizers, nine of which (eight RBD-specific, one NTD-specific) neutralized across all variants. The majority of these nine pan-neutralizers (six RBD-specific, one NTD-specific) also neutralized SARS-CoV.

### 2.6. Screening for neutralizing V_H_Hs by pseudotyped and live virus neutralization assays

Using a pseudotyped virus neutralization assay (PVNA), all 15 RBD-specific V_H_Hs tested were found to be neutralizing, and with the exception of 03 (*IC*_50_: 0.91 µM) and 06 (*IC*_50_: 2.5 µM), the V_H_Hs were potent neutralizers (*IC*_50_ range: 39 – 196 nM; median: 48 nM) (**Fig. S13A; Table 2**). For NTD-specific V_H_Hs, three of nine were neutralizing: two with similar *IC*_50_s of 188 nM (SR01) and 269 nM (SR03) and one with an *IC*_50_ of 41 nM (SR13), comparable to the most potent RBD-specific V_H_Hs. Reformatting to V_H_H-Fc had a universal enhancing effect on neutralization potencies of V_H_Hs irrespective of epitope bin origin (**Fig. 4A; Fig. S13B; Table 2**). For RBD-specific V_H_Hs, potency increases (*IC*_50_ decreases) of 2 – 100-fold were observed; only one V_H_H (18) was unaffected with reformatting (*IC*_50_ range: 2.3 - 30.8 nM; median: 7.6 nM). NTD-specific V_H_H-Fcs demonstrated weaker potencies (*IC*_50_ range: 11.3 – 86.9 nM; median: 18.5 nM; 4 of 9 non-neutralizing). However, bivalency also significantly improved (∼9-fold) the potencies of SR01 and SR03 and transformed a non-neutralizing V_H_H (SR16) into a potent neutralizing V_H_H-Fc. Consistent with the aforementioned SVNA results and previous data ^29^, the VHH-72 benchmark also improved, elevated from a weak V_H_H (*IC*_50_: 490 nM) to a strong V_H_H-Fc (25 nM) neutralizer. S2-specifiic V_H_Hs remained non-neutralizing with reformatting.

All RBD- and NTD-specific V_H_H-Fcs that were neutralizing by PVNA were also neutralizing in a live virus neutralization assay (LVNA) (**Fig. 4A; Table 2; Fig. S14**). However, compared to the former method, the LVNA *IC*_50_ values were lower and more variable. For RBD-specific V_H_H-Fcs an *IC*_50_ (range: 0.0008 – 76 nM; median: 2.8 nM) was observed. The most potent V_H_H-Fcs belonged to bin 2/3/4 (*IC*_50_ range: 0.0008 – 3.1 nM; median: 1 nM), with 05 showing the greatest potency (*IC*_50_: 0.0008 nM) followed closely by 02 and MRed05 (*IC*_50_s: 0.12 and 0.17 nM, respectively). Bin 1 neutralizers, to which VHH-72 belonged and displayed a similar *IC*_50_ (8.5 nM), exhibited intermediate potencies (range: 1.9 – 11.2 nM; median: 6.3 nM), followed by bin 5/6 neutralizers (range: 9.9 – 76 nM; median: 58 nM). Weaker neutralizing potencies were observed with NTD-specific V_H_H-Fcs. Here, six of nine V_H_H-Fcs, representing at least one epitope bin, were neutralizing. Interestingly, three new neutralizers emerged from the pool of S2- specific V_H_H-Fcs using the LVNA, with S2A3 the most potent.

The LVNAs were extended to include Alpha and Beta variants. With the exception of V_H_H-Fc 06, all remaining 16 RBD-specific Wuhan neutralizers maintained their ability to neutralize Alpha (**Table 2: Fig. 4C; Fig. S15**). Interestingly, many V_H_Hs from across different epitope bins showed improved *IC*_50_s by as high as 15-fold. Except for 05, which despite showing a reduced potency towards the Alpha variant (∼40- fold) still exhibited the highest potency of all against the variant, the remaining V_H_Hs demonstrated comparable potencies. Of the 16 Wuhan/Alpha neutralizers, 13 also neutralized the Beta variant, with the majority (10 of 13) demonstrating comparable potencies and two (14 and 17) showing reductions (∼10-fold). Although from the most potent bin (2/3/4), 02, 04 and 05, consistent with the cross-reactivity data (**Fig. 2A**), were completely abrogated presumably by the Beta mutations in the RBD (K417N, E484K, N501Y), several others including MRed05, 10 and 15 did retain their high neutralizing potencies against both Alpha and Beta variants. A similar trend was observed for the NTD-specific neutralizing V_H_Hs: against the Alpha variant, potencies either remained essentially the same as those for the Wuhan variant or improved, while against the Beta variant, potencies diminished. Nonetheless, SR01 and SR16 maintained respectable neutralization potencies against Beta. The potencies of S2-specific neutralizers (S2A3, S2G3, S2G4) were also decreased with variants. However, the lead S2A3 still maintained comparable potencies across all three variants (*IC*_50_ of 12.2 nM, 31 nM and 54 nM for Wuhan, Alpha and Beta [**Table 2**]). Collectively, the neutralization profiles across Wuhan, Alpha and Beta variants were consistent with cross-reactivity profiles (**Fig. 2A**). Based on the cross-reactivity (**Fig. 2A**) and surrogate cross-neutralization data (**Fig. 4B**; **Table S8**), it is likely that many V_H_Hs would also neutralize the Gamma, Kappa and Delta variants in LVNAs.

### 2.7. Epitope studies

To identify the number of non-overlapping epitopes, V_H_Hs were subjected to epitope binning experiments by SPR and sandwich ELISA. SPR assays were performed by injecting paired combinations of eight RBD-specific V_H_Hs, six NTD-specific V_H_Hs and ten S2-specific V_H_Hs over a SARS-CoV-2 spike glycoprotein surface (**Fig. S16A**). A conceptually similar assay to SPR was also performed by sandwich ELISA (**Fig. S16B**). From the 33 V_H_Hs tested, 14 unique epitope bins were identified: six for RBD-specific V_H_Hs, three for NTD-specific V_H_Hs and five for S2-specific V_H_Hs (**Fig. 5A; Table 1)**. The benchmark VHH-72 binned with RBD-specific V_H_Hs 1d, 07, 12, 18, 20 and MRed04. With the exception of V_H_H 04, all remaining bin 1, 2, 3 and 4 V_H_Hs (13 in total), as well as VHH-72, binned with ACE2, consistent with them being potent neutralizers. The results of sandwich ELISAs extended to include variants Alpha, Beta, Gamma, Delta and Kappa (**Fig. S16C**), using lead V_H_Hs from various bins as capture antibodies, were consistent with epitope binning as well as cross-reactivity (**Fig. 2A**) data.

**Fig. 5.**
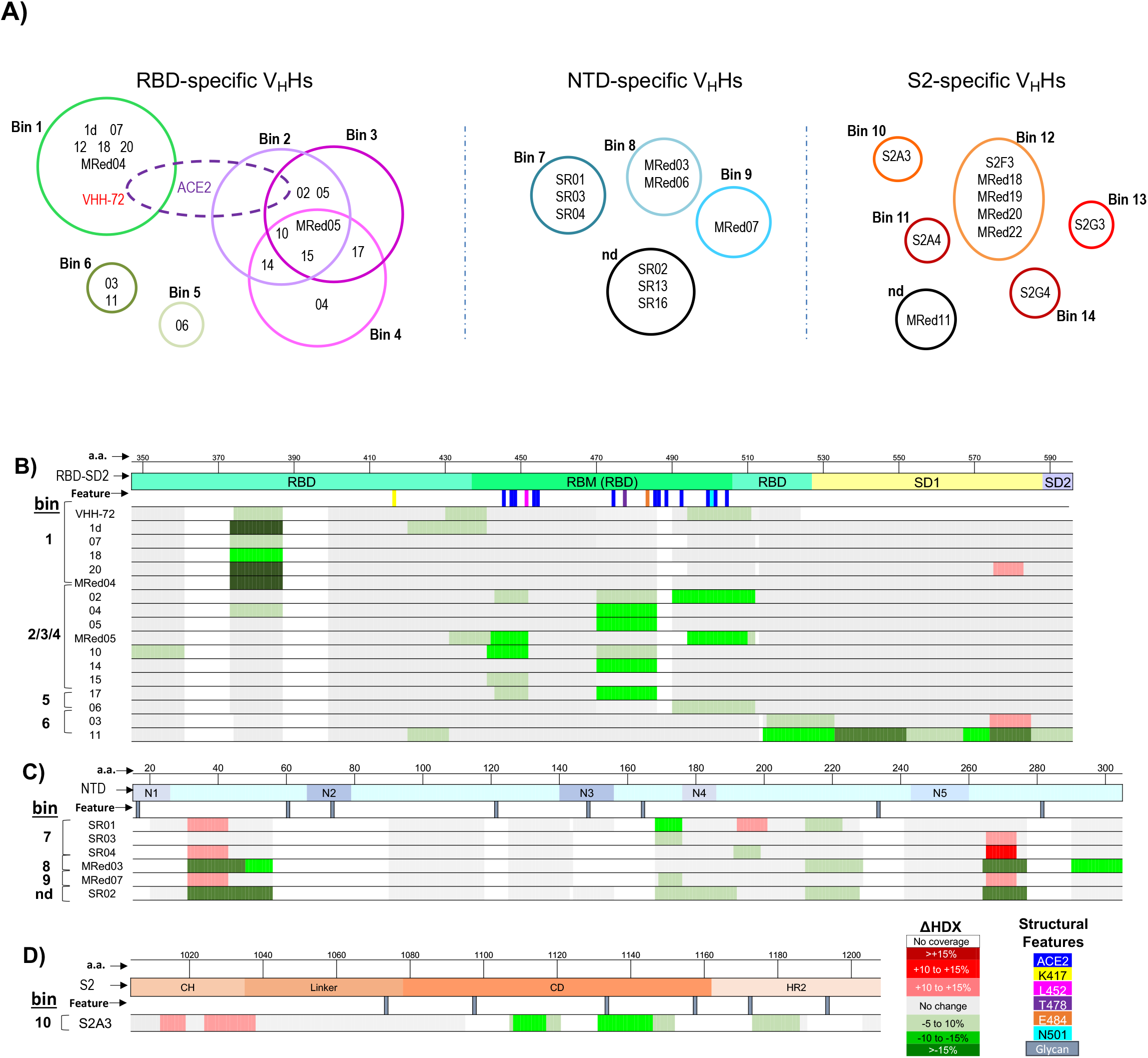
Epitope binning and mapping. **A**) Summary of epitope bins identified by SPR and ELISA (*see* **Fig. S16**). Bins are color-coded. **B – D**) HDX/MS epitope mapping of V_H_Hs binding RBD (**B**), NTD (**C**) and S2 (**D**). Changes in deuteration are mapped as colored rectangles corresponding to primary sequences. Stabilizations are shown in green, and destabilizations are shown in red, while regions with no significant changes in deuteration are shown in grey and missing coverage in white. Key structural features are highlighted by lines below the amino acid sequence, including the ACE2 binding site (blue), five mutations from VoCs, and N-linked glycans are included for reference. S2 subunit is shown partially.

Next, we investigated the conformational nature of the epitope bins at peptide-level resolution with hydrogen-exchange mass spectrometry (HDX-MS) (**Table S9**). A summary of normalized HDX shifts are shown in **Fig. 5**, and projected onto 3-D structures in **Fig. S17**. There is a strong agreement between HDX-MS profiles and previously described epitope bins and subunit/domain specificity. A common binding mode adjacent to the RBM and distant from known VoC mutations was observed for bin 1 (**Fig. 5B**). This overlaps with core binding contacts of VHH-72 on SARS-CoV ^29^ where neutralization is achieved by steric blocking of ACE2. The binding profiles for the strongest neutralizers in bins 2/3/4 overlap the ACE2 binding site ^10^ and known conformational hotspots ^60^. It was not possible to further resolve epitope diversity within the context of this dataset, however it is evident that a range of binding patterns exists ^61, 62^, and there is a correlation between stabilizations spanning mutations in VoCs and the loss/attenuation of neutralization (**Fig. 5B**). Such granularity assists in understanding and predicting neutralization potency as novel variants emerge.

Epitopes for bin 6 (V_H_H 03 and 11) span the C-terminus of the RBD and SD1 ^63^ (**Fig. 5B**), explaining why binding is limited to constructs containing SD1 (**Table S2**). Stabilization of the SD1 hinge responsible for RBD motion highlights a potential inhibitory mechanism for V_H_H 11 ^64^. While it is challenging to delineate between an epitope and conformational effects based on HDX profiles alone, distinct binding responses with common conformational hotspots were observed for the NTD binders (**Fig. 5C; Fig. S17**). Interestingly, none of the NTD-supersite loops ^14–17, 19, 65–67^ covered here displayed significant HDX shifts, except for N4 stabilized by SR02, suggesting a range of binding modes beyond the NTD-supersite. Further, stabilizations partially overlap a previously described conformationally active epitope with low variability and neutralization vulnerability ^15^. Supersite binders appear to be vulnerable to escape mutants ^7, 14, 68^, highlighting the importance of targeting and characterizing alternative NTD epitopes.

An epitope for S2A3 spanning the linker/CD/HR2 motifs ^21^ is described in **Fig. 5D**. This region is upstream of known S2 epitopes and is crucial for the structural transition required for virus-cell fusion ^69^. We cannot rule out the involvement of other residues within CD/HR2 regions due to gaps in coverage. Given that none of the mutations within the six SARS-CoV-2 variants overlap the epitope, the cross-reactivity against the six variants (**Fig. 2A**) and cross-neutralization against Alpha and Beta (**Table 2**), we predict similar neutralization potencies against the Gamma, Kappa and Delta variants.

Finally, epitope typing by denaturing SDS-PAGE and western blotting indicated 13 of the 37 V_H_Hs were recognizing linear epitopes, with the majority (9 out of 12) being S2-specific (**Table 1; Fig S18**).

### 2.8. *In vivo* therapeutic efficacy of V_H_H-Fcs

The *in vivo* therapeutic efficacy of V_H_H-Fcs which were neutralizing by LVNA were assessed in a hamster model of SARS-CoV-2 infection. Five V_H_H-Fcs were selected to cover a wide range of important attributes including *in vitro* neutralization potencies and breadth, epitope bin, subunit/domain specificity and cross-reactivity pattern. These included three RBD-specific (1d, 05, MRed05), one NTD-specific (SR01) and one S2-specific (S2A3) V_H_H-Fcs. Cocktails of two V_H_H-Fcs were also included to explore synergy between the antibody pairs recognizing distinct epitopes within the RBD (1d/MRed05) or RBD and NTD (1d/SR01).

Hamsters were administered IP with 1 mg of V_H_H-Fcs 24 h prior to intranasal challenge with SARS-CoV-2 Wuhan isolate. Daily weight change and clinical symptoms were monitored. At 5 dpi, lungs were collected to determine viral titers. Viral titer decrease and reversal of weight loss in antibody treated *versus* control animals were taken as measures of antibody efficacy. Animals treated with RBD binders 1d, 05, and MRed05 showed reduced lung viral burden by three, five and six orders of magnitude, respectively, relative to PBS or V_H_H-Fc isotype controls, with 05 and MRed05 reducing viral burden to below detectable levels (**Fig. 6A**). The RBD-specific VHH-72 benchmark caused a mean viral decrease of four orders of magnitude. The NTD binder SR01, and interestingly, the S2 binder S2A3, were also effective neutralizers, decreasing mean viral titers by four and three orders of magnitude, respectively. Both 1d/SR01 and 1d/MRed05 cocktails decreased viral titers by 6 orders of magnitude to undetectable levels of virus infection. While it was not possible to unravel potential synergies for 1d/MRed05, as MRed05 alone displayed essentially the same efficacy as the 1d/MRed05 combination, it was apparent that the 1d/SR01 combination benefited from synergy, decreasing viral titers by a further 2 - 3 orders of magnitude to undetectable levels, relative to 1d or SR01 alone. Moreover, in accordance with the viral titer decreases, a gradual reversal of weight loss in infected animals was observed with antibody treatment starting on 2 dpi (**Fig. 6B,C**). A strong negative correlation (r = -0.9436; p <0.0001) was observed between weight change and viral titer at 5 dpi (**Fig. 6D**).

**Fig. 6.**
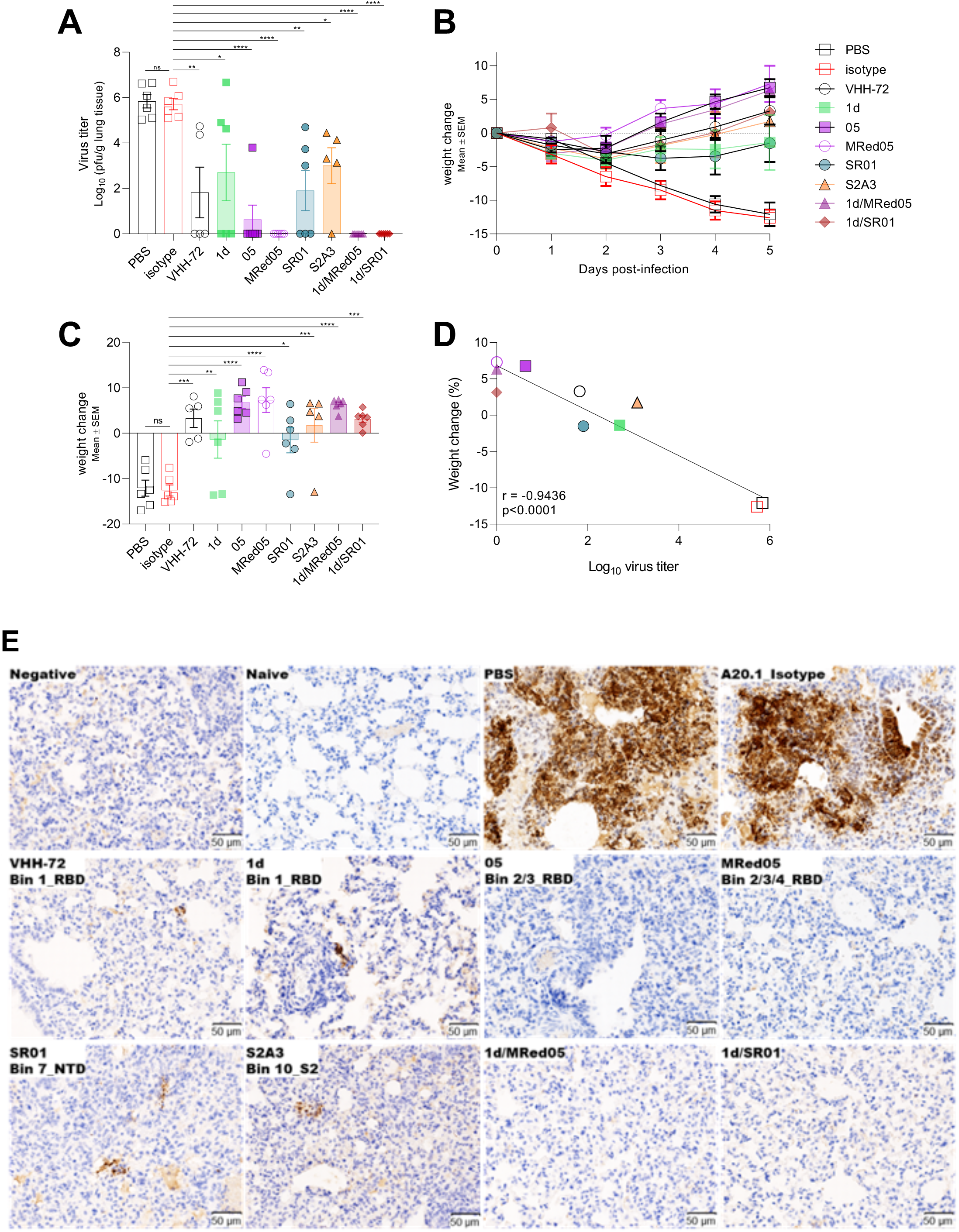
V_H_H-Fcs showed strong protective efficacy in hamsters challenged with SARS-CoV-2. **A**) Lung viral load in V_H_H-Fc-treated (VHH-72 benchmark, 1d, 05, MRed05, SR01, S2A3, 1d/MRed05, 1d/SR01) and control groups treated with PBS or isotype A20.1 V_H_H-Fc at 5 dpi. PFU, plaque-forming unit. **B**) Percent body weight change for antibody-treated and control groups. **C**) Percent body weight change at 5 dpi. In “**A**” and “**C**”, treatment effects, assessed by one-way ANOVA with Dunnett’s multiple comparison post hoc test, were significant (*p<0.05, **p<0.01, ***p<0.001 or ****p<0.0001). Dunnett’s test was performed by comparing treatment groups against the isotype control. ns, not significant. **D**) Correlation curve of body weight change *vs* viral titer at 5 dpi. A strong negative correlation (r = -0.9436, p<0.0001) between body weight change and lung viral titer was observed. **E**) Immunohistochemical demonstration of SARS-CoV-2 nucleocapsid (N) protein in the lungs of V_H_H-Fcs-treated animals. Untreated (PBS) and A20.1 isotype-treated animals showed strong viral N protein immunoreactivity which was mainly found in large multifocal patches of consolidated areas. Black arrow indicates the presence of viral N protein in bronchiolar epithelial cells. Omission of anti-nucleocapsid antibody eliminated the staining (Negative). Shown also is the absence of staining in healthy animals (Naïve). A marked reduction in viral N protein staining was seen in all lung tissues examined from V_H_H-Fc-treated animals (middle and bottom panels). While no staining was observed in 05, MRed05, 1d/SR01 and 1d/MRed05, small foci of viral N protein was detected in VHH-72, 1d, SR01 and S2A3. Representative images are shown from a single experiment.

Subsequent immunohistochemistry studies corroborated the viral titer and weight change results. First, in agreement with the viral titer observations, substantial viral antigen (nucleocapsid) reductions in hamster lungs were observed with antibody treatments (**Fig. 6E**; compare non-treated PBS and isotype controls to treated profiles). Although, small foci of viral antigen expression were detected in VHH-72-, 1d-, SR01- and S2A3-treated animals, none were detected in 05-, MRed05-, 1d/SR01- and 1d/MRed05- treated animals. Second, SARS-CoV-2 infection is characterized by an overt inflammatory response in the respiratory tract accompanied by an increased infiltration of inflammatory immune cells, e.g., macrophages and T lymphocytes, in the lung parenchyma ^70^. As expected, this was the case for the non-treated PBS and isotype control groups. In contrast, we observed a substantial reduction of macrophages and T lymphocytes infiltrate in lung parenchyma with antibody treatment (**Fig. S19, S20**). The most dramatic decreases in the number of macrophages and T lymphocytes were seen with 05, MRed05, 1d/MRed05 and 1d/SR01 treatments. Interestingly, a reduction in inflammatory responses was also associated with a decrease in the number of apoptotic cells in antibody-treated animals (**Fig. S21**). Altogether, the viral titer, weight change and immunohistochemistry results consistently demonstrate that a single dose of several of our V_H_H-Fcs reduced viral burden, immune cell infiltration and apoptosis in the lungs of infected hamsters.

## 3. DISCUSSION

The efficacy of COVID-19 therapeutic antibodies and vaccines is persistently being threatened by emergence of new VoC escape mutants, presenting a pandemic state with no end in sight. Developing broad-spectrum antibodies that can neutralize current and emerging VoCs represents an effective countermeasure against the current SARS-CoV-2 pandemic and future ones.

Two key structural features of nanobodies make them promising candidates as broad-spectrum therapeutics. First is their ability to access epitopes on the surface of the spike glycoprotein that are hidden from mAbs and conserved across VoCs ^34, 37, 45^. Second is their high modularity, allowing for their rapid conversion into multimeric/multi-paratopic constructs with favorable manufacturability profiles. Multimerization can lead to drastic increases in the efficacy of anti-COVID-19 nanobodies and broaden their cross-reactivity across variants ^29, 34, 36, 39^. Significantly, the chance of developing multimeric/multi-paratopic constructs with the desired cross-neutralization breadth improves with the diversity of nanobody building blocks available.

With the goal of developing broad-spectrum therapeutics, we employed multiple immunization, phage display library construction and panning strategies to identify a diverse collection of nanobodies. These were extensively characterized as monomeric V_H_Hs and homodimeric V_H_H-Fcs. Nanobodies were shown to have high intrinsic affinity (single-digit nM-pM); high thermal, thermodynamic and aerosolization stability; broad epitopic diversity falling into 14 different epitope clusters, broad subunit/domain specificity, recognizing NTD, RBD, and S2 regions; broad cross-reactivities recognizing up to 11 different Sarbecoviruses including several SARS-CoV-2 VoCs and high and broad *in vitro* and *in vivo* virus neutralization potencies/efficacies. Based on published results and our studies, a diversity of neutralization mechanisms of action can be envisaged for the anti-spike glycoprotein V_H_Hs, including inhibiting the ACE2-RBD interaction by direct competition, steric hindrance, locking the RBDs in the closed conformation and distorting the RBM, leading to inhibition of the virus-cell binding (RBD- and NTD-specific V_H_Hs) ^38, 71–73^ and inhibiting conformational rearrangements leading to inhibition of virus-cell fusion (NTD- and S2-specific V_H_Hs)^14, 16, 17, 74^.

Our study also provides valuable insights into how various *in vitro* neutralization assays predict antibody efficacy. The flow cytometry-based SVNA was shown to effectively identify neutralizing antibodies that were RBD-specific, providing a viable alternative to the PVNA or LVNA, which are labor-intensive, difficult to standardize, inconvenient and not readily accessible as they require operating in biosafety level 2 or 3 labs. However, the SVNA occasionally missed NTD-specific neutralizing antibodies, and, similar to the PVNA, failed altogether to identify neutralizing antibodies that were S2-specific. Furthermore, the SVNA did not have the sensitivity of the LVNA to identify V_H_Hs with weaker potencies or, similar to the PVNA, to fine-rank neutralizing antibodies.

For several reasons, the number of epitope bins (14) identified in the current study likely under- estimates the number of actual distinct epitopes. First, V_H_Hs that recognize (i) partially overlapping epitopes, (ii) fully overlapping epitopes of significantly different nature, or (iii) non-overlapping epitopes, but manifest exclusive binding as a consequence of conformational competition or steric clashes between the V_H_H pairs, would fall under the same epitope bins. Second, indicators of distinct epitopes such as differential HDX-MS footprints, epitope types, cross-reactivity profiles, neutralization potencies, and cross-neutralization profiles not accounted for by affinity alone, are seen amongst V_H_Hs within the same bin. Thus, the repertoire of structurally and functionally distinct epitopes are more diverse than what can be gleaned from epitope binning analysis alone.

*In vitro* neutralization assays with the Wuhan SARS-CoV-2 variant showed that the majority of the nanobodies were RBD-specific. Importantly, and to our knowledge for the first time, we demonstrated that several NTD- and S2-specific V_H_Hs were also potent and efficacious neutralizers *in vitro* and *in vivo*. Significantly, neutralizing nanobodies showed high epitopic diversity, originating from at least eight different epitope clusters. The vast majority of these V_H_Hs - including NTD and S2 V_H_Hs - remained potent *in vitro* neutralizers against the Alpha, Beta, Gamma, Delta and Kappa variants as well. A sample of *in vitro* neutralizing nanobodies, representing RBD-, NTD- and S2-specific V_H_Hs, were also shown to be efficacious *in vivo* neutralizers as single V_H_H-Fcs or as paired combinations of V_H_H-Fcs that targeted RBD and NTD, with some capable of complete viral clearance from hamster lungs. These results also confirm the strong positive correlation between the *in vitro* and *in vivo* neutralization data and indicate that the remaining *in vitro* neutralizers should also be *in vivo* neutralizers. Moreover, given the broad cross-reactivity and cross-neutralization profiles of many of these nanobodies, it is reasonable to expect these pan-reactive antibodies will also effectively neutralize currently untested and emerging VoCs, e.g., Omicron, and SARS-related viruses.

The VHH-72 benchmark nanobody, a modified, enhanced V_H_H-Fc version of which is currently being developed for COVID-19 therapy, has the advantage of being broadly neutralizing and binding to a highly conserved cryptic epitope in the RBD region which is difficult to access with conventional mAbs ^29, 39^. Here we have identified five VHH-72-like nanobodies (1d, 07, 12, 20, MRed04) that map to the same epitope as VHH-72 and demonstrate similar cross-reactivity and cross-neutralization profiles in the V_H_H- Fc format. In monomeric formats, however, the V_H_Hs significantly outperform VHH-72, but whether they would do the same against the enhanced version of VHH-72 remains to be seen. A similar broad cross-reactivity profile to VHH-72 was seen with two other neutralizing nanobodies (11, SR01) that by HDX/MS experiments map to different but well-conserved epitopes, indicating that these nanobodies may similarly bind to cryptic epitopes conserved across variants. The doubly pan-specific SR01/1d cocktail completely cleared hamsters of viral burden, making for a promising broad-spectrum combination therapeutic.

With an abundance of neutralizing nanobodies on hand, many possibilities exist for designing optimized multimeric/multi-paratopic therapeutic agents. This is implicitly evident from our epitope binning experiments performed against several SARS-CoV-2 variants (**Fig. S16C**), albeit they exemplified a fraction of potential possibilities. A recent study showed that the neutralization capability of a nanobody significantly increased when fused to a second, non-neutralizing nanobody in a bi-paratopic format ^43, 74^. Thus, the pool of V_H_Hs can be expanded to include the entire panel of neutralizing and non-neutralizing nanobodies in the current study, leading to a further exponential increase of multimer possibilities. Including the same or similar (same epitope bin) nanobodies in a multimer construct is plausible given the trimeric nature of the spike glycoprotein and its repetitive presentation on the surface of the virus which should accommodate avid inter-protomer, intra-spike, and/or inter-spike binding events. This is supported by the current data showing increased *in vitro* neutralization potency of nanobodies with homodimerization. CryoEM and X-ray crystallography structures of nanobody-spike glycoprotein complexes would reveal the relative positioning of nanobodies on the surface of the spike glycoprotein and could be used as a guide for designing highly effective and broad-spectrum therapeutic multimers. A comprehensive, high through-put campaign involving small scale expression of multimers followed by *in vitro* screening for broad neutralization can also be envisaged.

In summary, the nanobodies described here provide ample opportunities to develop broad-spectrum COVID-19 prophylactic/therapeutics of good manufacturability and efficacy. They can be complemented with nanobodies described in the literature for specified performance, although such alternatives would be lacking in the case of our unprecedented NTD- and S2-specific nanobodies. The ability to aerosolize our V_H_Hs provides the option of a cost-effective, patient-friendly, direct and effective delivery of therapeutic nanobodies to the nasal and lung epithelia by inhalation ^40, 41, 50–52, 75^. The nanobodies can additionally be utilized to develop cost-effective, highly sensitive detection/ diagnosis systems of desired cross-reactivity specifications. Epitope type diversity further provides the flexibility of virus detection/diagnosis under native and/or denaturing conditions.

## 4. MATERIALS AND METHODS

### 4.1. Recombinant antigens and ACE2

Purified recombinant spike and ACE2 proteins used in the current study are described in **Table S1**. They were either purchased or produced in-house as described (**Table S1**) (^76–80^). Proteins were purified using standard immobilized metal-ion affinity chromatography or protein A affinity chromatography.

### 4.2. Antigen validation

#### a) Binding to cognate human angiotensin converting enzyme (ACE2) receptor

ELISA was performed to determine if spike glycoprotein fragments (Wuhan) were able to bind to human ACE2 when passively adsorbed (S, S1, RBD and S2) or directionally captured (S1, RBD) on microtiter wells. For passive adsorption, wells of NUNC® Immulon 4 HBX microtiter plates (Thermo Fisher, Ottawa, Canada, Cat#3855) were coated with 50 ng of SARS-CoV-2 spike proteins (S, S1, S2, RBD) in 100 µL of phosphate-buffered saline (PBS) overnight at 4°C. Following removal of protein solutions and three washes with PBST (PBS supplemented with 0.05% [v/v] Tween 20), wells were blocked with PBSC (1% [w/v] casein [Sigma, Oakville, Canada, Cat#E3414] in PBS) at room temperature for 1 h. For capturing, *in vivo* biotinylated fragments harboring the AviTag^TM^ (AviTag-S1, AviTag-RBD) were diluted in PBS and added at 50 ng/well (100 µL) to pre-blocked Streptavidin Coated High Capacity Strip wells (Thermo Fisher, Cat#15501). After 1 h incubation at room temperature, wells were washed five times with PBST and incubated for an additional hour with 100 µL/well of 2-fold serially diluted ACE2-Fc (human ACE2 fused to human IgG1 Fc; ACROBiosystems, Newark, DE, Cat#AC2-H5257) in PBSTC (PBS/0.2% casein/0.1% Tween 20). Wells were washed five times and incubated for 1 h with 1 µg/mL HRP-conjugated goat anti-human IgG (Sigma, Cat#A0170). Finally, wells were washed 10 times and incubated with 100 µL peroxidase substrate solution (SeraCare, Milford, M, Cat#50-76-00) at room temperature for 15 min. Reactions were stopped by adding 50 µL 1 M H_2_SO_4_ to wells, and absorbance were subsequently measured at 450 nm using a Multiskan™ FC photometer (Thermo Fisher).

#### b) Binding to cognate anti-spike glycoprotein polyclonal antibody

The four spike glycoprotein antigens were passively adsorbed as described above. After blocking with PBSC, wells were emptied, washed five times with PBST and incubated at room temperature for 1 h with 100 µL of 1 µg/mL anti-SARS-CoV-2 spike rabbit polyclonal antibody (Sino Biological, Beijing, China, Cat#40589-T62) in PBSCT. Following 10 washes with PBST, wells were incubated with 100 µL 1/2500 dilution (320 ng/mL) of goat anti-rabbit:HRP (Jackson ImmunoResearch, West Grove, PA, Cat#111-035-144) in PBSCT for 1 h at room temperature. After 1 h incubation and final five washes with PBST, the peroxidase activity was determined as described above.

### 4.3. Llama immunization and serum analyses

#### a) Llama immunization

Immunizations were performed at Cedarlane Laboratories (Burlington, Canada) essentially as described ^81, 82^, using SARS-CoV-2 Wuhan spike glycoprotein fragments. Briefly, for priming, both llamas (Green and Red) were injected with 100 µg of S in 500 µL PBS combined with 500 µL of Freund’s Complete Adjuvant. For the three subsequent boosts (days 7, 14, 21), Green was injected with 70 µg of RBD (ACROBiosystems, Cat#SPD-S52H6) whereas Red received 100 µg of S ^78^ at day 7 and 50 µg of S on days 14 and 21 all with Freund’s Incomplete Adjuvant. Experiments involving animals were conducted using protocols approved by the National Research Council Canada Animal Care Committee and in accordance with the guidelines set out in the OMAFRA Animals for Research Act, R.S.O. 1990, c. A.22.

#### b) Serum ELISA

Llama sera were tested for antigen-specific immune response by ELISA essentially as described ^82, 83^. Briefly, dilutions of sera in PBST were added to wells pre-coated with S, S1, S2 or RBD. Negative antigen control wells were pre-coated with casein (100 µL of 1%) or dipeptidase 1, DPEP1 (50 ng/well; Sino Biological, Cat#13543-H08H). Following 1 h incubation at room temperature, wells were washed 10 times with PBST and incubated with HRP-conjugated polyclonal goat anti-llama IgG heavy and light chain antibody (Bethyl Laboratories, Montgomery, TX, Cat#A160-100P) for 1 h at room temperature. After 10 washes, the peroxidase activity was determined as described in **4.2*a***.

#### c) Serum surrogate neutralization assay by flow cytometry

SARS-CoV-2 S was chemically biotinylated using EZ-Link™ NHS-LC-LC-Biotin following manufacturer instructions (Thermo Fisher, Cat#21343). Vero E6 cells (ATCC, Cat#CRL-1586) were maintained according to ATCC protocols. Briefly, cells were grown to confluency in DMEM medium (Thermo Fisher, Cat#11965084) supplemented with 10% (w/v) heat inactivated fetal bovine serum (FBS; (Thermo Fisher, Cat#10438034) and 2 mM Glutamax (Thermo Fisher, Cat#35050061) at 37°C in a humidified 5% CO_2_ atmosphere in T75 flasks. For flow cytometry experiments, cells were harvested by Accutase (Thermo Fisher, Cat#A1110501) treatment, washed once by centrifugation with PBS, and resuspended at 1 × 10^6^ cells/mL in PBSB (PBS containing 1% [w/v] BSA and 0.05% [v/v] sodium azide [Sigma, Cat#S2002]). Cells were kept on ice until use. To determine the presence of antibodies that block the binding of S to ACE2 (surrogate for neutralization) in the immune sera of llamas, 400 ng of chemically biotinylated SARS-CoV-2 S was mixed with 1 × 10^5^ Vero E6 cells in the presence of 2-fold dilutions of sera (pre immune, day 21 and day 28 sera) in a final volume of 150 µL. Following 1 h of incubation on ice, cells were washed twice with PBSB by centrifugation for 5 min at 1200 rpm and then incubated for an additional hour with 50 µL of Streptavidin, R-Phycoerythrin Conjugate (SAPE, Thermo Fisher, Cat#S866) at 250 ng/mL diluted in PBSB. After a final wash, cells were resuspended in 100 µL PBSB and data were acquired on a CytoFLEX S flow cytometer (Beckman Coulter, Brea, CA) and analyzed by FlowJo software (FlowJo LLC, v10.6.2, Ashland, OR). Percent inhibition (neutralization) was calculated according to the following formula:

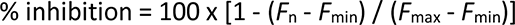

Where,

*F*_n_ is the measured fluorescence at any given competitor serum dilution

*F*_min_ is the background fluorescence measured in the presence of cells and SAPE only

*F*_max_ is the maximum fluorescence measured in the absence of competitor serum

### 4.4. Phage display library construction, selection and screening

#### a) Phage display library construction

On day 28, 100 mL of blood from each of the two llamas was drawn and peripheral blood mononuclear cells (PBMCs) were purified by ficoll gradient at Cedarlane Laboratories. Two independent phage-displayed V_H_/V_H_H libraries were constructed from ∼5 × 10^7^ PBMCs as described previously ^81, 82, 84^. Briefly, total RNA was extracted from PBMCs using TRIzol™ Plus RNA Purification Kit (Thermo Fisher, Cat#12183555) following the manufacturer’s instructions and used to reverse transcribe cDNA with SuperScript™ IV VILO™ Master Mix supplemented with random hexamer (Thermo Fisher, Cat#SO142) and oligo (dT) (Thermo Fisher, Cat#AM5730G) primers. V_H_/V_H_H genes were amplified using semi-nested PCR and cloned into the phagemid vector pMED1, followed by transformation of *E. coli* TG1 (Lucigen, Middleton, WI, Cat#60502-02) to construct two libraries with sizes of 1 × 10^7^ and 2 × 10^7^ independent transformants for Green and Red, respectively. Both libraries showed an insert rate of ∼95% as verified by DNA sequencing. Phage particles displaying the V_H_Hs were rescued from *E. coli* cell libraries using M13K07 helper phage (New England Biolabs, Whitby, Canada, Cat#N0315S) as described in ^81^ and used for selection experiments described below.

#### b) Library selection and screening

Library panning and screening were performed essentially as described ^81, 82, 85^, using SARS-CoV-2 Wuhan spike glycoprotein fragments as target antigens. Library selections were performed on microtiter wells under six different phage binding/elution conditions designated P1 – P6. Briefly, for the phage binding step, library phages were diluted at 1 × 10^11^ colony- forming units (CFU)/mL in PBSBT [PBS supplemented with 1% BSA and 0.05% Tween 20] and incubated in antigen-coated microtiter wells for 2 h at 4°C. For P1 – P4, phages were added to wells with passively- adsorbed S (10 µg/well; P1), passively-adsorbed S2 (10 µg/well; P2), streptavidin-captured biotinylated S1 (0.5 µg/well; P3) and streptavidin-captured biotinylated RBD (0.5 µg/well; P4). For P5, phages were pre-absorbed on passively-adsorbed RBD wells (10 µg/well) for 1 h at 4°C and then the unbound phage in the solution was transferred to wells with streptavidin-captured biotinylated S1 (0.5 µg/well) in the presence of non-biotinylated RBD competitor in solution (10 µg/well). Following the binding stage (P1 – P5), wells were washed 10 times with PBST and bound phages were eluted by treatment with 100 mM glycine, pH 2.2, for 10 min at room temperature, followed by immediate neutralization of phages with 2 M Tris. Similar to P4, in P6, phages were bound on streptavidin-captured biotinylated RBD but elution of bound phages were carried out competitively with 50 nM human ACE2-Fc following the washing step. For all pannings, a small aliquot of eluted phage was used to determine the titer on LB-agar/ampicillin plates and the remaining phage were used for subsequent amplification in *E.coli* TG1 strain ^81^. The amplified phages were used as input for the next round of selection as described above.

After two rounds of selection, 16 (Green) or 12 (Red) colonies from each of the P1 – P6 selections were screened for antigen binding by monoclonal phage ELISA against S, S1, S2 and RBD. Briefly, individual colonies from eluted-phage titer plates were grown in 96 deep well plates in 0.5 mL 2YT media/100 µg/mL-carbenicillin/1% (w/v) glucose at 37°C and 250 rpm to an OD_600_ of 0.5. Then, 10^10^ CFU M13K07 helper phage was added to each well and incubation continued for another 30 min under the same conditions. Cells were subsequently pelleted by centrifugation, the supernatant was discarded and the bacterial pellets were resuspended in 500 µL 2YT/100 µg/mL carbenicillin/50 µg/mL kanamycin and incubated overnight at 28°C. Next day, phage supernatants were recovered by centrifugation, diluted 3- fold in PBSTC and used in subsequent screening assays by ELISA. To this end, antigens were coated onto microtiter wells at 50 ng/well overnight at 4°C. Next day, plates were blocked with PBSC, washed five times with PBSTC, and 100 µL of phage supernatants prepared above were added to wells, followed by incubation for 1 h at room temperature in an orbital shaking platform. After 10 washes, binding of phages was detected by adding 100 µL/well of anti-M13:HRP (Santa Cruz Biotechnology, Santa Cruz, CA, Cat#SC-53004HRP) at 40 ng/mL in PBSTC and incubating as above. After 10 washes, the peroxidase activity was determined as described in **4.2*a***. After monoclonal phage ELISA confirmed the library panning was successful,, a total of ≈1200 clones (≈100 clones per panning strategy; ≈600 clones per library) were subjected to colony-PCR and DNA sequencing, resulting in the identification of 26 (Green) and 11 (Red) V_H_Hs.

### 4.5. Expression and purification of V_H_Hs and V_H_H-Fcs

#### a) Expression and validation of V_H_Hs

Positive V_H_Hs were cloned into a modified pET expression vector (pMRo.BAP.H6) for their production in BL21(DE3) *E.coli* as monomeric soluble protein ^84^. For the VHH-72 benchmark ^29^, the sequence of the V_H_H was synthesized as a GeneBlock (Integrated DNA Technologies, Coralville, IA) flanked by SfiI sites for cloning into pMRo.BAP.H6. Briefly, individual colonies were cultured overnight in 10 mL LB supplemented with 50 µg/mL of kanamycin (LB/Kan) at 37°C and 250 rpm. After 16 h, cultures were added to 250 mL LB/Kan and grown to an OD_600_ of 0.6. Expression of V_H_Hs was induced with 10 µM of IPTG (isopropyl β-D-1-thiogalactopyranoside) overnight at 28°C and 250 rpm. Next day, bacterial pellets were harvested by centrifugation at 6,000 rpm for 15 min at 4°C, V_H_Hs were extracted by sonication and purified by IMAC as described ^84^. Protein purity was evaluated by SDS-PAGE using 4–20% Mini-PROTEAN® TGX Stain-Free™ Gels (BioRad, Hercules, CA, Cat#17000435). In addition, for ELISA (*see* below), V_H_Hs were enzymatically biotinylated in their BAP tag by incubating 1 mg of purified V_H_Hs with 10 µM of ATP (Alfa Aesar, Haverhill, MA, Cat#CAAAJ61125-09), 100 µM of D-(+)-biotin (VWR, Mississauga, Canada; Cat#97061-446) and bacterial cell extract overexpressing *E.coli* BirA as described ^81^. V_H_Hs were validated for binding by soluble ELISA against spike glycoprotein fragments (S, S1, RBD, NTD, S2). Briefly, microtiter well plates were coated with 50 ng/well SARS-CoV-2 spike glycoprotein fragments in 100 µL PBS overnight at 4°C. Plates were blocked with PBSC for 1 h at room temperature, then washed five times with PBST and incubated with increasing concentrations of biotinylated V_H_Hs. After 1 h incubation, plates were washed 10 times with PBST and binding of V_H_Hs was probed using HRP-streptavidin (Jackson ImmunoResearch, Cat#016-030-084). Finally, plates were washed 10 times with PBST and peroxidase activity was determined as described in **4.2*a***.

#### b) Production of V_H_Hs in mammalian cells in fusion with human IgG1 Fc (V_H_H-Fcs)

Codon-optimized genes for bivalent V_H_H-Fcs were synthesized and cloned into pTT5 (GenScript; Piscataway, NJ. For VHH- 72 V_H_H-Fc, ^29^ the sequence of the V_H_H was synthetized as GeneBlock (Integrated DNA Technologies) flanked by NarI/HindIII for cloning into pTT5. V_H_H-Fcs were produced by transient transfection of HEK293-6E cells followed by protein A affinity chromatography as previously described ^84^. Proteins were buffer exchanged using Amicon® Ultra-15 Centrifugal Filter Units (Millipore-Sigma, Oakville, Canada, Cat#UFC905024) with PBS, pH 7.4. Protein purity was evaluated by SDS-PAGE using 4–20% Mini-PROTEAN® TGX Stain-Free™ Gels (BioRad; Cat#17000435).

### 4.6. Affinity and specificity assays

#### a) Cross-reactivity assays by ELISA

Recombinant coronavirus spike glycoproteins S (**Table S1**) were coated overnight onto NUNC® Immulon 4 HBX microtiter plates (Thermo Fisher) at 50 ng/well in 100 µL of PBS, pH 7.4. The next day, plates were blocked with 200 µL PBSC for 1 h at room temperature, then washed five times with PBST and incubated at room temperature for 1 h on rocking platform at 80 rpm with 1 µg/mL V_H_H-Fc diluted in PBSTC. Plates were washed five times with PBSTC and binding of V_H_H-Fcs was detected using 1 µg/mL HRP-conjugated goat anti-human IgG. Finally plates were washed five times and peroxidase (HRP) activity was measured as described in **4.2*a***.

#### b) Affinity/specificity determination of V_H_Hs against SARS-CoV spike (S), SARS-CoV-2 spike (S) and SARS-CoV-2 spike fragments by surface plasmon resonance (SPR)

Standard SPR techniques were used for binding studies. All SPR assays were performed on a Biacore T200 instrument (Cytiva, Vancouver, Canada) at 25°C with HBS-EP running buffer (10 mM HEPES, 150 mM NaCl, 3 mM EDTA, 0.005% Tween 20, pH 7.4) and CM5 sensor chips (Cytiva). Prior to SPR analyses all analytes in flow (V_H_Hs, ACE2 receptor) were purified by size exclusion chromatography (SEC) on a Superdex 75™ Increase 10/300 GL column (Cytiva) in HBS-EP buffer at a flow rate of 0.8 mL/min to obtain monomeric proteins. SARS-CoV spike glycoprotein (S), SARS-CoV-2 spike glycoprotein (S)^78^ and various SARS-CoV-2 spike glycoprotein fragments were immobilized on CM5 sensor chips through standard amine coupling (10 mM acetate buffer, pH 4.0, Cytiva). On the first sensor chip, 1983 response units (RUs) of SARS-CoV spike (SinoBiological, Cat#40634-V08B), 843 RUs of SARS-CoV-2 RBD/SD1 fused to human Fc (RBD/SD1-Fc) and 972 RUs of EGFR (Genscript, Piscataway, NJ, Cat# Z03194, as an irrelevant control surface) were immobilized. On a second sensor chip, 2346 RUs of SARS-CoV-2 S, 1141 RUs of SARS-CoV-2 S1 subunit and 1028 RUs of SARS-CoV-2 S2 subunit were immobilized. A third sensor chip contained 489 RUs of RBD_short^78^. The theoretical maximum binding response for V_H_Hs binding to these surfaces ranged from 224 – 262 RUs. An ethanolamine blocked surface on each sensor chip served as a reference. Single cycle kinetics was used to determine V_H_H and ACE2 binding kinetics and affinities. V_H_Hs at various concentration ranges (from 0.25 – 4 nM to 125 – 2000 nM) were flowed over all surfaces at a flow rate of 40 µL/min with 180 s of contact time and 600 s of dissociation time. Surfaces were regenerated with a 12 s pulse of 10 mM glycine, pH 1.5, at a flow rate of 100 µL/min. Injection of EGFR-specific V_H_H NRCsdAb022^84^ served as a negative control for the SARS-CoV and SARS-CoV-2 surfaces and as a positive control for the EGFR surface. The ACE2 affinity was determined using similar conditions by flowing a range of monomeric ACE2 concentrations (31.3 – 500 nM). SARS-CoV-2 spike glycoproteins from Alpha and Beta variants were also tested by SPR and amine coupled using the conditions described above. All affinities were calculated by fitting reference flow cell-subtracted data to a 1:1 interaction model using BIAevaluation Software v3.0 (Cytiva).

For V_H_H 12 and MRed05, V_H_H-Fc formats were used in SPR experiments. Approximately 200 RUs of V_H_H-Fcs (2 µg/mL) were captured on goat anti-human IgG surfaces (4000 RUs, Jackson ImmunoResearch, Cat#109-005-098) at a flow rate of 10 µL/min for 30 s. A range of SEC-purified RBD fragments (**Table S1;** Wuhan^78^, Alpha and Beta) at 0.62 – 10 nM were flowed over the captured V_H_H-Fc at a flow rate of 40 µL/min with 180 s of contact time and 300 s of dissociation. Surfaces were regenerated with a 120 s pulse of 10 mM glycine, pH 1.5, at a flow rate of 50 µL/min. Affinities were calculated from reference flow cell subtracted sensorgrams as described above.

#### c) Domain specificity determination of V_H_Hs by ELISA

V_H_Hs which bound to S1 subunit but not to the RBD domain in SPR assays were further examined by ELISA to determine if they were binding to the NTD domain of S1. Briefly, S, S1, NTD and RBD were coated onto NUNC® Immulon 4 HBX microtiter plates at 100 ng/well in 100 µL PBS, pH 7.4. Next day, plates were blocked with 200 µL PBSC for 1 h at room temperature, then washed five times with PBST and incubated with fixed (13 nM) or decreasing concentrations of V_H_H-Fcs diluted in PBSTC. After 1 h, plates were washed 10 times with PBSTC and binding of V_H_H-Fc fusions was detected by incubating wells with 100 µL of 1 µg/mL HRP-conjugated goat anti-human IgG. Finally, plates were washed 10 times with PBST and peroxidase activity was determined as described in **4.2*a***. *EC*_50_s for the binding of V_H_H-Fcs to S and S fragments were obtained from the plot of A_450 nm_ (binding) *vs* V_H_H-Fc concentration.

#### d) Cell binding assays by flow cytometry

CHO^55E1™^ cells expressing full-length (including transmembrane and C-terminal domains) SARS-CoV-2 Wuhan S (CHO-SPK) under control of the cumate-inducible CR5 promoter were generated by methionine sulfoximine (MSX) selection of plasmid-transfected cells, as described ^86^. Cells were grown in BalanCD™ CHO Growth A medium (Irvine Scientific, Santa Ana, CA) supplemented with 50 µM methionine sulfoximine (MSX) at 120 rpm and 37°C in a humidified 5% CO_2_ atmosphere. Expression of S was induced by adding cumate at 2 µg/mL for 48 h at 32°C. For flow cytometry experiments, cells were harvested by centrifugation and resuspended at 1 x 10^6^ cells/mL in PBSB. Cells were kept on ice until use. Serially, three-fold dilutions of V_H_H-Fcs were prepared in V-Bottom 96-well microtiter plates (Globe Scientific, Mahwah, NJ, Cat# 120130) and mixed with 50 µL of CHO-SPK cells. Plates were incubated for 1 h on ice, washed twice with PBSB by centrifugation for 5 min at 1200 rpm and then incubated for an additional hour with 50 µL of R-Phycoerythrin AffiniPure F(ab’)₂ Fragment Goat Anti-Human IgG (Jackson ImmunoResearch, Cat#109-116-170) at 250 ng/mL diluted in PBSB. After a final wash, cells were resuspended in 100 µL PBSB and data were acquired on a Beckman Coulter CytoFlex S and analyzed by FlowJo™ (FlowJo LLC, v10.6.2). *EC*_50_s for the binding of V_H_H-Fcs to CHO-SPK cells were obtained from the plot of MFI (Mean Fluorescent Intensity) *vs* V_H_H-Fc concentration.

### 4.7. Stability assays

#### a) Determination of aggregation resistance by SEC

Purified V_H_Hs were subjected to SEC to validate their aggregation resistance. Briefly, 2 mg of each affinity purified V_H_H was injected into Superdex™ 75 10/300 GL column (Cytiva) connected to an ÄKTA FPLC protein purification system (Cytiva) as previously described ^87^. PBS was used as the running buffer at 0.8 mL/min. Fractions corresponding to the monomeric peak were pooled and stored at 4°C until use.

#### b) Thermostability determinations by circular dichroism

To determine thermostability, V_H_H T_m_s were measured by circular dichroism as previously described ^87^. Ellipticity of V_H_Hs were determined at 200 µg/mL V_H_H concentrations and 205 nm wavelength in 100 mM sodium phosphate buffer, pH 7.4. Ellipticity measurements were normalized to percentage scale and *T*_m_s were determined from plot of % folded *vs* temperature and fitting the data to a Boltzmann distribution.

#### c) Isothermal chemical denaturation (ICD)

All ICD experiments were performed with the automated Hunky system (Unchained Labs, Pleasanton, CA), using Hunky Client software (v1.2). V_H_Hs were prepared in PBS buffer, pH 7.2 (Teknova, Hollister, CA, Cat#P0191), and diluted by Hunky automation to 20 µg/mL. They were denatured using a linear dilution gradient of 0 - 5.52 M guanidine·HCl (Sigma, Cat#G3272) and incubated for 2 h at 25°C. Samples were subjected to LED excitation at 280 nm, and emission spectra of the V_H_H were captured by a CCD spectrometer from 300 - 720 nm. The Hunky Analysis v1.2 Software automatically plotted the fluorescence intensity against denaturant concentration, and generated a data fit curve for two-state transitions of each V_H_H. All samples were analyzed by the Hunky software using the Barycentric Mean (BCM) except for 05 (348/342 nm ratio), 06 (single 348 nm) and 07 (wavelength diff. 348-420 nm) to determine Δ*G*^0^ (kJ/mol), *C*_m_ (M) and *m* (kJ/M*mol). The fraction denatured was automatically plotted against denaturant concentration to confirm the two-state model. The denatured-induced unfolding of V_H_Hs was considered to be reversible based on their small size as shown to be the case for small proteins and numerous times for V_H_Hs ^56–59, 88, 89^.

#### d) Serum stability

Three female Syrian hamsters were injected intraperitoneally (IP) with 1 mg of 1d or VHH-72 V_H_H-Fc diluted in 200 µL PBS. Animal were bled on day 0, 1 and 4 post injection and their sera were subjected to ELISA to detect their V_H_H-Fc levels. Briefly, sera were diluted 1/6000, in order to give final A_450nm_ readings of ≤ 1 in 15 min, and incubated 1 h at room temperature in wells coated with SARS-CoV-2 Wuhan S ^78^. The binding of V_H_H-Fcs within the sera was detected using 1 µg/mL goat anti-human IgG Fc, HRP-conjugated (Thermo Fisher, Cat#A18829). Plates were washed five times with PBST and peroxidase (HRP) activity was detected as described in **4.2a**. The levels of 1d and VHH-72 V_H_H-Fcs in sera were quantified by interpolating obtained A_450nm_ readings against A_450nm_ *vs* [V_H_H-Fc] standard curves generated with purified 1d and VHH-72 V_H_H-Fcs, respectively.

#### e) Aerosolization studies

Prior to aerosolization, 4 mg of each V_H_H were purified by SEC using a Superdex 75™ 10/300 GL column (Cytiva) and PBS as running buffer, as described above. Protein fractions corresponding to the chromatogram’s monomeric peak were pooled, quantified and its concentration adjusted to 0.5 mg/mL. One mL of each V_H_H was subsequently aerosolized at room temperature with a portable mesh nebulizer (AeroNeb Solo, Aerogen, Galway, Ireland), which produces 3.4-μm particles. Aerosolized V_H_Hs were collected into 15 mL round-bottom polypropylene test tubes (VWR, Cat#C352059) for 5 min to allow condensation and were subsequently quantified and kept at 4°C until use. Then 200 µL aliquots of pre- and post- aerosolized V_H_Hs were subjected to SEC to obtain chromatogram profiles. Additionally, condensed V_H_Hs were closely monitored for the formation of any visible aggregates, and in cases where aggregate formation were observed, they were removed by centrifugation prior to concentration determination, SEC analysis and ELISA. % soluble aggregate was determined as the proportion of a V_H_H that gave elution volumes (*V*_e_s) smaller than that of the monomeric V_H_H fraction. % recovery was determined as the proportion of a V_H_H that remained monomer following aerosolization.

To assess the effect of aerosolization on functionality of V_H_Hs, the activities of post-aerosolized V_H_Hs were determined by ELISA and compared to those for pre-aerosolized V_H_Hs. To perform ELISA, S1-Fc (ACRO Biosystems, Cat#S1N-C5255) was diluted in PBS to 500 ng/mL, and 100 µL/well were coated overnight at 4°C. Next day, plates were washed with PBST and blocked with 200 µL PBSC for 1 h at room temperature. After five washes with PBST, serial dilutions of the pre- and post- aerosolized V_H_Hs were added to wells and incubated for 1 h at room temperature. Then plates were washed 10 times with PBST and binding of V_H_Hs to S1-Fc was detected with rabbit anti-6xHis Tag antibody HRP Conjugate (Bethyl Laboratories, Cat#A190-114P), diluted at 10 ng/mL in PBST and added at 100 µL/well. Finally, after 1 h incubation at room temperature and final washes with PBST, peroxidase (HRP) activity was determined as described in **4.2*a***.

### 4.8. Epitope typing, binning and mapping

#### a) Epitope typing by sodium dodecyl sulphate-polyacrylamide gel electrophoresis/western blotting (SDS-PAGE/WB)

A standard SDS-PAGE/WB was performed to detect the binding of V_H_H-Fcs to nitrocellulose-immobilized, denatured SARS-CoV-2 Wuhan S. Briefly, 10 µg/lane of S was run on 4–20% Mini-PROTEAN® TGX Stain-Free™ Protein Gels (BioRad, Cat#4568081), transferred to nitrocellulose (Sigma, Cat#GE10600002) and blocked with 1% PBSC overnight at 4℃. Then, 0.5-cm nitrocellulose strips containing the denatured S were placed on Mini Incubation Trays (BioRad, Cat#1703902) and incubated with 1 mL of 1 µg/mL V_H_H-Fcs. After 1 h incubation at room temperature, strips were washed 10 times with PBST and the binding of V_H_H-Fcs to denatured S was probed by incubating strips with 1 mL of 100 ng/mL anti-human Ig Fc antibody-peroxidase conjugate (Jackson ImmunoResearch, Cat#016-030-084) at room temperature for 1 h. Finally, strips were washed 10 times with PBST and peroxidase activity was detected using chemiluminescent reagent (SuperSignal West Pico PLUS Chemiluminescent Substrate, Thermo Fisher, Cat#34580). Images of developed strips were acquired on Molecular Imager® Gel Doc™ XR System (BioRad, Cat#1708195EDU).

#### b) Epitope binning by SPR

Standard SPR techniques were used for binding studies. All SPR assays were performed on a Biacore T200 instrument (Cytiva) at 25°C with HBS-EP running buffer (10 mM HEPES, 150 mM NaCl, 3 mM EDTA, 0.005% Tween 20, pH 7.4) and CM5 sensor chips (Cytiva). Prior to SPR analyses all analytes in flow (V_H_Hs, ACE2 receptor) were SEC-purified on a Superdex 75™ Increase 10/300 GL column (Cytiva) in HBS-EP buffer at a flow rate of 0.8 mL/min to obtain monomeric proteins. V_H_H epitope binning was performed by SPR dual injection experiments on the SARS-CoV-2 S surface at a flow rate of 40 µL/min in HBS-EP buffer. Dual injections consisted of injection of V_H_H1 (at 50 – 100 × *K*_D_ concentration) for 150 s, followed by immediate injection of a mixture of V_H_H1 + V_H_H2 (both at 50 – 100 × *K*_D_ concentration) for 150 s. The opposite orientation was also performed (V_H_H2 followed by V_H_H2 + V_H_H1). Surfaces were regenerated using a 12 s pulse of 10 mM glycine, pH 1.5, at a flow rate of 100 µL/min. All pairwise combinations of V_H_Hs were analyzed and distinct or overlapping epitope bins determined.

#### c) Epitope binning by ELISA

The pairwise ability of V_H_Hs to bind to their antigen in a sandwich ELISA format was evaluated as described previously ^85, 90, 91^ and according to the design depicted in **Fig S16B**. Briefly, a matrix of 20 well (row) × 29 well (column) in nine NUNC® Immulon 4 HBX microtiter plates (Thermo Fisher) was coated overnight at 4°C with 4 µg/mL streptavidin (Jackson ImmunoResearch, Cat#016-000-113) in 100 µL PBS, pH 7.4. Wells were blocked with 200 µL PBSC for 1 h at room temperature and then biotinylated V_H_Hs at 10 µg/mL in 100 µL PBSCT were captured in each row (same V_H_H in each row; 20 rows for a total of 20 V_H_Hs) for 1 h at room temperature. Wells were washed 5 times with PBST and incubated with 100 ng/mL of SARS-CoV-2 Wuhan S1 diluted in PBSCT for 1 h. Wells were washed and each column was incubated with the pairing, V_H_H-Fcs/ACE2-Fc at 1 µg/mL used as detector antibodies (same V_H_H-Fc/ACE2-Fc in each column; 29 columns for a total of 28 V_H_H-Fcs and ACE2-Fc). The binding of V_H_H-Fcs/ACE2-Fc to S1 was detected using 100 µL 1 µg/mL HRP-conjugated goat anti-human IgG (Sigma, Cat#A0170). Finally, plates were washed 10 times with PBST and peroxidase (HRP) activity was determined as described in **4.2*a***.

#### d) Bottom-up hydrogen-exchange mass spectrometry

All antibody:antigen complexes were equilibrated at 3:1 ratio in PBS (pH 7.1) at 4°C prior to labelling. The labelling reaction was initiated upon 10x dilution with 10 mM Tris (45% D_2_O) at 20 °C with an HDx3-PAL autosampler (Trajan Scientific and Medical, Ringwood, Australia) for a final composition of 40% D_2_O. The reaction was quenched after 3 min by a 5x dilution with 1% formic acid (FA, pH 2.2, 4°C), and 75 µL (20 pmol) was injected. Labelled sample was flowed through a µ-Prepcell (Antec Scientific, Zoeterwoude, The Netherlands) at 50 µL/min in mobile phase A (0.23% FA) where electrochemical reduction was performed in pulse mode (E1 = 1.0 V, E2 = 0 V, t1 = 1 s, t2 = 0.1 s)^92^, followed by online digestion with a pepsin (Enzymate BEH, Waters, Milford, MA) or nepenthesin-II column (Affipro, Prague, Czech Republic). Peptides were trapped (Waters ACQUITY UPLC BEH C18 Vanguard Pre-column, 130 Å, 1.7 µm, 2.1 mm x 5 mm), separated (ACQUITY UPLC BEH C18, 130 Å, 1.7 µm, 1 x 100mm) with an acetonitrile gradient and analyzed by ESI-MS (300-1600 m/z) with a Synapt G2-Si (Waters), with ion mobility enabled for the S dataset. Data-dependent MS/MS acquisition was applied to an unlabeled sample to generate a peptide map, and peptides were identified with a database search in Mascot. Replicate data was collected in triplicate in five separate batches with unique non-binding controls, and deuteration was assigned with HDExaminer v2 (Batch 1 to 4, Sierra Analytics, Modesto, CA) or MS Studio ^93^ (Batch 5). Full experimental parameters for each batch are outlined in **Table S9**. Finally, significant changes in deuteration was assigned based on two cutoffs (3 x SD and 1-p = 0.98) using MS Studio ^94^.

### 4.9. Surrogate virus neutralization assays

#### a) ACE2 competition assay by ELISA

Wells of NUNC® Immulon 4 HBX microtiter plates (Thermo Fisher) were coated overnight at 4°C with 50 ng/well of S in 100 µL PBS, pH 7.4. Next day, plates were blocked with 250 µL PBSC for 1 h at room temperature. For ACE2/V_H_H competitive binding to SARS-CoV2 S (Wuhan), 50 µL of human ACE2-Fc (ACROBiosystems, Cat#AC2-H5257) at 400 ng/mL was mixed with 50 µL of V_H_H at 1 µM, and then transferred to SARS-CoV-2 S coated microtiter plate wells. After 1 h incubation at room temperature, plates were washed 10 times with PBST and the ACE2-Fc binding was detected using 1 µg/mL goat anti-human IgG (Fc specific) HRP conjugate antibody (Sigma, Cat#A0170) in 100 µL PBSCT. After 10 washes with PBST, the peroxidase (HRP) activity was determined as described in **4.2*a***.

#### b) ACE2 competition assay by SPR

Standard SPR techniques were used for binding studies. All SPR assays were performed on a Biacore T200 instrument (Cytiva) at 25°C with HBS-EP running buffer (10 mM HEPES, 150 mM NaCl, 3 mM EDTA, 0.005% Tween 20, pH 7.4) and CM5 sensor chips (Cytiva). Prior to SPR analyses all analytes in flow (V_H_Hs, ACE2) were SEC-purified on a Superdex 75™ Increase 10/300 GL column (Cytiva) in HBS-EP buffer at a flow rate of 0.8 mL/min to obtain monomeric proteins. V_H_Hs were analyzed for their ability to block the SARS-CoV-2 Wuhan S interaction with ACE2 using SPR dual injection experiments. V_H_Hs and ACE2 were flowed over the SARS-CoV-2 S surface at 40 µL/min in HBS-EP buffer. Dual injections consisted of injection of ACE2 (1 µM) for 150 s, followed by immediate injection of a mixture of ACE2 (1 µM) + V_H_H (at 20 – 40 × *K*_D_ concentration) for 150 s. The opposite orientation was also performed (V_H_H followed by V_H_H + ACE2). Surfaces were regenerated using a 12 s pulse of 10 mM glycine, pH 1.5, at a flow rate of 100 µL/min. All pairwise combinations of V_H_Hs and ACE2 were analyzed. V_H_Hs that competed with ACE2 for SARS-CoV-2 spike glycoprotein binding showed no increase in binding response during the second injection. Conversely, a binding response was seen during the second injection for V_H_Hs that did not compete with ACE2.

#### c) ACE2 competition assay by flow cytometry

Experiments were performed as described in **4.3*c***, except that biotinylated S/Vero E6 cells were mixed with V_H_Hs or V_H_H-Fcs instead of sera. Additionally, assays were performed against biotinylated SARS-CoV-2 Wuhan, Alpha, Beta, Gamma, Delta and Kappa as well as SARS-CoV S. As internal reference of competition experiments, competition assay with recombinant human ACE2-H_6_ in lieu of V_H_H was also included. A20.1, a *C. difficile* toxin A-specific V_H_H ^82^ was used as negative control V_H_H. Percent inhibition was calculated according the formula in **4.3*c***, with *F*_n_ and *F*_max_ relating to V_H_H not serum as the competitor.

### 4.10. Pseudotyped and live virus neutralization assays

#### a) Pseudotyped virus neutralization assays

(i) Generation of SARS-CoV-2 spike pseudotyped lentiviral particles (LVP): HEK293T cells were plated in a 100-mm tissue culture dish and transfected the next day at about 75% confluency with a combination of a lentiviral transfer vector encoding eGFPLuc (addgene#119816), the packaging plasmid psPAX2 (addgene#12260), and a plasmid encoding the viral glycoprotein of interest SARS-CoV-2 Spike-ΔS1/S2-Δ20 expressed in pcDNA3.1^+^. Transfection was performed using the jetPRIME transfection reagent (Polyplus-transfection, Illkirch, France) according to the manufacturer’s protocol at a 1:1:1 (eGFPLuc:psPAX2:Spike) ratio for a total of 10 µg. The media was replaced at 24 h post transfection and complete media added to the plate. The supernatant from cell culture was harvested at 48, 72 and 96 h, each time replenished with fresh media. The combined supernatants were centrifuged at 800 *g* for 10 min and supernatants passed through a 0.45-μm syringe filter. Then, 1 volume of concentrator (40% [w/v] PEG-8000, 1.2 M NaCl, pH 7.0) was added to 3 volumes of supernatant, mixed for 1 min then incubated with constant rocking at 60 rpm for at least 4 h at 4°C. The mixture was centrifuged at 1600 *g* for 60 min at 4°C and the supernatant carefully discarded without disturbing the pellet. The pellet was then resuspended in PBS buffer at 1/10 of the original supernatant volume with gentle up-and-down pipetting, aliquoted and stored at −80°C. (ii) Viral neutralization assays: HEK293T-hACE2 cell line (BEI Resources, Manassas, VA, Cat#NR-52511) were seeded in poly-L-Lysine (PLL)-coated white, clear bottom 384-wells plate (NUNC, Thermo Fisher) at a density of 9,000 cells/well in 45 µL of media (DMEM without phenol red supplemented with 5% [v/v] FBS) and incubated for 24 h at 37°C, 5% CO_2_. The next day, a half-log serial dilution of each nanobody (V_H_H/V_H_H-Fc) to be tested was prepared at 4× the final concentration ranging from 0 to 50 µg/mL in complete media. Then, 20 µL of the 4× nanobody solution was added to each well and incubated for 10 min at 37°C, 5% CO_2_. After incubation, 20 µL of LVP master mix was added to each well and incubated a further 48 h at 37°C, 5% CO_2_. LVP master mix was prepared as follow: 10 µL of LVPs (as prepared above) were mixed with 20 µg/mL of polybrene (4× final concentration) and media for a final volume of 20 µL per well. Forty-eight hours later, 30 µL of the substrate buffer (324 mM Tris–HCl, 125 mM Tris-Base, 225 mM NaCl, 9 mM MgCl_2_, 15 mM dithiothreitol (DTT), 0.6 mM coenzyme A, 0.42 mg/mL D-luciferin, 3.3 mM ATP, 0.75% (v/v) Triton X-100, 6 mM sodium hydrosulfite) was added to each well. Then the plate was vortexed at 400 rpm for 2 min and luminescence read on a Synergy Neo2 plate reader (BioTek, Winooski, VT).

#### b) Live virus neutralizations assays

Neutralization activity of antibodies to SARS-CoV-2 was determined with the microneutralization assay. In brief, V_H_H-Fc stocks were prepared at 1 mg/mL in PBS and sterilized by passing through 0.22-µm filters. Quantitative microneutralization assay was performed on Vero E6 cells with SARS-CoV-2 strains hCOV-19/Canada/ON-VIDO-01/2020, NR-53565; hCOV- 19/England/204820464/2020, NR-54000; or hCOV-19/South Africa/KRISP-EC-K005321 /2020, NR-54008. In brief, 15 µg of antibody was diluted in 300 µL of infection media (1× DMEM, high glucose media supplemented with 1× nonessential amino acid, 100 U/mL penicillin-streptomycin, 1 mM sodium pyruvate, and 1% FBS), from which a subsequent 1:5 serial dilutions was carried out. Fifty microliters of each antibody dilution was mixed with 250 plaque-forming units (PFU) of virus in 50 µL volume. The virus/antibody mix was incubated at 37°C for 1 h. Fifty microliters of the virus/antibody mix was used to infect Vero E6 cells for 1 h at 37°C. The inoculum was removed and 100 µL of antibody dilution was added to each corresponding infected cell monolayer. Infection was incubated at 37°C for 72 h, from then they were fixed in 10% formaldehyde. Virus infection was detected with mouse mAb to SARS-CoV- 2 nucleocapsid (N) (R&D Systems, Minneapolis, MN, Cat#MAB10474) and counterstained with rabbit anti-mouse IgG-HRP (Rockland Immunochemicals, Inc., Limerick, PA) and developed with *o*- Phenylenediamine dihydrochloride (Sigma) and detected on BioTek Synergy H1 microplate reader at 490 nm. *IC*_50_ was determined from non-linear regression [Inhibitor] vs. response, Variable slope (four parameters) using GraphPad Prism version 9 (La Jolla, CA).

### 4.11. *In vivo* efficacy studies

#### a) In vivo challenge

Female Golden Syrian hamsters (81 – 90 g) were obtained from Charles River Laboratories (Saint-Constant, Canada) and maintained at the NRC small animal facility. All animal procedures and animal husbandry in this study were carried out in accordance to regulations and guidelines outlined under the Canadian Council on Animal Care and approved by the NRC Human Health Therapeutics Animal Care Committee. Sixty hamsters (six /treatment group) were prebled for 300 µL of blood followed by IP infusion with 1 mg of antibodies or with PBS 24 h before challenge. Baseline body weights were determined before challenge. The animals were then infected intranasally under ketamine/xylazine with 8.5 x 10^4^ PFU (in 100 µL) of SARS-CoV-2. Daily weight and clinical symptoms were monitored for 5 days post-infection (dpi). At 5 dpi, animals were euthanized and their lungs were collected for virus titers determination and immunohistochemistry studies. Virus titer was determined by plaque assay on Vero cells.

#### b) Immunohistochemistry

Lungs were immersed in 10% neutral buffered formalin and fixed for 1 week at room temperature and then transferred into 70% ethanol. All 4 lobes of right lung were processed and embedded in paraffin wax. The paraffin block was cut into 5-µm sections and placed on Superfrost Plus slides (Thermo Fisher). Sections were dried overnight and then subjected to immunohistochemistry (IHC) using a modified protocol F on the Bond-Max III fully automated staining system (Leica Biosystems, Wetzlar, Germany). All reagents from the Bond Polymer Refine Detection Kit DC9800 (Leica, Buffalo Grove, IL) were used for IHC. Immune cell infiltrate was detected using rabbit polyclonal antibodies against CD3 (1:500, Dako, Cat#A0452), and Iba-1 (ionized calcium binding adaptor protein, 1:5000, Dako, Cat#019-19741). Mouse anti-SARS-CoV-2 N monoclonal antibody (1:5000, R&D Systems, Cat#MAB10474) was used for the detection of SARS-CoV-2. Following deparaffinization and rehydration, sections were pre-treated with the Epitope Retrieval Solution 1 (ER1, Citrate buffer, pH 5.0) or Epitope Retrieval Solution 2 (ER2, EDTA buffer, pH 8.8) at 98°C for 20 min. Epitopes were exposed using ER1 for Iba-1 and SARS-CoV-2 N and ER2 for CD3. After washing steps, non-specific endogenous peroxidases were quenched using peroxidase block for 5 min. Sections were washed again and then incubated for 15 min at room temperature with primary antibodies. After washes, sections were incubated with polymer refine for 8 min at room temperature and developed with 3, 3’-diaminobenzidine (DAB) chromogen for 10 min. Sections were washed and counterstained for 5 min with hematoxylin, dehydrated, cleared and mounted. Each antibody was titrated and optimized for the detection of positive signals. Negative controls included omission of primary antibody and incubation with secondary antibody alone as well as lung tissue from naïve animals.

The DeadEnd Colorimetric TUNEL (<u>T</u>dT-mediated d<u>U</u>TP <u>N</u>ick-<u>E</u>nd Labeling) System (Promega Madison, WI, Cat#G7132) was used to detect apoptosis as per manufacturer’s instructions. Briefly, sections were deparaffinized and permeabilized with proteinase K for 15 min followed by labeling with biotinylated nucleotides in the presence of recombinant terminal deoxynucleotidyl tranferase. Sections were then incubated with horseradish peroxidase-streptavidin conjugate and developed using DAB for 10 min. Sections were washed and counterstained for 5 min with hematoxylin, dehydrated, cleared and mounted.

All images were acquired with an Olympus IX81 microscope, equipped with a DP27 color CCD camera using the 20× objective (Shinjuku, Tokyo, Japan).

## Acknowledgements

We thank Bassam Hallis for contributing SARS-Related Coronavirus 2, Isolate hCoV-19/England/204820464/2020, NR-54000, and Alex Sigal and Tulio de Oliveira for contributing SARS-Related Coronavirus 2, Isolate hCoV-19/South Africa/KRISP-EC-K005321/2020, NR-54008; both virus isolates were obtained through BEI Resources, NIAID, NIH. We thank the NRC Animal Resources team for animal husbandry and technical support, and Luc Lemay and Jennifer Wellman for technical support in the CL3. We thank Sonia Leclerc and Julie Haukenfrers for performing DNA sequencing. We thank Qingling Yang and Shannon Ryan for assistance with expression of ACE2-H_6_ and His tagged and Fc-fused RBD/SD1, and Mary Foss for assistance with aerosolization studies. We thank the NRC Mammalian Cell Expression Section for assistance with production of recombinant proteins and Joline Cormier for contribution to protein samples management. We thank Alina Burlacu for generating the CHO-SPK cells and Tahir Maqbool for his support with llama immunizations. We thank Unchained Labs (Pleasanton, CA) for conducting isothermal chemical denaturation experiments. This work was supported by funding from the Pandemic Response Challenge Program of the National Research Council Canada and in part by a CIHR grant (OV1 - 170355) to M.A.L.

