## Supplemental Tables for "Arsenal of Nanobodies for Broad-Spectrum Countermeasures against Current and Future SARS-CoV-2 Variants of Concerns"

**Table S1.** Coronavirus spike glycoprotein fragments and ACE2 used in this study

| Description | Accession number | Source | Tag | Reference describing expression & purification |
|---|---|---|---|---|
| S1 (aa16-685) (Wuhan) | QHD43416.1[b] | in-house | FLAG, 6xHis | Akache et al.,2021[1] |
| S1 (aa16-685) (Wuhan) | QHD43416.1[b] | ACROBiosystems | AviTag, 6xHis | na |
| S1 (aa16-685) (Wuhan) | QHD43416.1[b] | ACROBiosystems | Human IgG1 Fc | na |
| NTD (aa16-305) (Wuhan) | QHD43416.1[b] | in-house | FLAG, 6xHis | Akache et al.,2021[1] |
| RBD/SD1 (aa319-591) (Wuhan) | QHD43416.1[b] | in-house | Human IgG1 Fc | Wrapp et al., 2020[2] |
| RBD/SD1 (aa319-591) (Wuhan) | QHD43416.1[b] | in-house | 6xHis | Wrapp et al., 2020[2] |
| RBD_short (aa331-521) (Wuhan) | QHD43416.1[b] | in-house | 6xHis | Akache et al.,2021[1] |
| RBD (aa319-541) (Wuhan) | QHD43416.1[b] | in-house | FLAG-6xHis (N-term), E5 (C-term) | Akache et al.,2021[1] |
| RBD_α (aa319-541) (B.1.1.7) | QHD43416.1[b] | in-house | FLAG, 6xHis | Colwill et al., 2021[3] |
| RBD_β (aa319-541) (B.1.351) | QHD43416.1[b] | in-house | FLAG, 6xHis | Colwill et al., 2021[3] |
| RBD_Wuhan (aa319-541) | QHD43416.1[b] | ACROBiosystems | AviTag, 6xHis | na |
| S2 (aa686-1208) (Wuhan) | QHD43416.1[b] | in-house | FLAG, 6xHis | Akache et al.,2021[1] |
| Swine deltaCoV S[a] | AIH06857.1[b] | in-house | FLAG-Dual Strep-6xHis | Galipeau et al., 2021[4] |
| Avian_IBV S[a] | AAP92675.1[b] | in-house | FLAG-Dual Strep-6xHis | Galipeau et al., 2021[4] |
| Pangolin CoV S[a] | QIA48632.1[b] | in-house | FLAG-Dual Strep-6xHis | Galipeau et al., 2021[4] |
| Hedgehog CoV HKU31 S[a] | QGA70702.0[b] | in-house | FLAG-Dual Strep-6xHis | Galipeau et al., 2021[4] |
| Bat CoV HKU9 S[a] | YP_001039971.1[c] | in-house | FLAG-Dual Strep-6xHis | Galipeau et al., 2021[4] |
| Bat SARS like CoV-WIV1 S[a] | AGZ48828.1[b] | in-house | FLAG-Dual Strep-6xHis | Galipeau et al., 2021[4] |
| Bat 229E-related CoV S[a] | APD51507.1[b] | in-house | FLAG-Dual Strep-6xHis | Galipeau et al., 2021[4] |
| Bat CoV 512 S[a] | YP_001351684.1[c] | in-house | FLAG-Dual Strep-6xHis | Galipeau et al., 2021[4] |
| Bat SARS like CoV[a] | ATO98157.1[b] | in-house | FLAG-Dual Strep-6xHis | Galipeau et al., 2021[4] |
| Civet SARS-CoV[1] S[a] | AAU04646.1[b] | in-house | FLAG-Dual Strep-6xHis | Galipeau et al., 2021[4] |
| Human MERS-CoV S[a] | AGH58717.1[b] | in-house | FLAG-Dual Strep-6xHis | Galipeau et al., 2021[4] |
| Human CoV-NL63 S | APF29071.1[b] | Sino Biological | 6xHis | na |
| Human CoV-OC43 S[a] | AGT51431.1[b] | in-house | FLAG-Dual Strep-6xHis | Galipeau et al., 2021[4] |
| Human CoV-HKU1 S | Q0ZME7.1[d] | Sino Biological | 6xHis | na |
| Human CoV-229E S[a] | AAK32191.1[b] | Sino Biological | 6xHis | na |
| Human SARS-CoV S[a] | AAP13442.1[b] | in-house | FLAG-Dual Strep-6xHis | Galipeau et al., 2021[4] |

| | | | | |
|---|---|---|---|---|
| Human SARS-CoV-2_Wuhan S[a] (SmT1) | QHD43416.1[b] | in-house | FLAG, 6xHis | Colwill et al., 2021[3] |
| Human SARS-CoV-2_α (B.1.351) S[a] | - | in-house | FLAG, 6xHis | Galipeau et al., 2021[4] |
| Human SARS-CoV-2_β (B. 1.1.7) S[a] | - | in-house | FLAG, 6xHis | Galipeau et al., 2021[4] |
| Human SARS-CoV-2_g (P.1) S[a] | - | in-house | none | Galipeau et al., 2021[4] |
| Human SARS-CoV-2_D (B.1.617.2) S[a] | - | in-house | FLAG, 6xHis | Stuible et al., 2021[5] |
| Human SARS-CoV-2_k (B.1.617.1) S[a] | - | in-house | FLAG, 6xHis | Stuible et al., 2021[5] |
| Human ACE2 (aa18-740) | Q9BYF1-1[d] | ACROBiosystems | Human IgG1 Fc | na |
| Human ACE2 (aa18-615) | Q9BYF1-1[d] | in-house | 6xHis | Wrapp et al., 2020[2] |

[a]Proteins are C-terminally fused to the resistin trimerization domain. [b]GenBank; [c]NCBI; [d]UniProt. na, not applicable

**Table S2.** Kinetic and equilibrium constants for $V_H$Hs binding to SARS-CoV-2 spike glycoprotein fragments

| $V_H$H/ACE2 | RBD_short[a] | | | RBD/SD1[a] | | | S[a] | | | S1[a] | | | S2[a] | | |
|---|---|---|---|---|---|---|---|---|---|---|---|---|---|---|---|
| | $k_a$ (1/Ms) | $k_d$ (1/s) | $K_D$ (M) | $k_a$ (1/Ms) | $k_d$ (1/s) | $K_D$ (M) | $k_a$ (1/Ms) | $k_d$ (1/s) | $K_D$ (M) | $k_a$ (1/Ms) | $k_d$ (1/s) | $K_D$ (M) | $k_a$ (1/Ms) | $k_d$ (1/s) | $K_D$ (M) |
| **RBD-specific $V_H$H** | | | | | | | | | | | | | | | |
| 1d | 9.26E+05 | 3.76E-03 | 4.06E-09 | 5.62E+05 | 1.18E-03 | 2.10E-09 | 1.06E+06 | 1.17E-03 | 1.10E-09 | 8.67E+05 | 1.14E-03 | 1.32E-09 | - | - | - |
| 02 | 2.01E+06 | 1.73E-03 | 8.60E-10 | 1.83E+06 | 1.41E-03 | 7.73E-10 | 2.14E+06 | 1.41E-03 | 6.61E-10 | 2.10E+06 | 1.38E-03 | 6.59E-10 | - | - | - |
| 03 | - | - | - | 1.66E+05 | 2.55E-04 | 1.53E-09 | 2.50E+05 | 3.34E-04 | 1.34E-09 | 2.41E+05 | 2.63E-04 | 1.09E-09 | - | - | - |
| 04 | 1.40E+06 | 2.02E-02 | 1.45E-08 | 1.53E+06 | 1.92E-02 | 1.25E-08 | 1.97E+06 | 1.98E-02 | 1.00E-08 | 1.61E+06 | 1.80E-02 | 1.12E-08 | - | - | - |
| 05 | 2.97E+06 | 7.13E-03 | 2.40E-09 | 2.51E+06 | 5.85E-03 | 2.33E-09 | 4.33E+06 | 7.43E-03 | 1.72E-09 | 4.03E+06 | 8.06E-03 | 2.00E-09 | - | - | - |
| 06 | 9.83E+03 | 6.87E-03 | 6.99E-07 | 5.94E+04 | 4.15E-03 | 6.99E-08 | 3.02E+04 | 4.69E-03 | 1.55E-07 | 1.04E+05 | 1.33E-02 | 1.29E-07 | - | - | - |
| 07 | 4.25E+05 | 4.54E-04 | 1.07E-09 | 3.18E+05 | 2.84E-04 | 8.94E-10 | 3.50E+05 | 4.03E-04 | 1.15E-09 | 3.01E+05 | 3.08E-04 | 1.02E-09 | - | - | - |
| 10 | 3.79E+05 | 1.01E-04 | 2.66E-10 | 3.51E+05 | 9.32E-05 | 2.66E-10 | 4.83E+05 | 9.27E-05 | 1.92E-10 | 4.48E+05 | 9.54E-05 | 2.13E-10 | - | - | - |
| 11 | - | - | - | 8.92E+05 | 2.26E-04 | 2.53E-10 | 1.21E+06 | 4.73E-05 | 3.91E-11 | 1.08E+06 | 1.80E-04 | 1.67E-10 | - | - | - |
| 12 | 5.33E+05 | 4.78E-05 | 8.97E-11 | 7.60E+05 | 3.69E-05 | 4.86E-11 | | | | | nd | | | | |
| 14 | 2.98E+05 | 1.11E-03 | 3.71E-09 | 2.61E+05 | 9.44E-04 | 3.61E-09 | 5.28E+05 | 1.64E-03 | 3.10E-09 | 3.19E+05 | 1.12E-03 | 3.49E-09 | - | - | - |
| 15 | 9.01E+05 | 2.89E-04 | 3.21E-10 | 6.82E+05 | 2.33E-04 | 3.42E-10 | 7.06E+05 | 2.21E-04 | 3.13E-10 | 6.75E+05 | 2.25E-04 | 3.33E-10 | - | - | - |
| 17 | 9.53E+05 | 3.48E-04 | 3.65E-10 | 5.67E+05 | 2.24E-04 | 3.95E-10 | 6.59E+05 | 9.79E-05 | 1.49E-10 | 6.29E+05 | 9.66E-05 | 1.53E-10 | - | - | - |
| 18 | 3.35E+05 | 1.80E-03 | 5.37E-09 | 2.68E+05 | 1.97E-04 | 7.36E-10 | 3.95E+05 | 1.52E-04 | 3.84E-10 | 3.08E+05 | 1.65E-04 | 5.36E-10 | - | - | - |
| 20 | 2.67E+06 | 2.10E-02 | 7.84E-09 | 1.43E+06 | 1.23E-02 | 8.61E-09 | 2.37E+06 | 1.59E-02 | 6.73E-09 | 1.96E+06 | 1.89E-02 | 9.63E-09 | - | - | - |
| MRed04 | 1.58E+06 | 1.08E-02 | 6.86E-09 | 6.04E+05 | 1.51E-03 | 2.51E-09 | 1.39E+06 | 1.50E-03 | 1.09E-09 | 1.09E+06 | 1.83E-03 | 1.68E-09 | - | - | - |
| MRed05 | 4.68E+05 | 6.33E-04 | 1.36E-09 | 6.50E+05 | 4.96E-04 | 7.63E-10 | | | | | nd | | | | |
| **NTD-specific $V_H$H** | | | | | | | | | | | | | | | |
| SR01 | - | - | - | - | - | - | 2.77E+06 | 1.23E-03 | 4.45E-10 | 3.63E+06 | 1.58E-03 | 4.37E-10 | - | - | - |
| SR02 | - | - | - | - | - | - | 9.67E+05 | 5.71E-04 | 5.90E-10 | 1.05E+06 | 5.55E-04 | 5.30E-10 | - | - | - |
| SR03 | - | - | - | - | - | - | 1.01E+06 | 1.02E-03 | 1.01E-09 | 9.05E+05 | 1.03E-03 | 1.13E-09 | - | - | - |
| SR04 | - | - | - | - | - | - | 2.39E+06 | 3.35E-04 | 1.40E-10 | 2.18E+06 | 4.93E-04 | 2.26E-10 | - | - | - |
| SR13 | - | - | - | - | - | - | 1.83E+06 | 4.81E-03 | 2.62E-09 | 2.25E+06 | 5.45E-03 | 2.43E-09 | - | - | - |
| SR16 | - | - | - | - | - | - | 6.57E+05 | 1.20E-03 | 1.82E-09 | 7.39E+05 | 1.15E-03 | 1.55E-09 | - | - | - |

| | | | | | | | | | | | | | | | |
|---|---|---|---|---|---|---|---|---|---|---|---|---|---|---|---|
| MRed03 | - | - | - | - | - | - | 1.57E+05 | 8.14E-05 | 5.20E-10 | 2.76E+05 | 6.15E-05 | 2.23E-10 | - | - | - |
| MRed06 | - | - | - | - | - | - | 1.85E+05 | 8.87E-04 | 4.80E-09 | 3.71E+05 | 1.57E-03 | 4.22E-09 | - | - | - |
| MRed07 | - | - | - | - | - | - | 1.60E+06 | 3.78E-04 | 2.36E-10 | 1.13E+06 | 4.27E-04 | 3.78E-10 | - | - | - |
| **S2-specific $V_HH$** | | | | | | | | | | | | | | | |
| S2A3 | - | - | - | - | - | - | 8.40E+04 | 1.30E-04 | 1.55E-09 | - | - | - | 4.60E+04 | 1.03E-04 | 2.23E-09 |
| S2A4 | - | - | - | - | - | - | 3.49E+04 | 4.46E-04 | 1.28E-08 | - | - | - | 2.81E+04 | 4.14E-04 | 1.47E-08 |
| S2F3 | - | - | - | - | - | - | 1.56E+05 | 4.73E-04 | 3.03E-09 | - | - | - | 1.01E+05 | 6.29E-04 | 6.22E-09 |
| S2G3 | - | - | - | - | - | - | 1.62E+05 | 6.07E-04 | 3.74E-09 | - | - | - | 1.47E+05 | 6.27E-04 | 4.28E-09 |
| S2G4 | - | - | - | - | - | - | 8.93E+05 | 2.07E-04 | 2.32E-10 | - | - | - | 9.25E+05 | 3.82E-04 | 4.13E-10 |
| MRed11 | - | - | - | - | - | - | 2.34E+04 | 4.18E-04 | 1.78E-08 | - | - | - | 4.09E+04 | 2.13E-04 | 5.21E-09 |
| MRed18 | - | - | - | - | - | - | 2.02E+05 | 1.53E-03 | 7.56E-09 | - | - | - | 1.52E+05 | 1.03E-03 | 6.80E-09 |
| MRed19 | - | - | - | - | - | - | 1.59E+05 | 7.99E-04 | 5.01E-09 | - | - | - | 1.02E+05 | 8.40E-04 | 8.20E-09 |
| MRed20 | - | - | - | - | - | - | 1.60E+05 | 1.46E-05 | 9.18E-11 | - | - | - | 1.13E+05 | 3.38E-05 | 2.99E-10 |
| MRed22 | - | - | - | - | - | - | 3.47E+05 | 1.76E-04 | 5.06E-10 | - | - | - | 2.70E+05 | 2.84E-04 | 1.05E-09 |
| MRed25 | - | - | - | - | - | - | 1.12E+05 | 1.15E-04 | 1.02E-09 | - | - | - | 1.22E+05 | 1.90E-04 | 1.56E-09 |
| **Reference** | | | | | | | | | | | | | | | |
| VHH-72[b] | 8.64E+05 | 4.22E-01 | 4.89E-07 | 6.67E+05 | 1.34E-01 | 2.00E-07 | 1.10E+06 | 1.56E-01 | 1.42E-07 | 9.40E+05 | 1.46E-01 | 1.56E-07 | - | - | - |
| ACE-2[b] | 5.92E+04 | 1.35E-02 | 2.28E-07 | 3.71E+04 | 1.18E-02 | 3.17E-07 | 6.02E+04 | 9.96E-03 | 1.65E-07 | 6.21E+04 | 1.24E-02 | 2.00E-07 | - | - | - |
| NRCsdAb022[b] | - | - | - | - | - | - | - | - | - | - | - | - | - | - | - |

[a]For any given $V_HH$, $K_D$ values across different spike fragments were essentially consistent, except for $V_HH$ 03/11 and 06/18 which respectively showed no or ~10-fold lower binding to RBD_short compared to RBD/SD1 and S1. Lack of $V_HH$ binding for certain spike fragments are consistent with $V_HH$s' subunit/domain specificities. Binding parameters were determined by flowing monomeric $V_HH$s over sensorchip surfaces immobilized with various spike fragments, except for $V_HH$ 12 and MRed05, which were obtained by flowing monomeric RBDs over $V_HH$-Fc-captured surfaces (for 12, RBD [aa319-541] instead of RBD/SD1 was used). Dashes indicate lack of binding. See **Table S1** for the primary structures of spike glycoprotein fragments. [b]ACE2-$H_6$ and VHH-72[6], positive controls, EGFR-specific $V_HH$ NRCsdAb022[7], negative control.

**Table S3.** ELISA data for the binding of $V_H$H-Fcs to SARS-CoV-2 spike glycoprotein fragments

| $V_H$H-Fc[a] | $EC_{50}$app[b] (nM) | | | | Domain specificity |
|---|---|---|---|---|---|
| | S | S1 | NTD | RBD | |
| SR01 | 0.1 | 0.2 | 0.2 | - | NTD |
| SR02 | 0.1 | 0.1 | 0.3 | - | NTD |
| SR03 | 0.2 | 0.2 | 0.2 | - | NTD |
| SR04 | 0.2 | 0.3 | 0.2 | - | NTD |
| SR13 | 0.4 | 0.4 | 0.2 | - | NTD |
| SR16 | 0.6 | 2.7 | 0.4 | - | NTD |
| MRed03 | 0.2 | 0.5 | 0.7 | - | NTD |
| MRed06 | 0.4 | 0.6 | 0.4 | - | NTD |
| MRed07 | 1.2 | 1.5 | 0.8 | - | NTD |
| 02 | 0.1 | 0.1 | - | 0.2 | RBD |

[a]S1-specfic $V_H$Hs that did not bind to RBD by SPR (**Table S2**) were tested for specificity against recombinant NTD domain (**Table S1**). The RBD-specific $V_H$H 02 internal control gave the expected binding specificity profile; [b]$EC_{50}$app, apparent $EC_{50}$.

**Table S4.** Kinetic and equilibrium dissociation constants for the binding of $V_H$Hs to SARS-CoV-2 Wuhan, SARS-CoV-2 Alpha, SARS-CoV-2 Beta and SARS-CoV spike glycoprotein fragments

| $V_H$H/ACE2 | SARS-CoV-2 Wuhan[a] | | | SARS-CoV-2 Alpha[a] | | | SARS-CoV-2 Bet[a] | | | SARS-CoV[a] | | |
|---|---|---|---|---|---|---|---|---|---|---|---|---|
| | $k_a$ (1/Ms) | $k_d$ (1/s) | $K_D$ (M) | $k_a$ (1/Ms) | $k_d$ (1/s) | $K_D$ (M) | $k_a$ (1/Ms) | $k_d$ (1/s) | $K_D$ (M) | $k_a$ (1/Ms) | $k_d$ (1/s) | $K_D$ (M) |
| **RBD-specific $V_H$H** | | | | | | | | | | | | |
| 1d | 1.39E+06 | 1.05E-03 | 7.50E-10 | 1.13E+06 | 1.02E-03 | 9.07E-10 | 8.12E+05 | 9.56E-04 | 1.18E-09 | - | - | - |
| 02 | 2.04E+06 | 1.27E-03 | 6.24E-10 | 1.64E+06 | 2.22E-02 | 1.36E-08 | - | - | - | - | - | - |
| 03 | 1.16E+05 | 1.81E-04 | 1.56E-09 | 1.03E+05 | 1.54E-04 | 1.49E-09 | 5.54E+04 | 2.26E-04 | 4.08E-09 | - | - | - |
| 04 | 1.84E+06 | 1.88E-02 | 1.02E-08 | 1.67E+06 | 1.96E-02 | 1.17E-08 | - | - | - | - | - | - |
| 05 | 2.76E+06 | 7.09E-03 | 2.57E-09 | 2.28E+06 | 2.61E-02 | 1.14E-08 | - | - | - | - | - | - |
| 06 | 2.05E+04 | 4.56E-03 | 2.23E-07 | 2.05E+04 | 4.68E-03 | 2.29E-07 | 2.48E+04 | 6.14E-03 | 2.48E-07 | - | - | - |
| 07 | 3.78E+05 | 3.55E-04 | 9.39E-10 | 3.39E+05 | 3.78E-04 | 1.12E-09 | 3.05E+05 | 3.21E-04 | 1.05E-09 | 1.20E+05 | 1.46E-03 | 1.22E-08 |
| 10 | 5.81E+05 | 1.16E-04 | 1.99E-10 | 4.84E+05 | 1.02E-04 | 2.10E-10 | 2.70E+05 | 2.62E-03 | 9.73E-09 | - | - | - |
| 11 | 9.64E+05 | 1.71E-05 | 1.77E-11 | 9.28E+05 | 1.59E-05 | 1.71E-11 | 7.57E+05 | 1.72E-05 | 2.27E-11 | 1.93E+06 | 2.70E-05 | 1.40E-11 |
| 12[a] | 7.47E+05 | 3.50E-05 | 4.69E-11 | 9.47E+05 | 4.38E-05 | 4.63E-11 | 9.70E+05 | 3.89E-05 | 4.01E-11 | | nd | |
| 14 | 3.89E+05 | 1.01E-03 | 2.60E-09 | 3.75E+05 | 9.17E-04 | 2.44E-09 | - | - | - | - | - | - |
| 15 | 6.91E+05 | 2.22E-04 | 3.21E-10 | 6.37E+05 | 1.94E-04 | 3.05E-10 | 1.48E+05 | 3.28E-03 | 2.22E-08 | - | - | - |
| 17 | 6.14E+05 | 9.45E-05 | 1.54E-10 | 6.14E+05 | 7.64E-05 | 1.25E-10 | 1.14E+06 | 5.88E-03 | 5.14E-09 | - | - | - |
| 18 | 2.95E+05 | 9.39E-05 | 3.18E-10 | 2.78E+05 | 9.79E-05 | 3.53E-10 | 2.76E+05 | 1.01E-04 | 3.65E-10 | - | - | - |
| 20 | 2.64E+06 | 1.16E-02 | 4.39E-09 | 2.48E+06 | 1.23E-02 | 4.97E-09 | 2.08E+06 | 1.14E-02 | 5.47E-09 | - | - | - |
| MRed04 | 1.51E+06 | 1.31E-03 | 8.63E-10 | 1.34E+06 | 1.22E-03 | 9.09E-10 | 1.14E+06 | 1.23E-03 | 1.07E-09 | 3.23E+05 | 9.70E-02 | 3.00E-07 |
| MRed05[a] | 5.62E+05 | 5.09E-04 | 9.05E-10 | 5.73E+05 | 1.76E-04 | 3.07E-10 | 6.12E+05 | 5.44E-04 | 8.88E-10 | | nd | |
| **NTD-specific $V_H$H** | | | | | | | | | | | | |
| SR01 | 1.43E+06 | 8.05E-04 | 5.64E-10 | 1.01E+06 | 5.94E-04 | 5.91E-10 | 1.09E+06 | 2.21E-04 | 2.02E-10 | 6.81E+05 | 1.05E-04 | 1.54E-10 |
| SR02 | 4.98E+06 | 6.69E-04 | 1.35E-10 | 4.54E+06 | 2.54E-04 | 5.59E-11 | 4.19E+06 | 6.28E-04 | 1.50E-10 | - | - | - |
| SR03 | 6.70E+05 | 1.13E-03 | 1.69E-09 | 5.55E+05 | 9.55E-04 | 1.72E-09 | 5.64E+05 | 1.40E-03 | 2.49E-09 | - | - | - |
| SR04 | 3.68E+06 | 5.15E-04 | 1.40E-10 | 2.80E+06 | 7.50E-04 | 2.67E-10 | 2.24E+06 | 7.23E-04 | 3.24E-10 | - | - | - |
| SR13 | 1.06E+06 | 3.76E-03 | 3.56E-09 | 5.92E+05 | 3.44E-03 | 5.82E-09 | 5.71E+05 | 4.01E-03 | 7.02E-09 | - | - | - |
| SR16 | 4.88E+05 | 9.57E-04 | 1.96E-09 | 4.04E+05 | 6.33E-04 | 1.57E-09 | 3.95E+05 | 1.01E-03 | 2.57E-09 | - | - | - |
| MRed03 | 2.37E+05 | 1.21E-04 | 5.08E-10 | 2.71E+05 | 9.89E-05 | 3.64E-10 | 2.13E+05 | 1.43E-04 | 6.72E-10 | - | - | - |

| | | | | | | | | | | | | |
|---|---|---|---|---|---|---|---|---|---|---|---|---|
| MRed06 | 1.92E+05 | 9.96E-04 | 5.19E-09 | 2.27E+05 | 1.30E-03 | 5.72E-09 | 1.39E+05 | 1.01E-03 | 7.24E-09 | - | - | - |
| MRed07 | 4.58E+06 | 4.81E-04 | 1.05E-10 | 3.90E+06 | 1.03E-03 | 2.64E-10 | 2.38E+06 | 5.52E-04 | 2.31E-10 | - | - | - |
| **S2-specific $V_HH$** | | | | | | | | | | | | |
| S2A3 | 9.83E+04 | 5.51E-05 | 5.61E-10 | 8.70E+04 | 1.89E-04 | 2.18E-09 | 6.69E+04 | 5.71E-05 | 8.53E-10 | - | - | - |
| S2A4 | 3.49E+04 | 4.46E-04 | 1.28E-08 | | nd | | | nd | | - | - | - |
| S2F3 | 1.56E+05 | 4.73E-04 | 3.03E-09 | | nd | | | nd | | 2.82E+05 | 1.39E-03 | 4.91E-09 |
| S2G3 | 3.24E+05 | 6.06E-04 | 1.87E-09 | 2.98E+05 | 5.30E-04 | 1.78E-09 | 3.04E+05 | 5.63E-04 | 1.85E-09 | 1.87E+05 | 8.00E-04 | 4.27E-09 |
| S2G4 | 8.93E+05 | 2.07E-04 | 2.32E-10 | | nd | | | nd | | 9.20E+05 | 7.35E-04 | 7.99E-10 |
| MRed11 | 4.57E+04 | 2.83E-04 | 6.20E-09 | 3.11E+04 | 4.26E-04 | 1.37E-08 | 4.54E+04 | 2.82E-04 | 6.21E-09 | - | - | - |
| MRed18 | 1.97E+05 | 1.19E-03 | 6.03E-09 | 3.82E+05 | 4.93E-03 | 1.29E-08 | 3.69E+05 | 2.39E-03 | 6.48E-09 | 3.00E+05 | 6.77E-03 | 2.25E-08 |
| MRed19 | 1.31E+05 | 1.18E-03 | 9.07E-09 | 1.83E+05 | 3.70E-03 | 2.02E-08 | 1.29E+05 | 1.04E-03 | 8.07E-09 | 2.49E+05 | 6.14E-03 | 2.46E-08 |
| MRed20 | 1.60E+05 | 1.46E-05 | 9.18E-11 | | nd | | | nd | | 3.80E+05 | 4.06E-03 | 1.07E-08 |
| MRed22 | 3.47E+05 | 1.76E-04 | 5.06E-10 | | nd | | | nd | | - | - | - |
| MRed25 | 1.12E+05 | 1.15E-04 | 1.02E-09 | | nd | | | nd | | 2.18E+04 | 5.01E-05 | 2.29E-09 |
| **Control** | | | | | | | | | | | | |
| $ACE2\text{-}H_6$[b] | 6.38E+04 | 9.79E-03 | 1.53E-07 | 8.52E+04 | 1.56E-03 | 1.83E-08 | 3.66E+04 | 4.78E-03 | 1.31E-07 | 1.11E+05 | 3.89E-02 | 3.51E-07 |
| VHH-72[b] | 1.23E+06 | 1.06E-01 | 8.62E-08 | 1.05E+06 | 1.01E-01 | 9.60E-08 | 8.01E+05 | 9.92E-02 | 1.24E-07 | 1.01E+06 | 6.56E-03 | 6.52E-09 |
| NRCsdAb022[b] | - | - | - | - | - | - | - | - | - | - | - | - |

[a]Binding parameters were determined by flowing monomeric $V_HH$s over sensorchip surfaces immobilized with S, except for $V_HH$ 12 and MRed05, which were obtained by flowing monomeric RBDs (aa319-541) over $V_HH$-Fc-captured surfaces. Dashes indicate lack of binding. nd, not determined. See **Table S1** for the primary structures of spike glycoprotein fragments; [b]$ACE2\text{-}H_6$ and VHH-72[6], positive controls, EGFR-specific $V_HH$ NRCsdAb022[7] negative control.

**Table S5.** Summary of $V_HH$ stability data

| $V_HH$ | Temperature-induced denaturation | GdnHCl-induced denaturation | | |
|---|---|---|---|---|
| | $T_m$ (°C) | $\Delta G^0$ (kJ/mol) | $C_m$ (M) | *m* (kJ/M*mol) |
| 1d | 65.5 | 27.1 | 1.8 | 15.4 |
| 02 | 67.3 | 31.3 | 1.9 | 16.9 |
| 03 | 79.5 | nd | nd | nd |
| 04 | 74.9 | 30.9 | 2.1 | 14.8 |
| 05 | 70.1 | 37.4 | 2.2 | 16.7 |
| 06 | 71.9 | 44 | 2.3 | 18.7 |
| 07 | 79.8 | 31.3 | 2.3 | 13.6 |
| 10 | 68.8 | 35.8 | 2.1 | 17.1 |
| 11 | 60.4 | 21.4 | 1.6 | 13.1 |
| 14 | 68.9 | 27.5 | 2.2 | 12.3 |
| 15 | 76.9 | 35.9 | 2.2 | 16 |
| 17 | 73.3 | 35 | 2.6 | 13.6 |
| 18[a] | 67.3 | nd | 0.3 | nd |
| 20 | 76.1 | nd | nd | nd |
| MRed04 | 71.3 | 24.6 | 2.0 | 12.1 |
| MRed05 | 69.7 | nd | nd | nd |
| SR01 | 77.4 | nd | nd | nd |
| SR03 | 76.3 | 53.4 | 3.1 | 17.5 |
| SR04 | 66.5 | 30.5 | 1.9 | 15.9 |
| SR13 | 69.7 | 28.5 | 2.0 | 14 |
| S2A3 | 61.2 | nd | nd | nd |
| S2A4 | 70.4 | 29.5 | 1.9 | 15.4 |
| S2F3 | 64.6 | nd | nd | nd |
| S2G3 | 70.4 | nd | nd | nd |
| S2G4 | 69.3 | 28.4 | 2.2 | 12.9 |
| MRed03 | 79.8 | 25.4 | 2.5 | 10.3 |
| MRed07 | 68.4 | 26 | 2.2 | 12.1 |

| | | | | |
|---|---|---|---|---|
| MRed11 | 65.1 | nd | nd | nd |
| MRed18 | 72.5 | nd | nd | nd |
| MRed19 | 69.1 | nd | nd | nd |
| MRed20 | 77.1 | 43.1 | 2.2 | 19.8 |
| MRed22 | 72.5 | 31.3 | 2.2 | 14.3 |
| MRed25 | 70.5 | 25.9 | 2.4 | 10.7 |
| VHH-72 | 73 | 30.9 | 2.3 | 13.6 |

[a]Lack of a lower plateau due to an immediate melting upon exposure to the denaturant did not allow for a reliable curve fitting, hence $\Delta G^0$ and *m* values for $V_HH$ 18 could not be obtained. nd, not determined.

**Table S6.** Stability of $V_HH$s against aerosolization

| $V_HH$ | Recovery (%)[a] | Soluble aggregates (%)[b] | | ΔSoluble agg.[c] | Visible aggregates |
|---|---|---|---|---|---|
| | | Pre-aerosolization | Post-aerosolization | | |
| 1d | 83 | 2 | 4 | 2 | No |
| 02 | 89 | 2 | 2 | 0 | No |
| 03 | 81 | 2 | 1 | -1 | No |
| 04 | 62 | 6 | 5 | -1 | Yes |
| 05 | 89 | 2 | 5 | 3 | No |
| 06 | 51 | 7 | 5 | -2 | Yes |
| 07 | 75 | 2 | 3 | 1 | No |
| 10 | 83 | 5 | 5 | 0 | No |
| 11 | 24 | 6 | 5 | -1 | Yes |
| 14 | 55 | 6 | 6 | 0 | Yes |
| 15 | 69 | 4 | 5 | 1 | Yes |
| 17 | 85 | 5 | 6 | 1 | No |
| 18 | 99 | 9 | 5 | -4 | No |
| 20 | 97 | 2 | 1 | -1 | No |
| SR03 | 43 | 3 | 11 | 8 | Yes |
| SR04 | 52 | 5 | 3 | -2 | Yes |
| SR13 | 83 | 4 | 6 | 2 | No |
| S2A4 | 84 | 7 | 11 | 4 | No |
| S2G4 | 91 | 3 | 6 | 3 | No |
| MRed03 | 96 | 3 | 2 | -1 | No |
| MRed04 | 59 | 4 | 4 | 0 | Yes |
| MRed07 | 90 | 10 | 3 | -7 | No |
| MRed11 | 89 | 4 | 5 | 1 | No |
| MRed18 | 96 | 3 | 10 | 7 | No |

| | | | | | |
|---|---|---|---|---|---|
| MRed19 | 87 | 5 | 9 | 4 | No |
| MRed20 | 76 | 2 | 18 | 16 | No |
| MRed22 | 86 | 3 | 9 | 6 | No |
| MRed25 | 44 | 3 | 3 | 0 | Yes |
| VHH-72 | 78 | 1 | 14 | 13 | No |

[a]% recovery was determined as the proportion of a $V_H$H that remained monomer following aerosolization. Data were used to construct **Fig. 3E**;
[b]% soluble aggregate was determined as the proportion of a $V_H$H that gave elution volume(s) smaller than that for the monomeric $V_H$H fraction;
[c]ΔSoluble agg. = "Post-aerosolization" – "Pre-aerosolization".

**Table S7.** $V_HH$ binding data obtained by tandem SPR SVNAs against surface-immobilized SARS-CoV-2 S

| Summary orientation #1: $V_HH$ followed by $V_HH$ + ACE2 | | | | | | | Summary orientation #2: ACE2 followed by ACE2 + $V_HH$ | | | | | | |
|---|---|---|---|---|---|---|---|---|---|---|---|---|---|
| **Cycle** | **Solution 1** | **Solution 2** | **End 1st inj[a] (RU)** | **End 2nd inj[a] (RU)** | **inj[a] 2-1 (ΔRU)** | **Blocker** | **Cycle** | **Solution 1** | **Solution 2** | **End 1st inj[a] (RU)** | **End 2nd inj[a] (RU)** | **inj[a] 2-1 (ΔRU)** | **Blocker** |
| 1 | Buffer | ACE2 | -3.1 | 56 | 59.1 | **No** | 1 | - | - | - | - | - | **-** |
| 2 | 1d | 1d+ACE2 | 26.3 | 40 | 13.7 | **Yes** | 2 | ACE2 | 1d+ACE2 | 57.1 | 56.4 | -0.7 | **Yes** |
| 3 | 02 | 02+ACE2 | 33 | 39.3 | 6.3 | **Yes** | 3 | ACE2 | 02+ACE2 | 55.8 | 59.6 | 3.8 | **Yes** |
| 4 | 03 | 03+ACE2 | 18.1 | 82.3 | 64.2 | **No** | 4 | ACE2 | 03+ACE2 | 52.8 | 75.3 | 22.5 | **No** |
| 5 | 04 | 04+ACE2 | 41.4 | 95.8 | 54.4 | **No** | 5 | ACE2 | 04+ACE2 | 52.4 | 98.8 | 46.4 | **No** |
| 6 | 05 | 05+ACE2 | 36.9 | 46.8 | 9.9 | **Yes** | 6 | ACE2 | 05+ACE2 | 52.2 | 60.5 | 8.3 | **Yes** |
| 7 | 06 | 06+ACE2 | 38.5 | 92.2 | 53.7 | **No** | 7 | ACE2 | 06+ACE2 | 52.3 | 94.4 | 42.1 | **No** |
| 8 | 07 | 07+ACE2 | 19.1 | 37.5 | 18.4 | **Yes** | 8 | ACE2 | 07+ACE2 | 52 | 59.1 | 7.1 | **Yes** |
| 10 | 10 | 10+ACE2 | 9.6 | 58.8 | 49.2 | **No** | 10 | ACE2 | 10+ACE2 | 52.1 | 64.1 | 12 | **No** |
| 11 | 11 | 11+ACE2 | 21.8 | 80.7 | 58.9 | **No** | 11 | ACE2 | 11+ACE2 | 52.3 | 83.6 | 31.3 | **No** |
| 12 | 14 | 14+ACE2 | 26.9 | 70.4 | 43.5 | **+/-** | 13 | ACE2 | 14+ACE2 | 52.3 | 75.9 | 23.6 | **No** |
| 13 | 15 | 15+ACE2 | 10.1 | 50.7 | 40.6 | **+/-** | 14 | ACE2 | 15+ACE2 | 50 | 59.7 | 9.7 | **No** |
| 14 | 17 | 17+ACE2 | 16.7 | 69.5 | 52.8 | **No** | 16 | ACE2 | 17+ACE2 | 52 | 71.7 | 19.7 | **No** |
| 15 | 18 | 18+ACE2 | 12.6 | 52.5 | 39.9 | **+/-** | 17 | ACE2 | 18+ACE2 | 50.9 | 61.7 | 10.8 | **+/-** |
| 16 | 20 | 20+ACE2 | 27.1 | 60.5 | 33.4 | **+/-** | 18 | ACE2 | 20+ACE2 | 51.3 | 66.2 | 14.9 | **+/-** |
| 17 | SR01 | SR01+ACE2 | 39.4 | 92 | 52.6 | **No** | 19 | ACE2 | SR01+ACE2 | 51.4 | 97.2 | 45.8 | **No** |
| 18 | SR02 | SR02+ACE2 | 10.3 | 60.2 | 49.9 | **No** | 20 | ACE2 | SR02+ACE2 | 49.1 | 67.8 | 18.7 | **No** |
| 19 | SR03 | SR03+ACE2 | 18.5 | 70.7 | 52.2 | **No** | 21 | ACE2 | SR03+ACE2 | 50 | 76.4 | 26.4 | **No** |
| 20 | SR04 | SR04+ACE2 | 10.9 | 63.6 | 52.7 | **No** | 22 | ACE2 | SR04+ACE2 | 50.5 | 68.5 | 18 | **No** |
| 21 | SR13 | SR13+ACE2 | 36.9 | 86.8 | 49.9 | **No** | 12 | ACE2 | SR13+ACE2 | 52.3 | 92.8 | 40.5 | **No** |
| 22 | SR16 | SR16+ACE2 | 37.4 | 88.8 | 51.4 | **No** | 15 | ACE2 | SR16+ACE2 | 52.1 | 95.7 | 43.6 | **No** |
| 23 | S2A3 | S2A3+ACE2 | 15 | 82 | 67 | **No** | 23 | ACE2 | S2A3+ACE2 | 50.8 | 72.2 | 21.4 | **No** |
| 24 | S2A4 | S2A4+ACE2 | 11.6 | 73.3 | 61.7 | **No** | 24 | ACE2 | S2A4+ACE2 | 50.9 | 71.1 | 20.2 | **No** |
| 25 | S2F3 | S2F3+ACE2 | 50 | 111.1 | 61.1 | **No** | 25 | ACE2 | S2F3+ACE2 | 50.7 | 106.9 | 56.2 | **No** |
| 26 | S2G3 | S2G3+ACE2 | 59 | 113.2 | 54.2 | **No** | 26 | ACE2 | S2G3+ACE2 | 51 | 114.1 | 63.1 | **No** |
| 27 | S2G4 | S2G4+ACE2 | 21.3 | 81.7 | 60.4 | **No** | 27 | ACE2 | S2G4+ACE2 | 50.9 | 79.3 | 28.4 | **No** |

| | | | | | | | | | | | | | |
|---|---|---|---|---|---|---|---|---|---|---|---|---|---|
| 28 | MRed04 | MRed04+ACE2 | 31.9 | 73.3 | 41.1 | **No** | 29 | ACE2 | MRed04+ACE2 | 62.6 | 73.3 | 10.7 | **+/-** |
| 29 | VHH-72 | VHH-72+ACE2 | 25.8 | 37.8 | 12 | **Yes** | 28 | ACE2 | VHH-72+ACE2 | 51 | 51.2 | 0.2 | **Yes** |

[a]inj, injection. $V_H$Hs were used at 20 – 40x $K_D$ concentrations, ACE2 (ACE2-$H_6$) used at 1 μM.

**Table S8.** Flow cytometry SVNAs against SARS-CoV-2 variants and SARS-CoV

| $V_HH$-Fc | SVNA $IC_{50}$ (nM) | | | | | | |
|---|---|---|---|---|---|---|---|
| | SARS-CoV-2 S | | | | | | SARS-CoV S |
| | Wuhan | Alpha | Beta | Gamma | Delta | Kappa | |
| **RBD-specific $V_HH$** | | | | | | | |
| 1d | 4.7 | 6.1 | 13.1 | 4.8 | 6.4 | 5.5 | - |
| 02 | 4.7 | 4.2 | - | - | 8.4 | 7.2 | - |
| 03 | - | - | - | - | - | - | - |
| 04 | 10.8 | 21.9 | - | - | - | - | - |
| 05 | 4.9 | 4.8 | - | - | 7.6 | 6.8 | - |
| 06 | - | - | - | - | - | - | - |
| 07 | 4.7 | 5.7 | 3.6 | 3.2 | 2.3 | 3.6 | 4.2 |
| 10 | 8.8 | 11.3 | 10.8 | 21.8 | - | - | - |
| 11 | 3.2 | 6.6 | 10.7 | 4.7 | 7.7 | 3.3 | 7.7 |
| 12 | 3.5 | 5.2 | 8 | 3.4 | 6.4 | 6.2 | 3.1 |
| 14 | 7.5 | 18 | 58 | 177 | - | - | - |
| 15 | 5.8 | 9.8 | 12.2 | 10.8 | - | - | - |
| 17 | 8.6 | 10.6 | 26.4 | 214 | - | - | - |
| 18 | 9.1 | 12.2 | 16.6 | 12.1 | 10.2 | 17.4 | 12.7 |
| 20 | 6.5 | 5.2 | 11.9 | 7.5 | 12.6 | 4.1 | 10.3 |
| MRed04 | 5 | 6.4 | 24.4 | 8.7 | 11.8 | 10.4 | 24.6 |
| MRed05 | 4.3 | 4.2 | 4.4 | 4.7 | 4.8 | 4.8 | - |
| **NTD-specific $V_HH$** | | | | | | | |
| SR01 | 4.2 | 3.1 | 8.8 | 2.1 | 3.1 | 2.3 | 5.1 |
| SR02 | 1.7 | 7.3 | - | 4.7 | 6.1 | 3.1 | - |
| SR13 | 7.7 | 22.4 | - | 16.5 | - | 12.2 | 15 |
| **Reference** | | | | | | | |
| VHH-72 | 5.6 | 10.6 | 5.1 | 3.3 | 10.5 | 8.5 | 7.8 |

Dash indicates lack of neutralization.

**Table S9**. Summary of HDX-MS experimental conditions and statistics

| Data Set | Batch 1 | Batch 2 | Batch 3 | Batch 4 | Batch 5 |
|---|---|---|---|---|---|
| **Target protein** | S1 | | | | S |
| **Protein states included** | 03,11, SR03, MRed03, MRed07 | 04,05,06,10,14,15 | 1d, 02,17,18, MRed04 | 07, 20, MRed05, SR01, SR02 | S2A3 |
| **Technical replicates** | 3 | | | | 4 |
| **HDX time course (min)** | 3 | | | | 1 |
| **HDX reaction details** | 10 mM Tris, pD = 7.0, 20 °C in 45% $D_2O$ | | | | |
| **Protease for on-line digestion** | Pepsin | | | | Nepenthesin-II |
| **LC Gradient** | 1-35% ACN, 8 min | | | | 2-40% ACN, 15 min |
| **# of peptides found across all samples** | 215 | 222 | 219 | 196 | 278 |
| **Sequence coverage** | 71 | 75 | 75 | 75 | 75 |
| **Average peptide length / Redundancy** | 15.1/4.9 | 13.7/4.5 | 13.7/4.5 | 13.7/4.5 | 12.2/2.8 |
| **Repeatability (average standard deviation)** | 0.8 | 0.9 | 0.9 | 0.9 | 1.2 |
| **3 x SD cut-off[a]** | 6 | 4.7 | 5 | 4.9 | 3.5 |
| **Significant differences in HDX** | Two-state student T-Test performed for each state (ΔD > 3 x SD, 1-p value > 0.98) | | | | |

[a]SD cutoff was calculated for each state within a batch, and the highest cutoff was applied to all states in that batch.
