## Supplemental Figures for "Arsenal of Nanobodies for Broad-Spectrum Countermeasures against Current and Future SARS-CoV-2 Variants of Concerns"

**Figure S1**

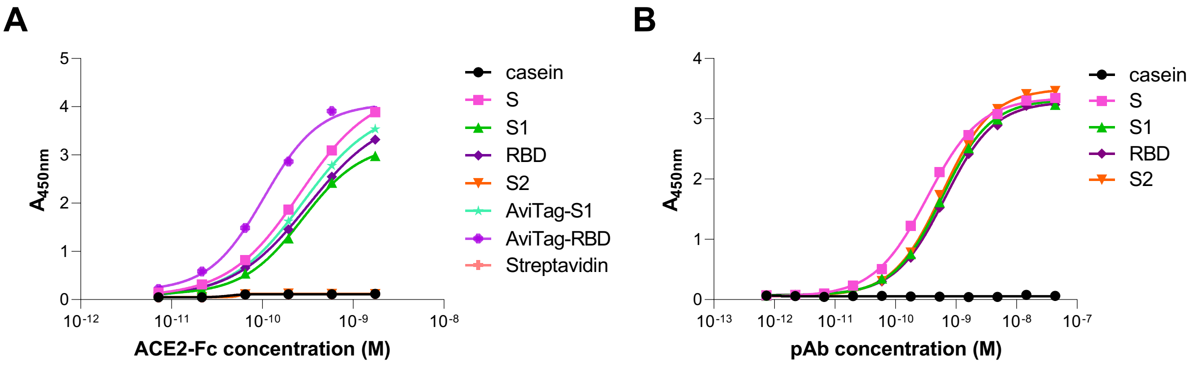


| Antigen | S | S1 | S2 | RBD | AviTag-RBD | AviTag-S1 |
|---|---|---|---|---|---|---|
| $EC_{50}$ (nM) | 0.26 | 0.28 | - | 0.32 | 0.1 | 0.28 |

| Antigen | S | S1 | S2 | RBD |
|---|---|---|---|---|
| $EC_{50}$ (nM) | 0.34 | 0.59 | 0.6 | 0.65 |

**Fig. S1. Antigen validation.** Prior to immunization, serology and panning experiments, SARS-CoV-2 spike protein antigens were validated for functionality in adsorbed/captured states on microtiter wells by measuring their binding to ACE2 and an anti-S polyclonal antibody. **A**) ELISA assessing the binding of microtiter well-adsorbed (S, S1, S2 and RBD) and microtiter well-captured (AviTag-S1, AviTag-RBD) SARS-CoV-2 spike glycoprotein fragments to their human receptor ACE2 (fused to human IgG1, ACE2-Fc). AviTag-S1 and AviTag-RBD were captured on streptavidin-coated microtiter wells through their C-terminal biotin tag. **B**) ELISA confirming the binding of microtiter well-adsorbed SARS-CoV-2 spike glycoprotein fragments S, S1, S2 and RBD to a commercial rabbit anti-SARS-CoV-2 S polyclonal antibody (pAb). No binding was observed to S2 (**A**) or to casein (**A** and **B**), demonstrating binding specificities. Calculated $EC_{50}$s (Half-maximal effective concentration) are tabulated below the graphs. SARS-CoV-2 Wuhan spike glycoprotein fragments were used in the assays.

#### Figure S2

**A**

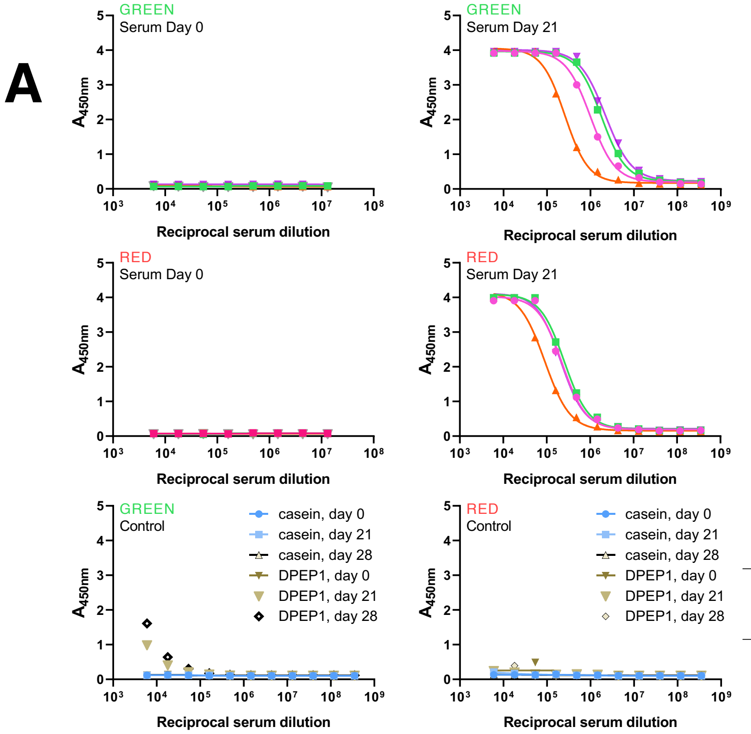


| Llama | Day | $EC_{50}$ ( ×$10^6$ reciprocal serum dilution) | | | |
|---|---|---|---|---|---|
| | | S | S1 | S2 | RBD |
| Green | 0 | - | - | - | - |
| | 21 | 1 | 1.8 | 0.3 | 2.2 |
| | 28 | 1.2 | 1.8 | 0.4 | 2 |
| Red | 0 | - | - | - | - |
| | 21 | 0.2 | 0.3 | 0.1 | 0.2 |
| | 28 | 0.3 | 0.3 | 0.2 | 0.3 |

**B**

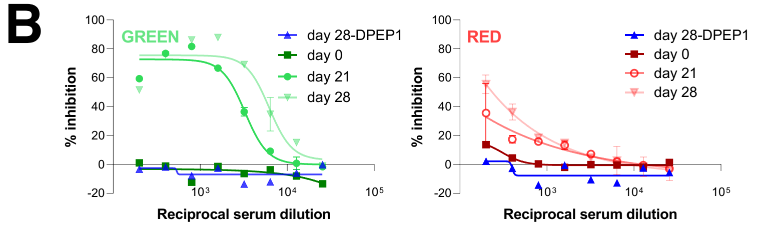


| Llama | $IC_{50}$ (reciprocal serum dilution) | |
|---|---|---|
| | Day 21 | Day 28 |
| Green | 3300 | 6200 |
| Red | <200 | <400 |

**Fig. S2. Llama serology. A)** ELISA performed with pre-immune (day 0) and immune (days 21 and 28) sera demonstrated SARS-CoV-2 spike glycoprotein-immunized llamas Green and Red generated strong immune responses towards S, S1, S2 and RBD. ELISA performed with day 0, 21 and 28 sera showed immunized llama sera did not react with non-target antigens (casein and dipeptidase 1 [DPEP1]), demonstrating specificity of the immune response. $EC_{50}$, measures of the strength of immune response, are shown in the table. Based on $EC_{50}$ values, Green generated a stronger immune response in comparison to Red, up to 10-fold, consistently across all four spike protein fragments. **B)** Flow cytometry surrogate virus neutralization assays (SVNAs) performed with pre-immune (day 0) and immune (days 21 and 28) sera demonstrated llama Green mounted a polyclonal immune response that was more potent in blocking the binding of SARS-CoV-2 S to Vero E6 cell surface-associated ACE2 in comparison to llama Red. $IC_{50}$s (Half-maximal inhibitory concentrations), measures of the strength of serum neutralization capability, are shown in the table. Due to a lack of complete curves, $IC_{50}$s for llama Red sera were estimated by assuming similar upper plateaus as those for llama Green sera. $IC_{50}$s of 3300 (day 21) and 6200 (day 28) reciprocal serum dilution (RSD) were obtained in the case of the Green sera compared to far weaker ones, <200 RSD (day 21) and <400 RSD (day 28), for the Red sera. SARS-CoV-2 Wuhan spike glycoprotein fragments were used for immunizations and in assays.

**Figure S3**

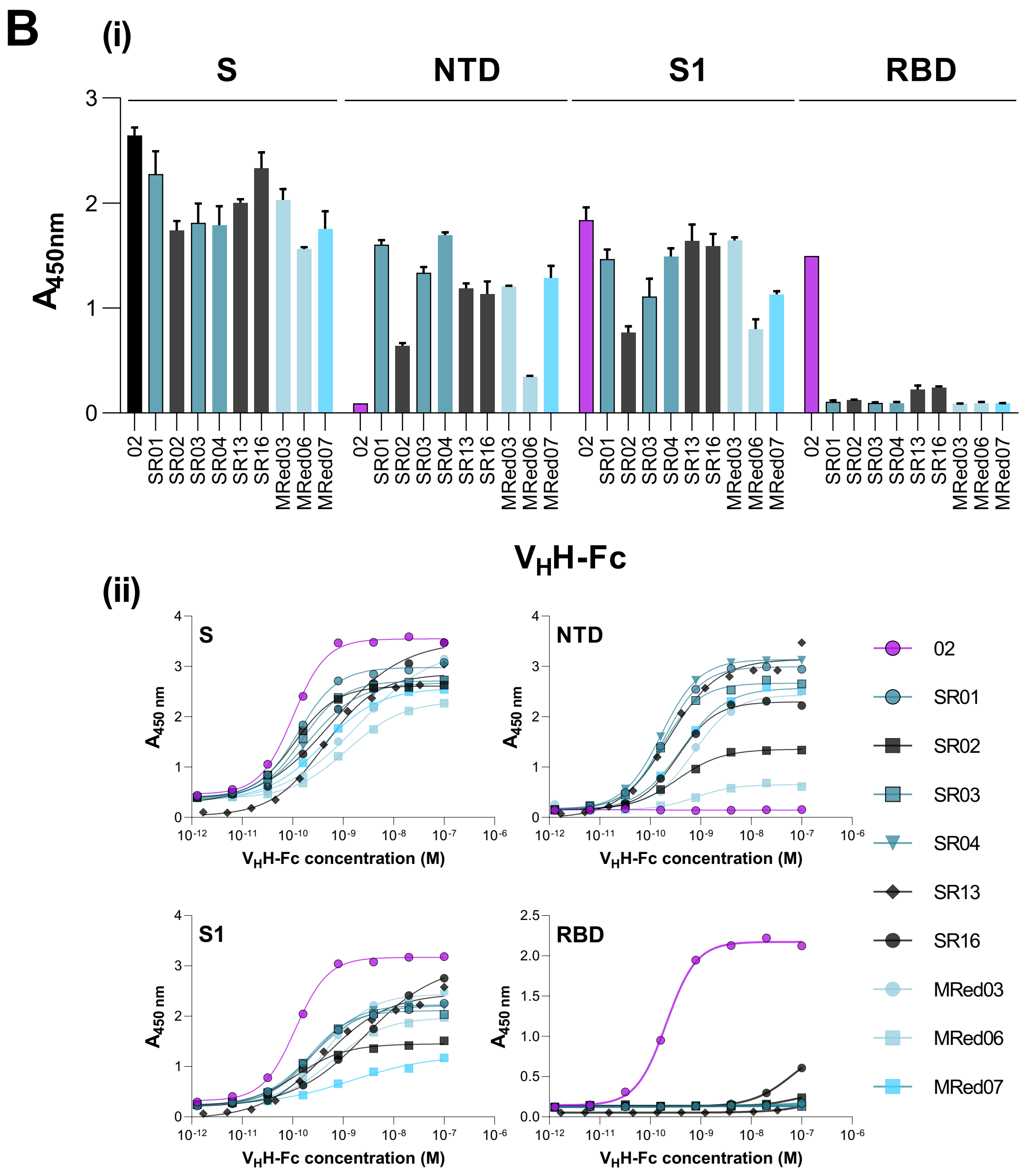


**Fig. S3. Binding affinity and subunit/domain specificity of SARS-CoV-2 $V_H$Hs and $V_H$H-Fcs by SPR and ELISA. A**) Representative SPR sensorgrams showing single-cycle kinetics analysis of 02, SR03 and S2A4 $V_H$H binding to SARS-CoV-2 S, S1, S2 and RBD. Spike glycoprotein fragments were immobilized on CM5 sensorchip surfaces and $V_H$Hs flowed over at concentration ranges described in the figure. Black lines represent raw data points, red lines are fits to a 1:1 binding model. The 02, SR03 and S2A4 antibodies represent SPR binding profiles for $V_H$Hs specific to RBD, NTD and S2, respectively. Calculated kinetic and equilibrium dissociation constants ($k_a$s, $k_d$s and $K_D$s) obtained for binding of $V_H$Hs against SARS-CoV-2 spike glycoprotein fragments are reported in **Table S2**. **B**) ELISA assessing the domain specificity of SARS-CoV-2 $V_H$H-Fcs. Assays were performed against SARS-CoV-2 S, S1, NTD and RBD at a fixed (13 nM) (**i**) or varying (**ii**) concentrations of $V_H$H-Fcs. 02 is included as internal control. (**ii**) Apparent $EC_{50}$s ($EC_{50}$apps) obtained from graphs are recorded in **Table S3**. SARS-CoV-2 Wuhan spike glycoprotein fragments were used in assays.

#### Figure S4

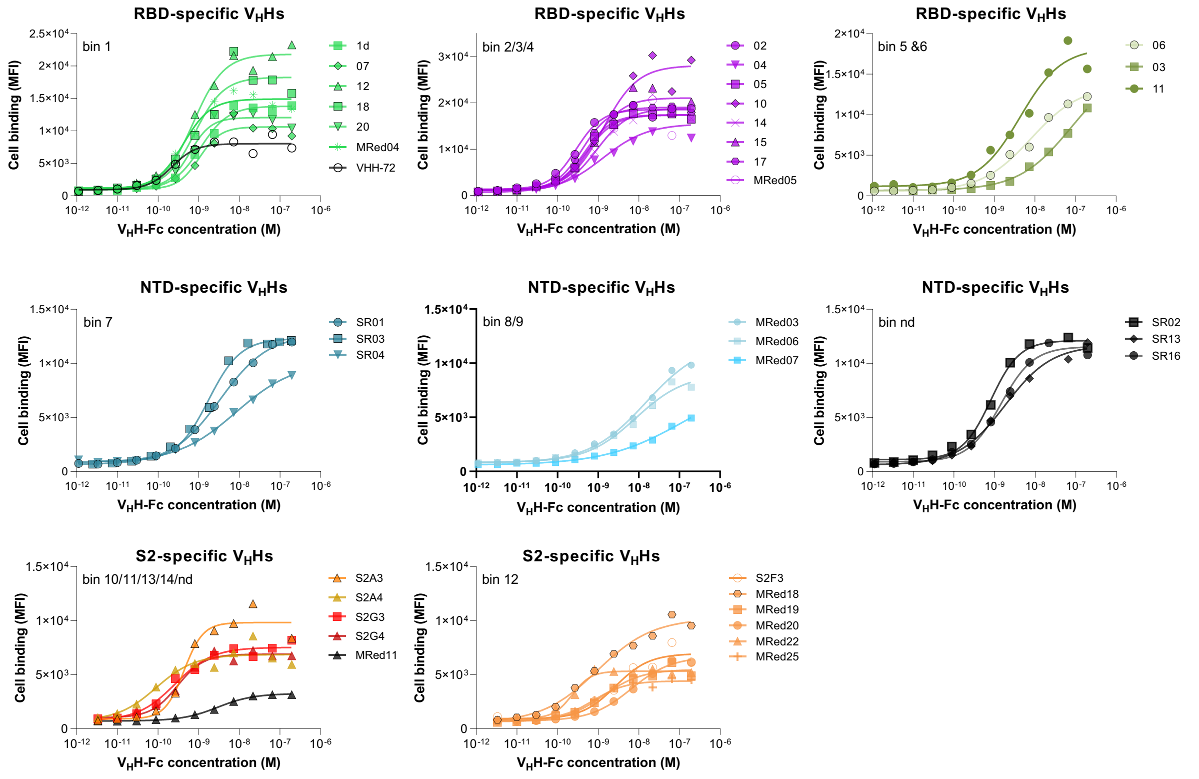


**Fig. S4. Assessing binding of SARS-CoV-2 $V_H$H-Fcs to S-expressing CHO cells (CHO-SPK) by flow cytometry.** Calculated apparent $EC_{50}$s ($EC_{50}$apps) obtained from graphs are reported in **Table 1** and used in **Fig. 1D**. None of the $V_H$H-Fcs bound to parental non-S-expressing CHO cells at the highest concentration tested (200 nM).

**Figure S5**

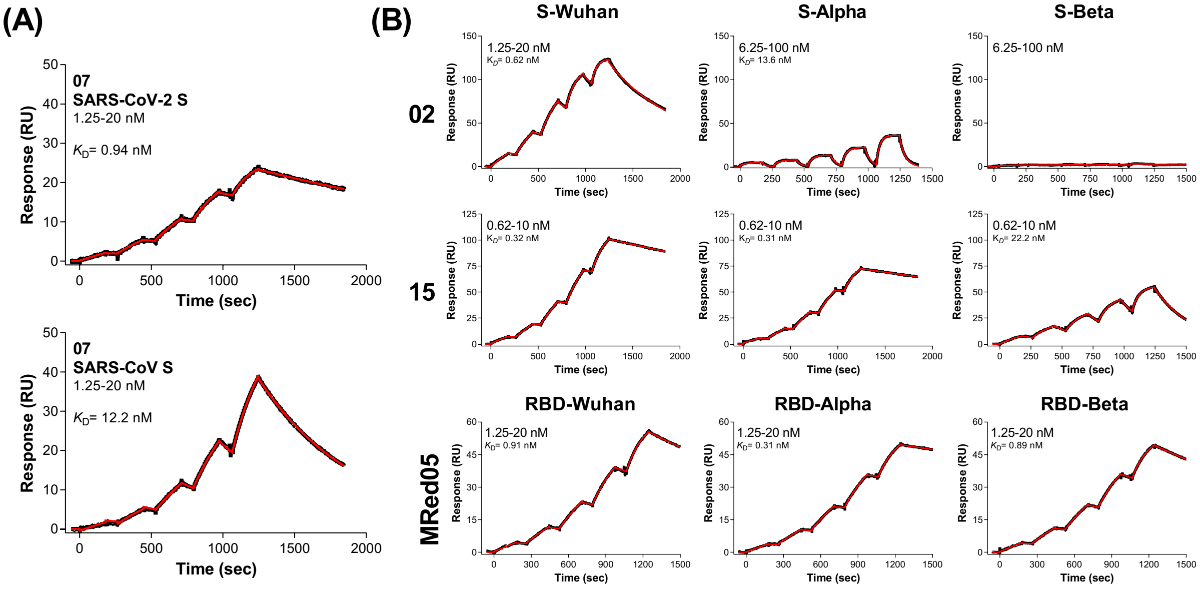


**Fig. S5. Cross-reactivity assessment of SARS-CoV-2 $V_H$Hs by SPR.** Representative SPR sensorgrams showing single-cycle kinetics analysis of 07 binding to SARS-CoV-2 S and SARS-CoV S **(A)**, and 02, 15 and MRed05 binding to Wuhan, Alpha and Beta S (02, 15) and RBD (MRed05) **(B)**. Spike glycoproteins (S) were immobilized on CM5 sensorchip surfaces followed by flowing 07, 02 and 15 $V_H$Hs at concentration ranges described in the figure. MRed05 $V_H$H-Fc was captured on anti-human Fc sensorchip surfaces followed by flowing RBDs at concentration ranges described in the figure. Black lines represent raw data points, red lines are 1:1 model fits. Calculated kinetic and equilibrium dissociation constants ($k_a$s, $k_d$s and $K_D$s) obtained for binding of $V_H$Hs/$V_H$H-Fc to spike glycoprotein fragments are reported in **Fig. 2B**, **Fig. 2C**, **Table 1 and Table S4**.

#### Figure S6

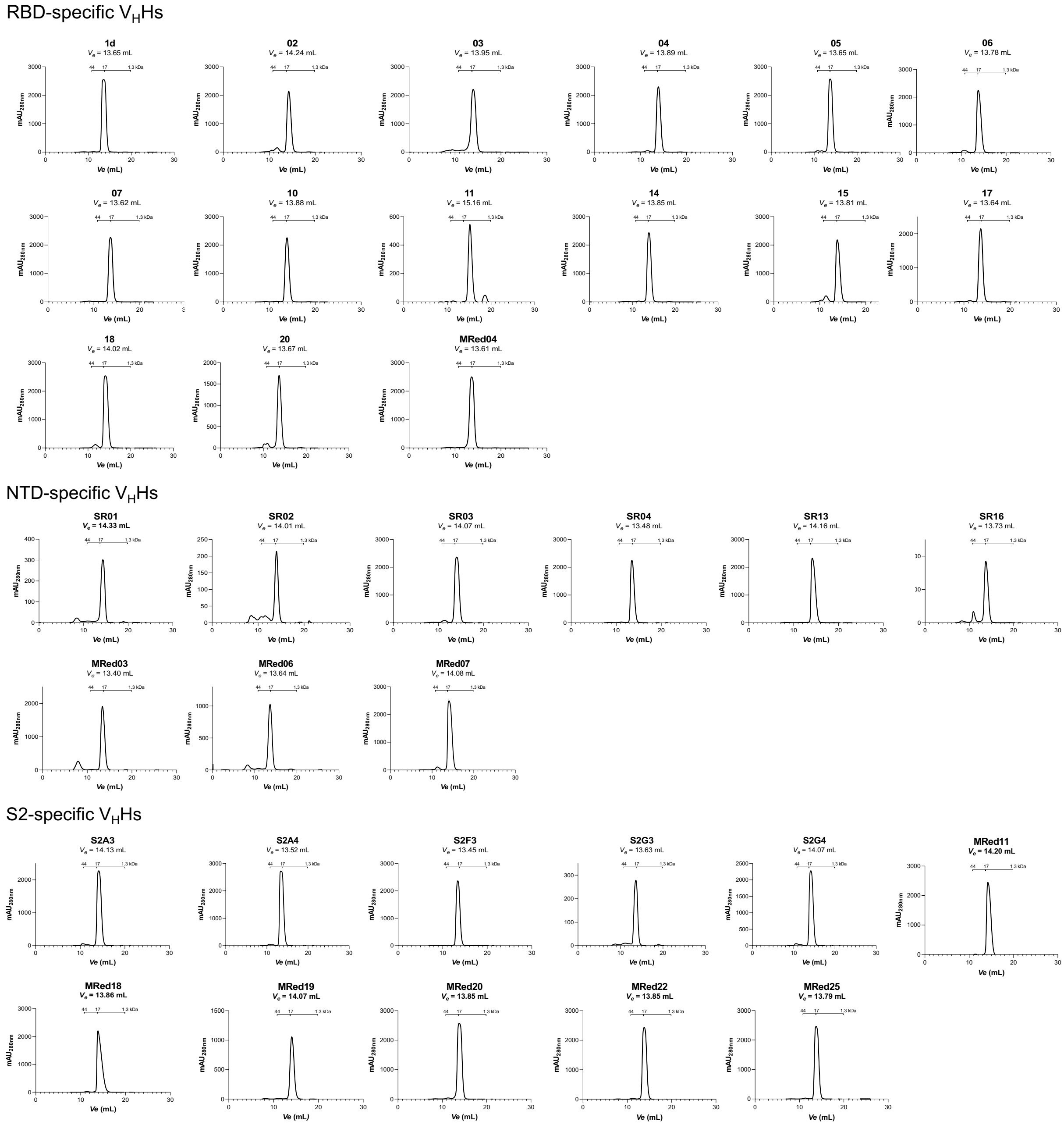


**Fig. S6. Aggregation propensity of SARS-CoV-2 $V_H$Hs.** SEC profiles of SARS-CoV-2 $V_H$Hs demonstrated $V_H$Hs were aggregation resistant and had elution volumes ($V_e$s) consistent with their monomeric states. The $V_e$ positions of molecular mass standards (44 kDa, 17 kDa, 1.3 kDa) are shown. SEC was performed using a Superdex® 75 Increase column. $V_H$Hs 12 and MRed05 were not tested due to insufficient quantities. mAU, milliabsorbance unit.

**Figure S7**

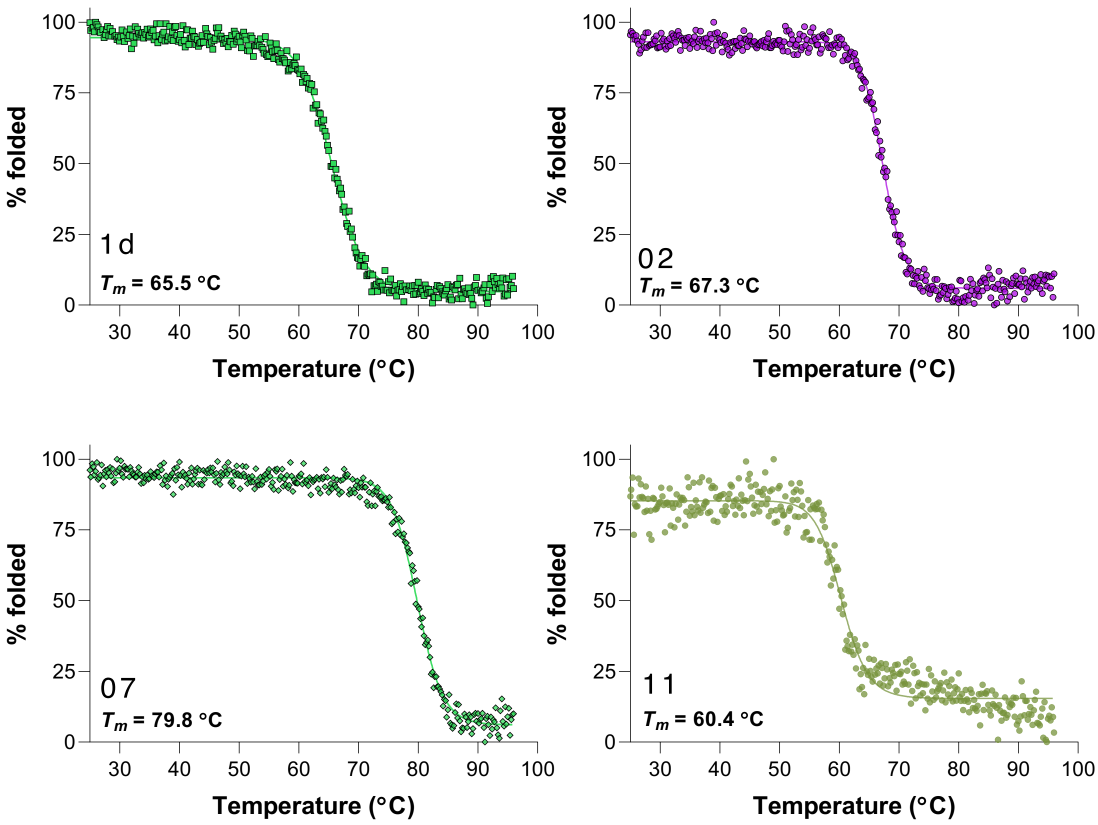


**Fig. S7. Temperature-induced unfolding profiles of SARS-CoV-2 $V_H$Hs.** Representative examples showing the thermal unfolding of 1d, 02, 07 and 11 determined using circular dichroism (CD) spectroscopy. $V_H$H thermal unfolding midpoint temperatures ($T_m$s), obtained from graphs of % folded *vs* temperature, were used to construct **Fig. 3B and Table S5**.

**Figure S8**

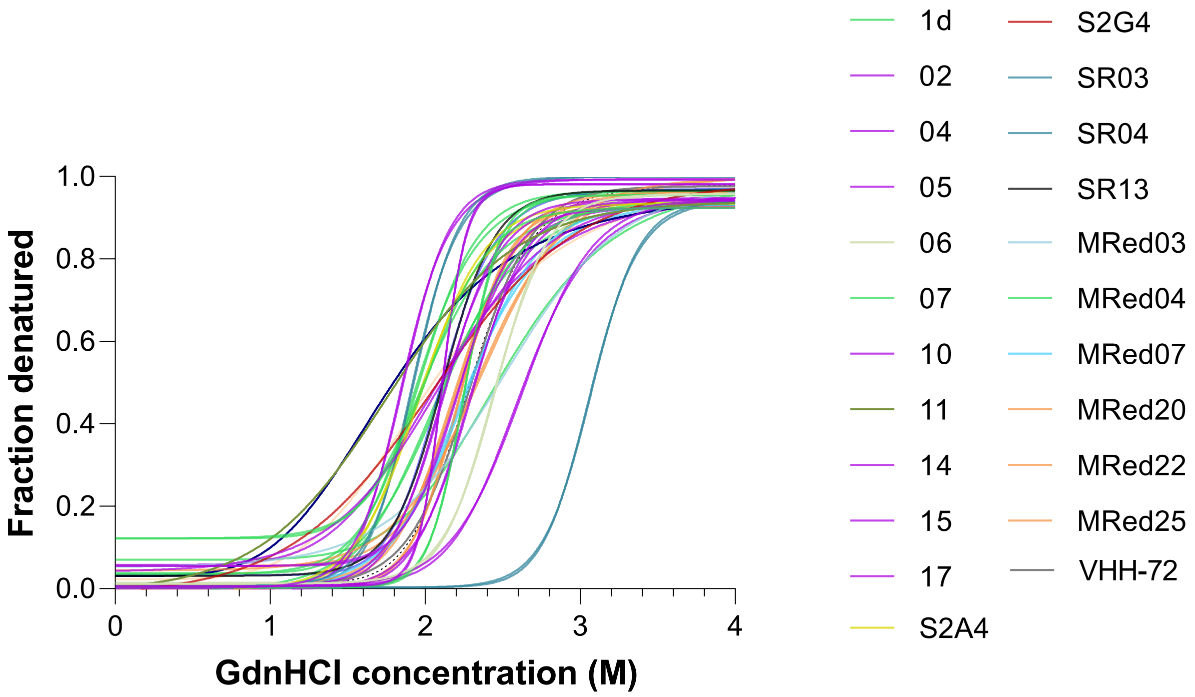


**Fig. S8. GdnHCl-induced unfolding profiles of SARS-CoV-2 $V_H$Hs.** Unfolding of $V_H$Hs was monitored using the automated Hunky system as described in Methods. Plots of fraction denatured *vs* denaturant (GdnHCl) concentration confirm the two-state transition model. The midpoints of the unfolding transition, $C_m$s, are reported in **Table S5**.

**Figure S9**

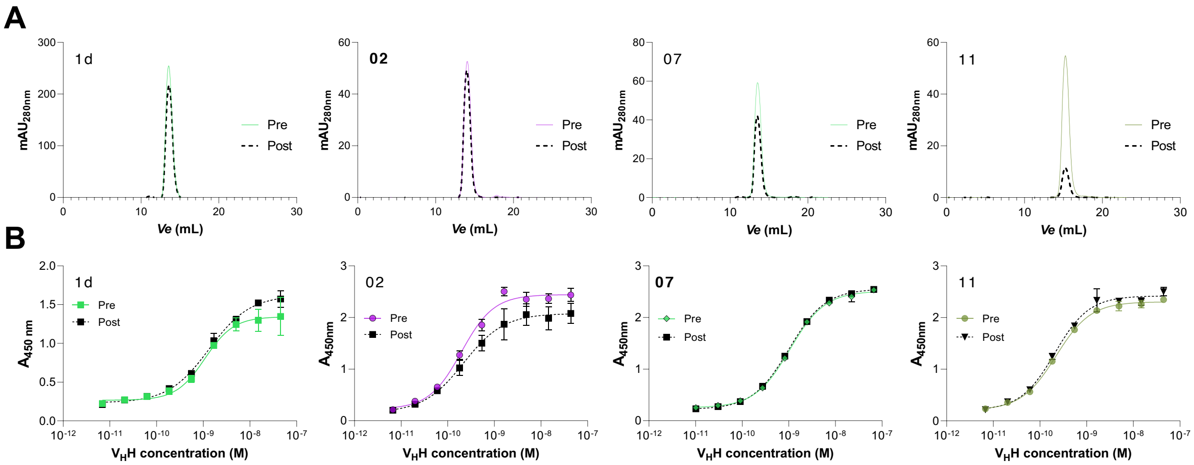


**Fig. S9. Stability of SARS-CoV-2 V$_H$Hs against aerosolization. A**) Assessing the effect of aerosolization on the aggregation resistance of V$_H$Hs by comparing the SEC profiles of pre- *vs* post-aerosolized V$_H$Hs. Results for representative V$_H$Hs are shown. 1d, 02 and 07 represent the majority of V$_H$Hs which were resistant to aerosolization-induced aggregation, showing homogenous monomeric peaks following aerosolization. V$_H$H 11 represents one of the few V$_H$Hs which formed visible, precipitating aggregates reflected by a significant reduction of their monomeric peak area (compare monomeric peak for pre- *vs* post-aerosolized V$_H$H 11). Aggregation data for a complete set of V$_H$Hs are summarized in **Table S6**. **B**) ELISA assessing the effect of aerosolization on the functionality of V$_H$Hs by comparing the binding activity of pre- *vs* post-aerosolized V$_H$Hs against SARS-CoV-2 S. Essentially identical $EC_{50}$s for pre- *vs* post-aerosolized V$_H$Hs indicates aerosolization had no effect on the functional activity of V$_H$Hs. $EC_{50}$s are summarized in **Fig. 3E**. Pre, pre-aerosolized V$_H$H; post, post-aerosolized V$_H$H. mAU, milliabsorbance; $V_e$, elution volumes.

**Figure S10**

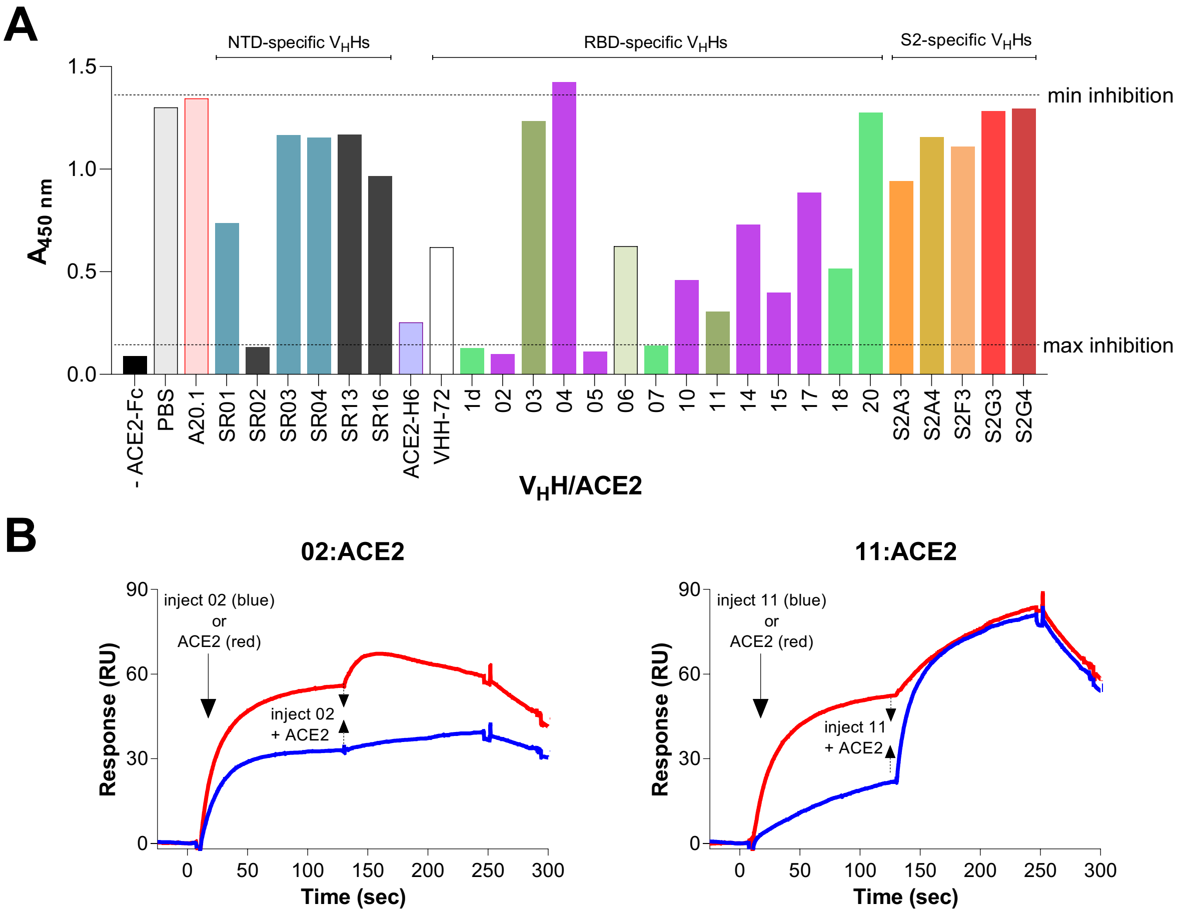


**Fig. S10. $V_HH$ SVNAs by ELISA and SPR. A)** ELISA assessing the ability of monomeric $V_HHs$ to block the binding of ACE2 to SARS-CoV-2 S. A dimeric, Fc-fused ACE2 (ACE2-Fc) was used in the assays at 100 ng and mixed with a fixed concentration of $V_HHs$ (1 µM). VHH-72 $V_HH$[1] and monomeric ACE2-$H_6$ served as positive and reference controls, while toxin A-specific A20.1 $V_HH$ [2] was a negative control. "PBS" and "A20.1" represent assays in which a SARS-CoV-2 $V_HH$ was substituted with PBS or A20.1 $V_HH$ and provide reference binding signals for lack of blocking ("min inhibition"). "-ACE2-Fc" control represent assays with ACE2-Fc omitted and provides a reference binding signal for 100% blocking ("max inhibition"). $V_HHs$ are color-coded according to their epitope bin designation (*see* **Fig. 5A**). **B)** Representative sensorgrams assessing the ability of monomeric $V_HHs$ in blocking the binding of ACE2 receptor to its ligand SARS-CoV-2 S. SPR assays were performed in two orientations, where injection of $V_HH$ (orientation #1) or ACE2 (orientation #2) at 20 – 40× $K_D$ concentration ($V_HH$) or 1 µM (ACE2) over immobilized S, was followed by injection of a mixture of $V_HH$ + ACE2 at the same $V_HH$ and ACE2 concentrations. Blue and red profiles represent binding results with the two orientations. "02:ACE2" represents a profile for a blocking $V_HH$, where the addition of the $V_HH$ or ACE2 results in no significant increase in binding over that achieved by the injection of the ACE2 or $V_HH$ on the antigen surface. "11:ACE2" represents a profile for non-blocking $V_HH$, where the addition of the $V_HH$ or ACE2 results in significant increase in binding over that achieved by the injection of the ACE2 or $V_HH$ on the antigen surface. ΔRUs, representing binding differences between the first and second injection, were calculated from the sensorgrams and used to identify $V_HHs$ that block the binding of ACE2 receptor to its ligand RBD (*see* **Table S7**). SARS-CoV-2 Wuhan S were used in assays. ACE2, monomeric ACE2-$H_6$.

#### Figure S11

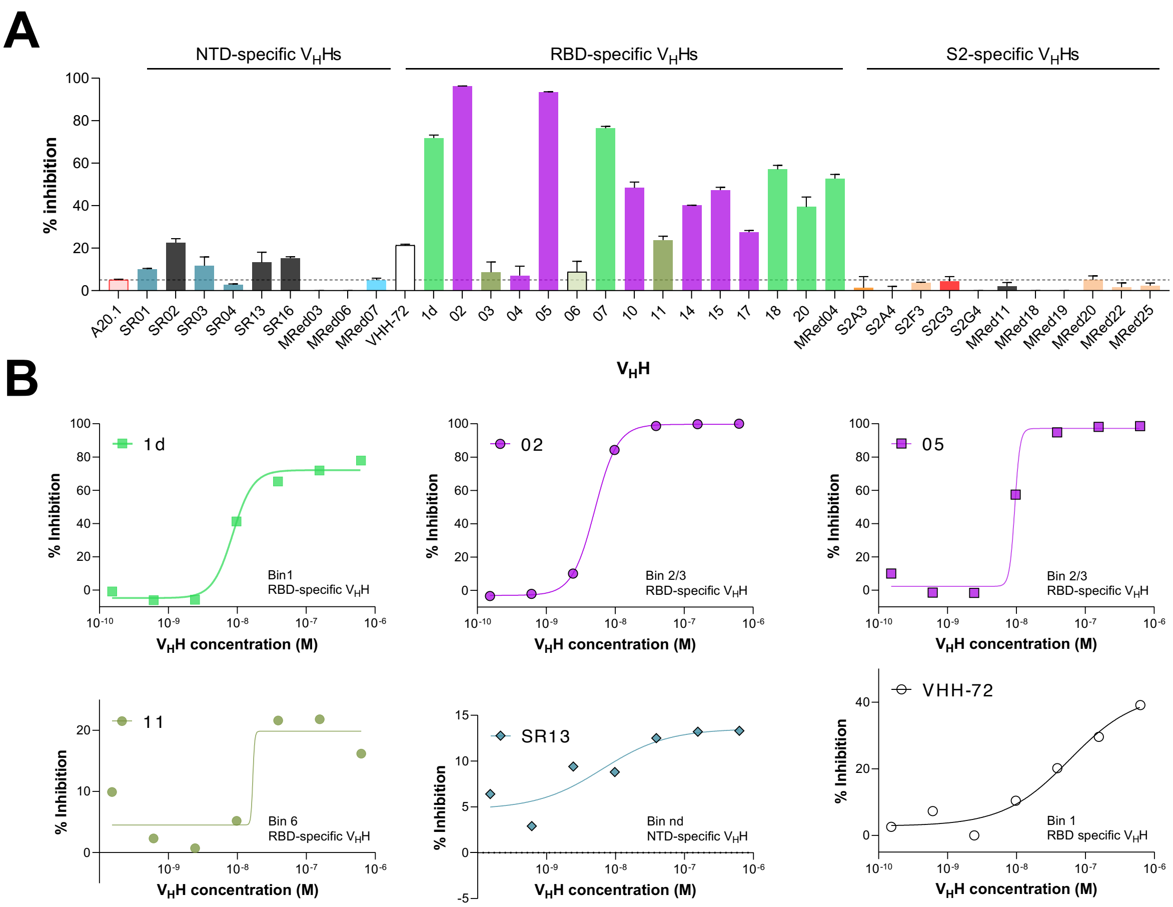


**Fig. S11. Flow cytometry SVNAs assessing the ability of monomeric SARS-CoV-2 $V_H$Hs in blocking the binding of SARS-CoV-2 S to ACE2-expressing Vero E6 cells at 100 nM (A) or varying (B) $V_H$H concentrations.** Representative plots are shown in "**B**". $V_H$Hs 1d, 02, 05 and 11 are RBD-specific, SR13 is NTD-specific. The complete list of $IC_{50}$s calculated from graphs exemplified in "**B**" are shown in **Table 2**. Monomeric "ACE2" (ACE2-$H_6$) serves as positive "antibody" control and reference; VHH-72 $V_H$H[1] is included as benchmark. "A20.1" and "PBS" represent negative control assays in which $V_H$Hs were replaced with *C. difficile* toxin A-specific A20.1 $V_H$H [2] and PBS, respectively. $V_H$Hs are color-coded according to their epitope bin designation (*see* **Fig. 5A**).

#### Figure S12

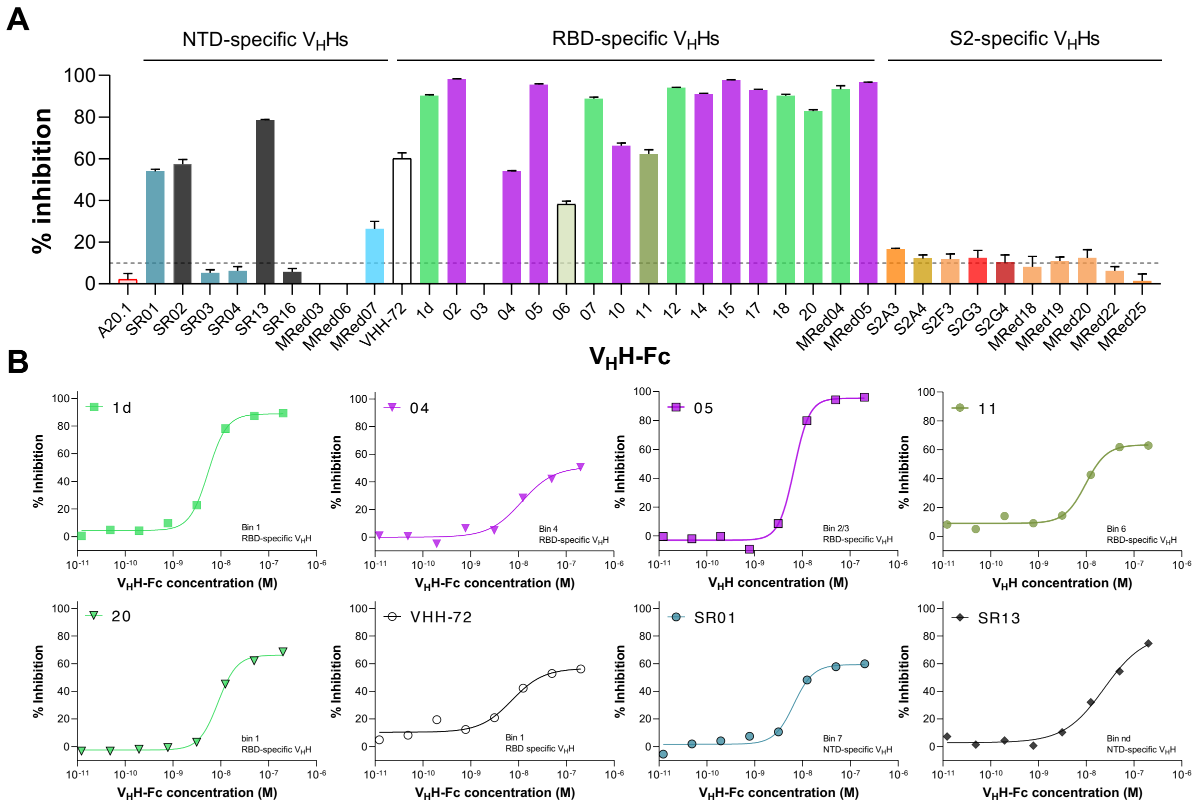


**Fig. S12. Flow cytometry SVNAs assessing the ability of bivalent SARS-CoV-2 $V_H$H-Fcs in blocking the binding of SARS-CoV-2 S to ACE2-expressing Vero E6 cells at 250 nM (A) or varying (B) $V_H$H-Fc concentrations.** Representative plots are shown in "**B**". 1d, 02, 04, 05, 11 and 20 $V_H$H-Fcs are RBD-specific, SR01 and SR13 are NTD-specific. The complete list of $IC_{50}$s calculated from graphs exemplified in "**B**" are reported in **Table 2**. VHH-72 $V_H$H-Fc[1] is included as benchmark. "A20.1" is a negative control in which $V_H$H-Fcs were replaced with *C. difficile* toxin A-specific A20.1 $V_H$H-Fc[2]. Biotinylated SARS-CoV-2 Wuhan S was used in assays. $V_H$H-Fcs are color-coded according to their epitope bin designation (*see* **Fig. 5A**).

### Figure S13

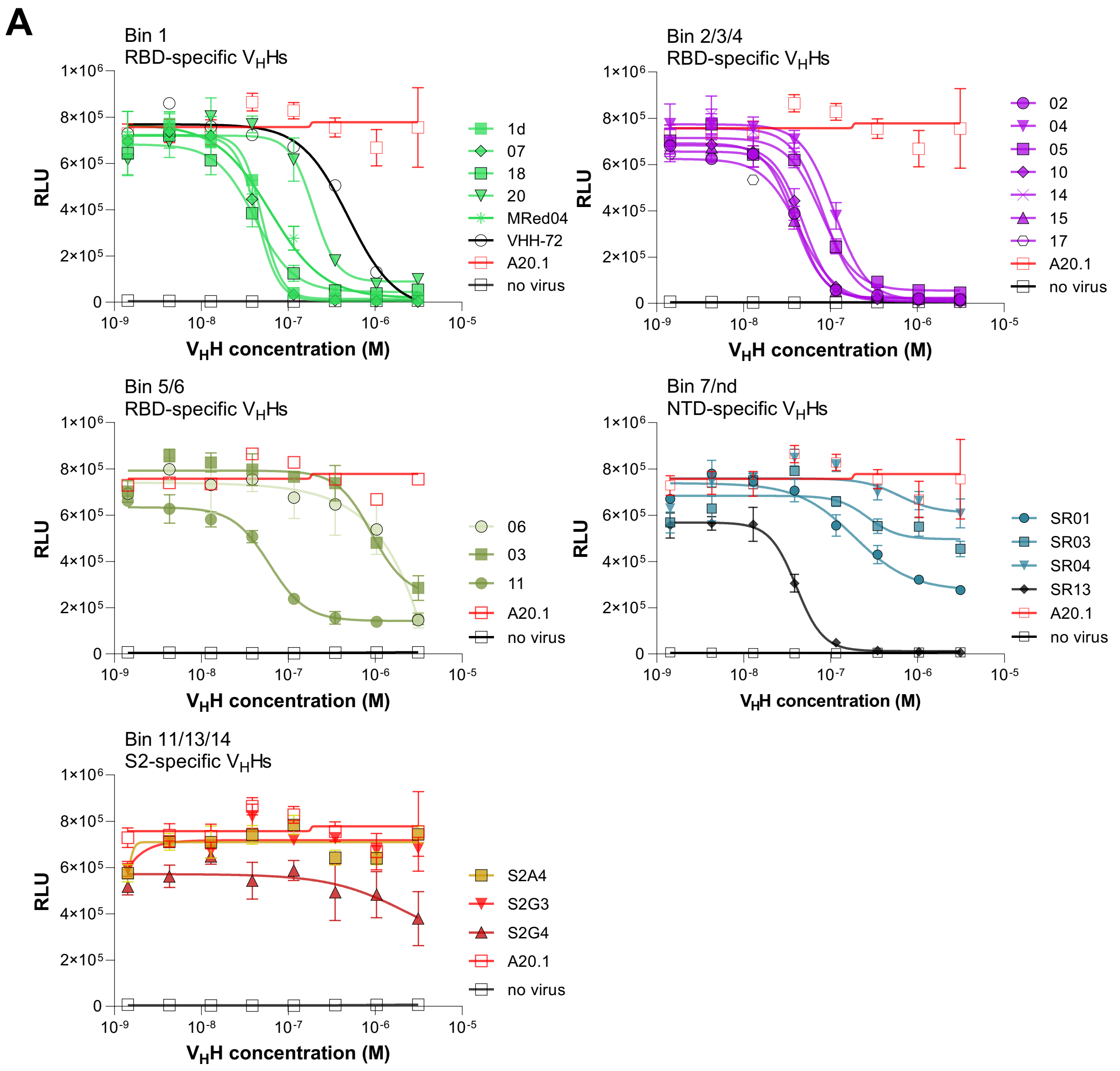
A
Bin 1
RBD-specific VHHs
1d
07
18
20
MRed04
VHH-72
A20.1
no virus
Bin 2/3/4
RBD-specific VHHs
02
04
05
10
14
15
17
A20.1
no virus
Bin 5/6
RBD-specific VHHs
06
03
11
A20.1
no virus
Bin 7/nd
NTD-specific VHHs
SR01
SR03
SR04
SR13
A20.1
no virus
Bin 11/13/14
S2-specific VHHs
S2A4
S2G3
S2G4
A20.1
no virus
RLU
VHH concentration (M)

**B**

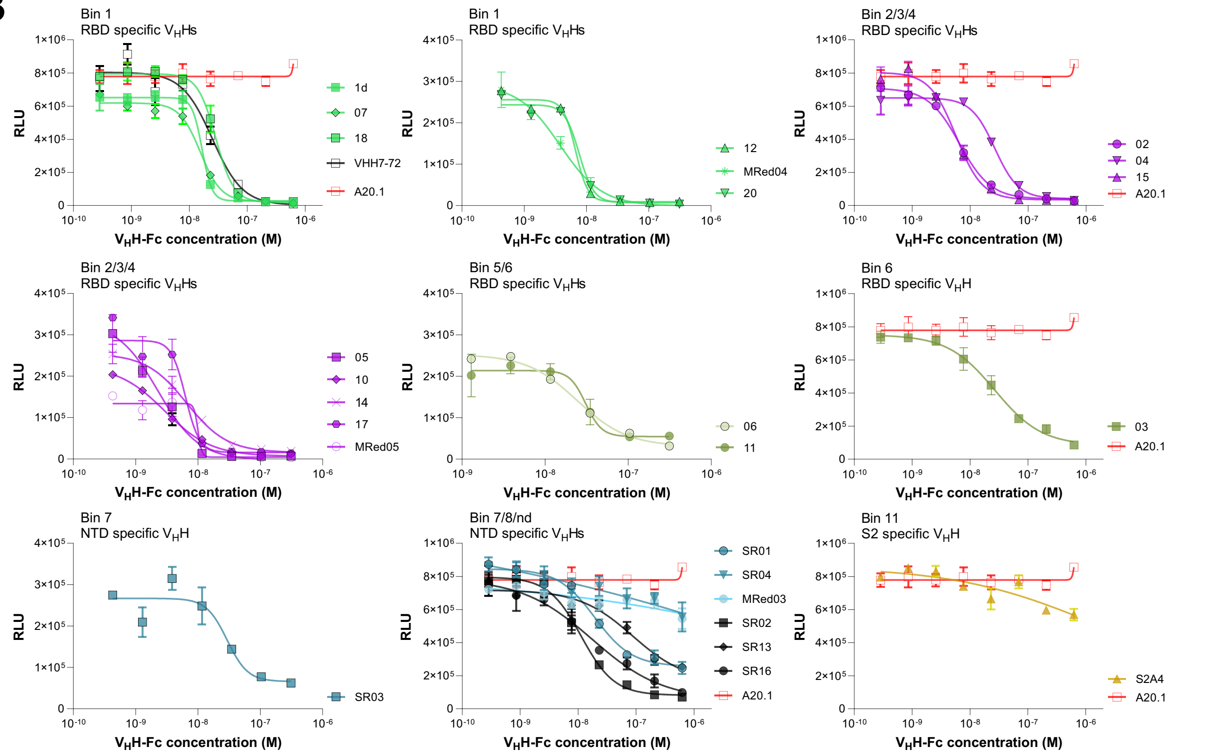


**Fig. S13. Pseudotyped virus neutralization assays (PVNAs) assessing the ability of SARS-CoV-2 $V_H$Hs (A) and $V_H$H-Fcs (B) in blocking the infection of ACE2-expressing HEK293T cells by spike-pseudotyped lentiviruses.** Results, obtained from assays performed at multiple antibody concentrations, are shown only for those $V_H$Hs/$V_H$H-Fcs that were positive at a threshold inhibition of 75% in preliminary inhibition screening assays. The complete list of $IC_{50}$s calculated from graphs are recorded in **Table 2**. VHH-72 $V_H$H-Fc[1] is included as benchmark. “A20.1” represent negative antibody control assays in which SARS-CoV-2 $V_H$Hs/$V_H$H-Fcs were replaced with *C. difficile* toxin A-specific A20.1 $V_H$H/$V_H$H-Fc[2], and provide upper limit infection RLU (relative light unit). “no virus” represent control assays in which pseudo-typed virus is omitted, and provide lower limit infection RLU. $V_H$Hs/$V_H$H-Fcs are color-coded according to their epitope bin designation (*see* **Fig. 5A**). Data are mean ± s.e.m. of technical triplicates.

Figure S14

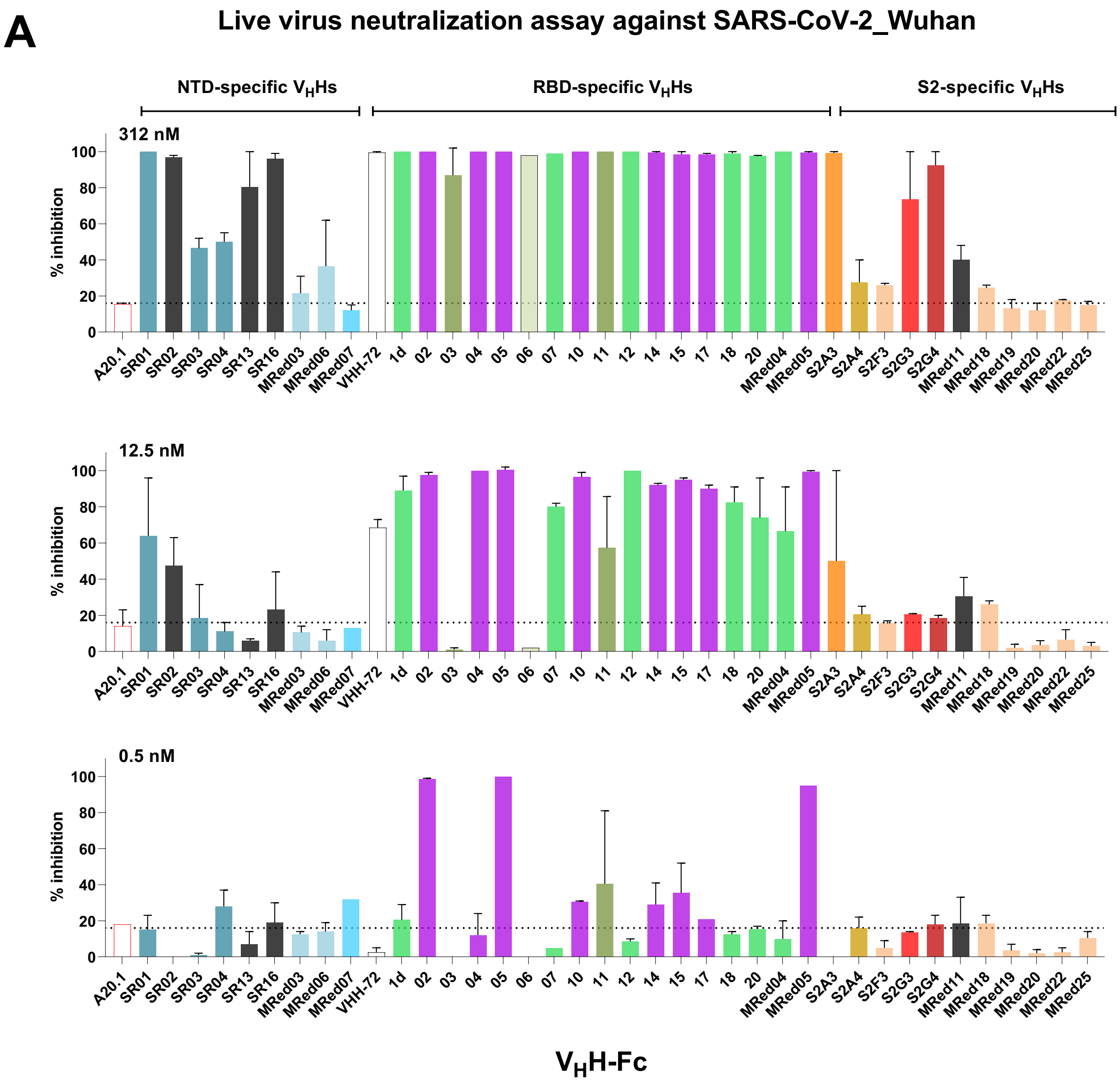
A
Live virus neutralization assay against SARS-CoV-2_Wuhan
NTD-specific VHHs
RBD-specific VHHs
S2-specific VHHs
312 nM
12.5 nM
0.5 nM
% inhibition
0
20
40
60
80
100
A20.1
SR01
SR02
SR03
SR04
SR13
SR16
MRed03
MRed06
MRed07
VHH-72
1d
02
03
04
05
06
07
10
11
12
14
15
17
18
20
MRed04
MRed05
S2A3
S2A4
S2F3
S2G3
S2G4
MRed11
MRed18
MRed19
MRed20
MRed22
MRed25
VHH-Fc

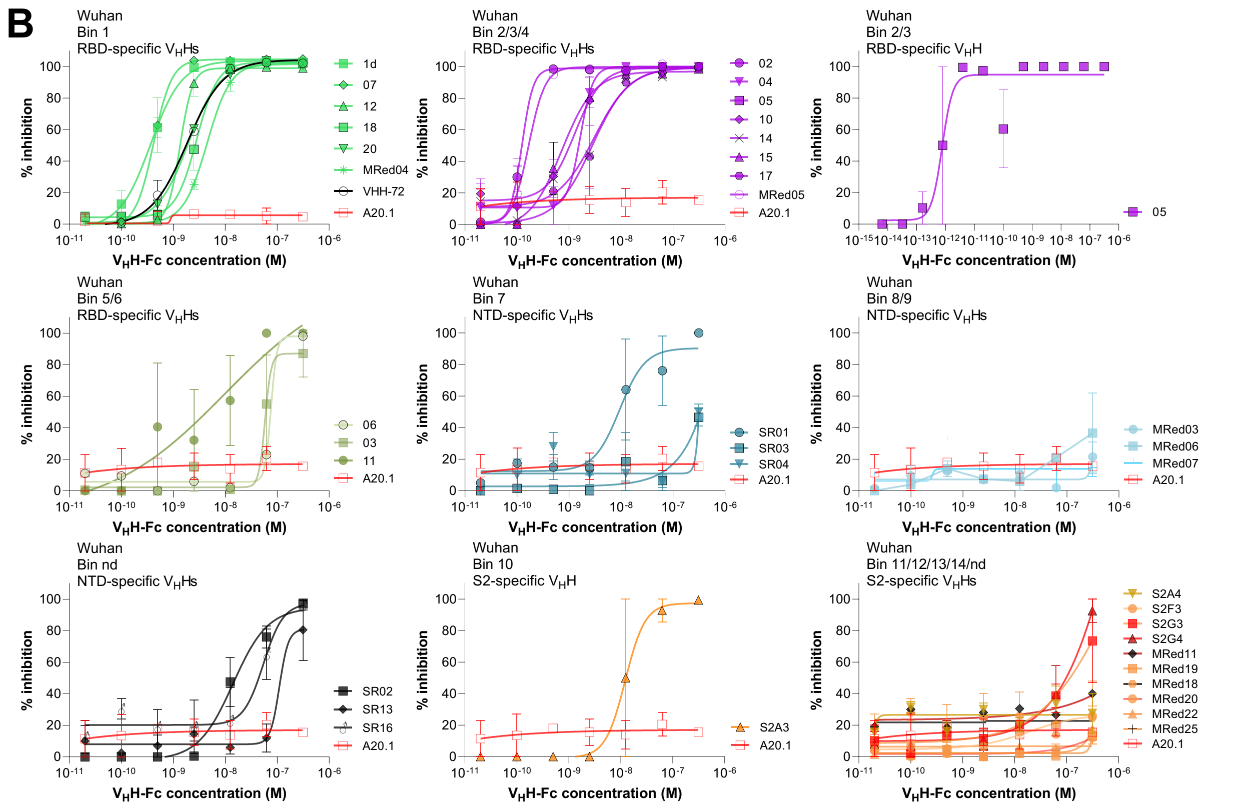


**Fig. S14. Live virus neutralization assays (LVNAs) assessing the ability of SARS-CoV-2 $V_H$H-Fcs in blocking the infection of ACE2-expressing Vero E6 cells by SARS-CoV-2 Wuhan variant at fixed (A) or varying (B) $V_H$H-Fc concentrations. A)** Inhibition assays were performed at 312, 12.5 or 0.5 nM $V_H$H-Fc concentrations. $IC_{50}$s calculated from graphs in **B** are reported in **Table 2**. VHH-72[1] and *C. difficile* toxin A-specific $V_H$H A20.1[2] are included as a benchmark and negative antibody control, respectively. $V_H$H-Fcs are color-coded according to their epitope bin designation (*see* **Fig. 5A**).

**Figure S15**

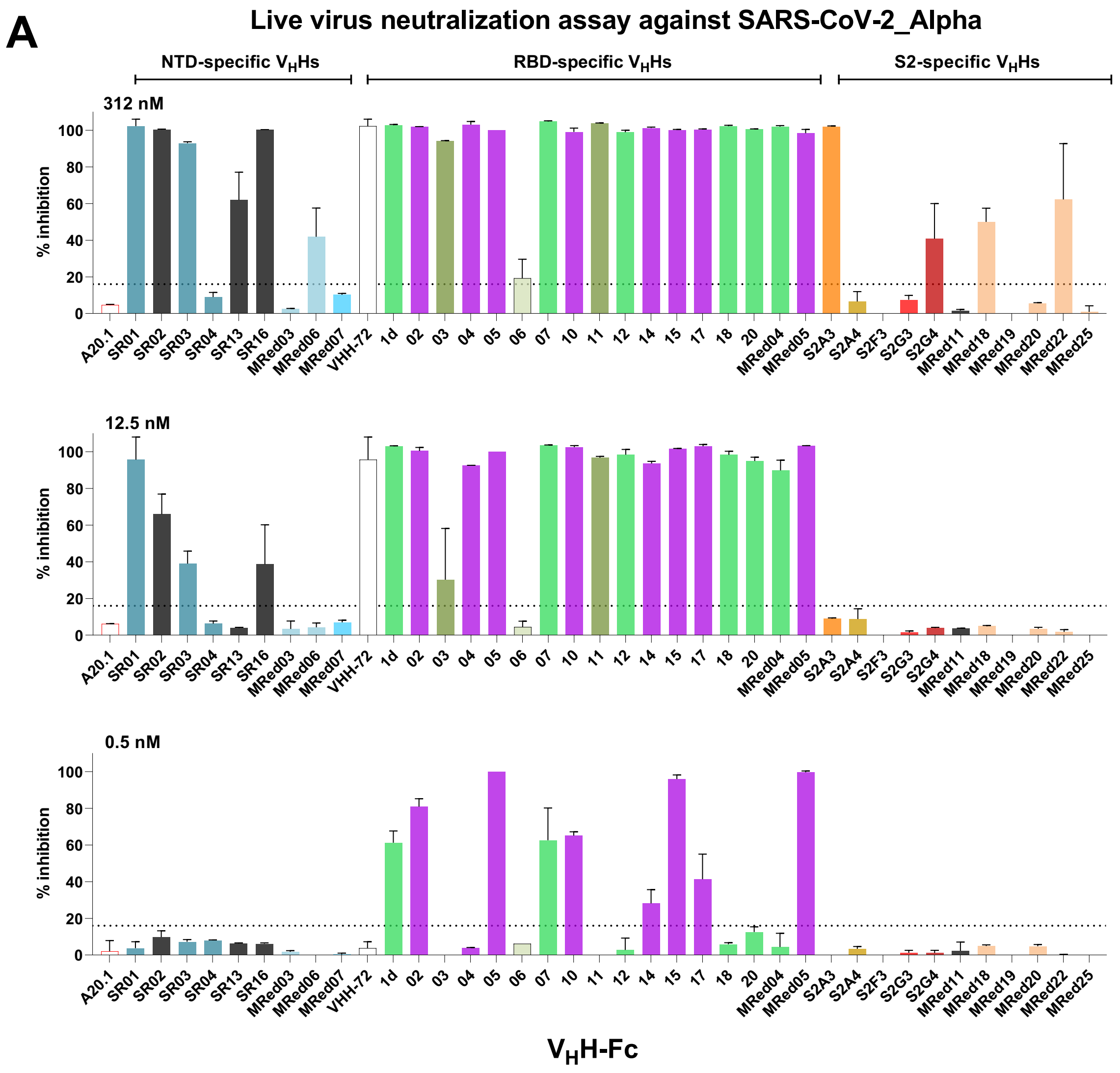
A
Live virus neutralization assay against SARS-CoV-2_Alpha
NTD-specific VHHs
RBD-specific VHHs
S2-specific VHHs
312 nM
12.5 nM
0.5 nM
% inhibition
A20.1
SR01
SR02
SR03
SR04
SR13
SR16
MRed03
MRed06
MRed07
VHH-72
1d
02
03
04
05
06
07
10
11
12
14
15
17
18
20
MRed04
MRed05
S2A3
S2A4
S2F3
S2G3
S2G4
MRed11
MRed18
MRed19
MRed20
MRed22
MRed25
VHH-Fc

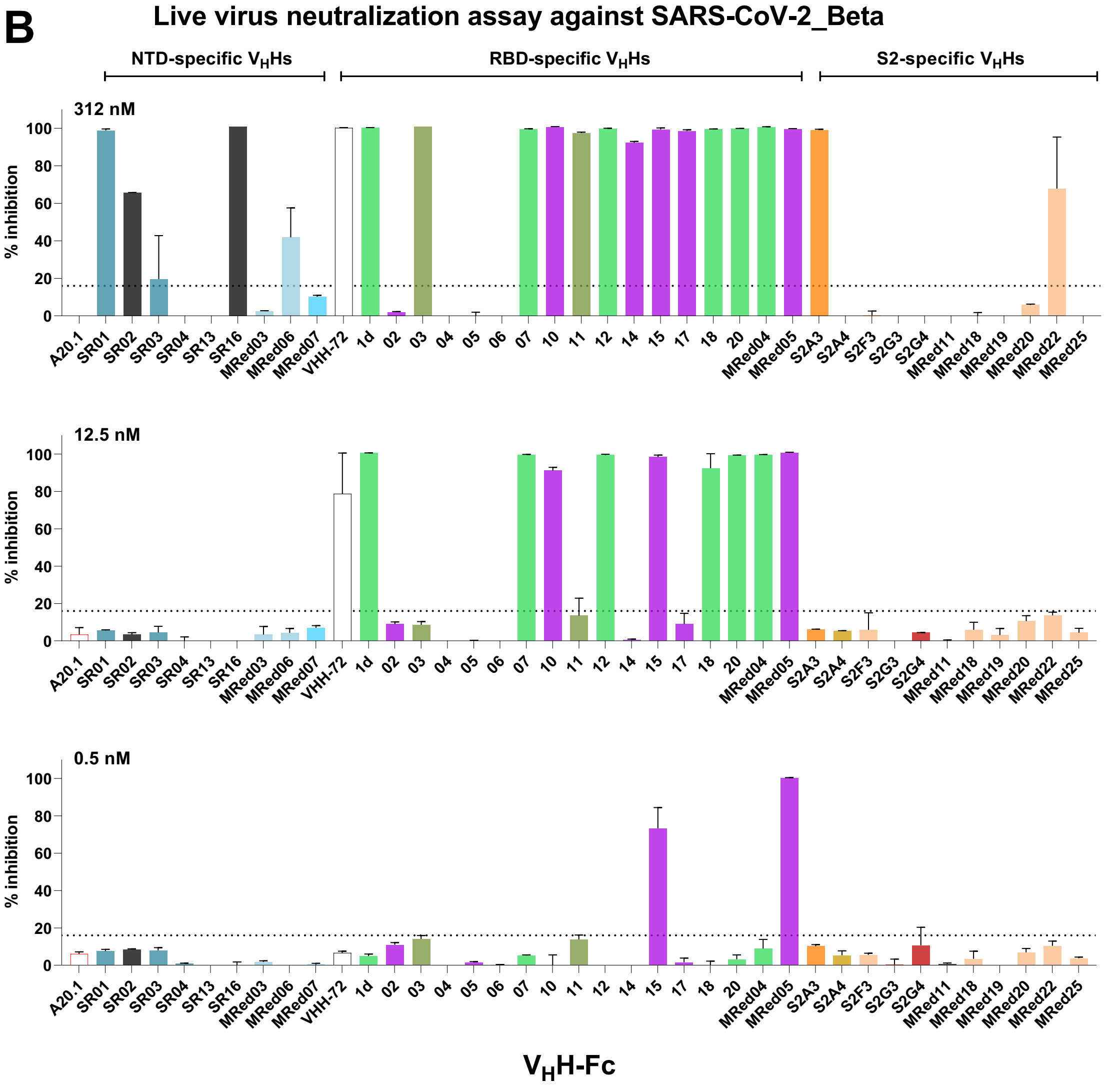
B
Live virus neutralization assay against SARS-CoV-2_Beta
NTD-specific $V_H$Hs
RBD-specific $V_H$Hs
S2-specific $V_H$Hs
312 nM
12.5 nM
0.5 nM
% inhibition
A20.1
SR01
SR02
SR03
SR04
SR13
SR16
MRed03
MRed06
MRed07
VHH-72
1d
02
03
04
05
06
07
10
11
12
14
15
17
18
20
MRed04
MRed05
S2A3
S2A4
S2F3
S2G3
S2G4
MRed11
MRed18
MRed19
MRed20
MRed22
MRed25
$V_H$H-Fc

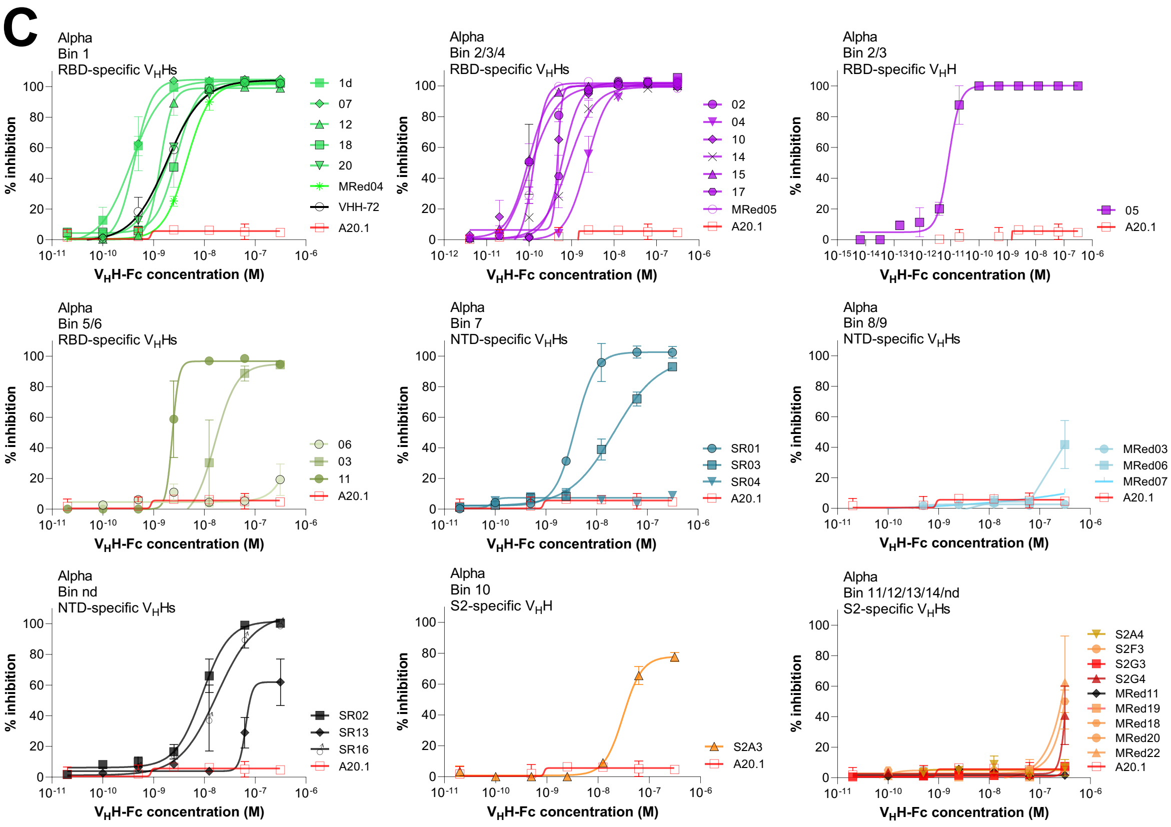
C
Alpha
Bin 1
RBD-specific $V_H$Hs
1d
07
12
18
20
MRed04
VHH-72
A20.1
Alpha
Bin 2/3/4
RBD-specific $V_H$Hs
02
04
10
14
15
17
MRed05
A20.1
Alpha
Bin 2/3
RBD-specific $V_H$H
05
A20.1
Alpha
Bin 5/6
RBD-specific $V_H$Hs
06
03
11
A20.1
Alpha
Bin 7
NTD-specific $V_H$Hs
SR01
SR03
SR04
A20.1
Alpha
Bin 8/9
NTD-specific $V_H$Hs
MRed03
MRed06
MRed07
A20.1
Alpha
Bin nd
NTD-specific $V_H$Hs
SR02
SR13
SR16
A20.1
Alpha
Bin 10
S2-specific $V_H$H
S2A3
A20.1
Alpha
Bin 11/12/13/14/nd
S2-specific $V_H$Hs
S2A4
S2F3
S2G3
S2G4
MRed11
MRed19
MRed18
MRed20
MRed22
A20.1
% inhibition
$V_H$H-Fc concentration (M)

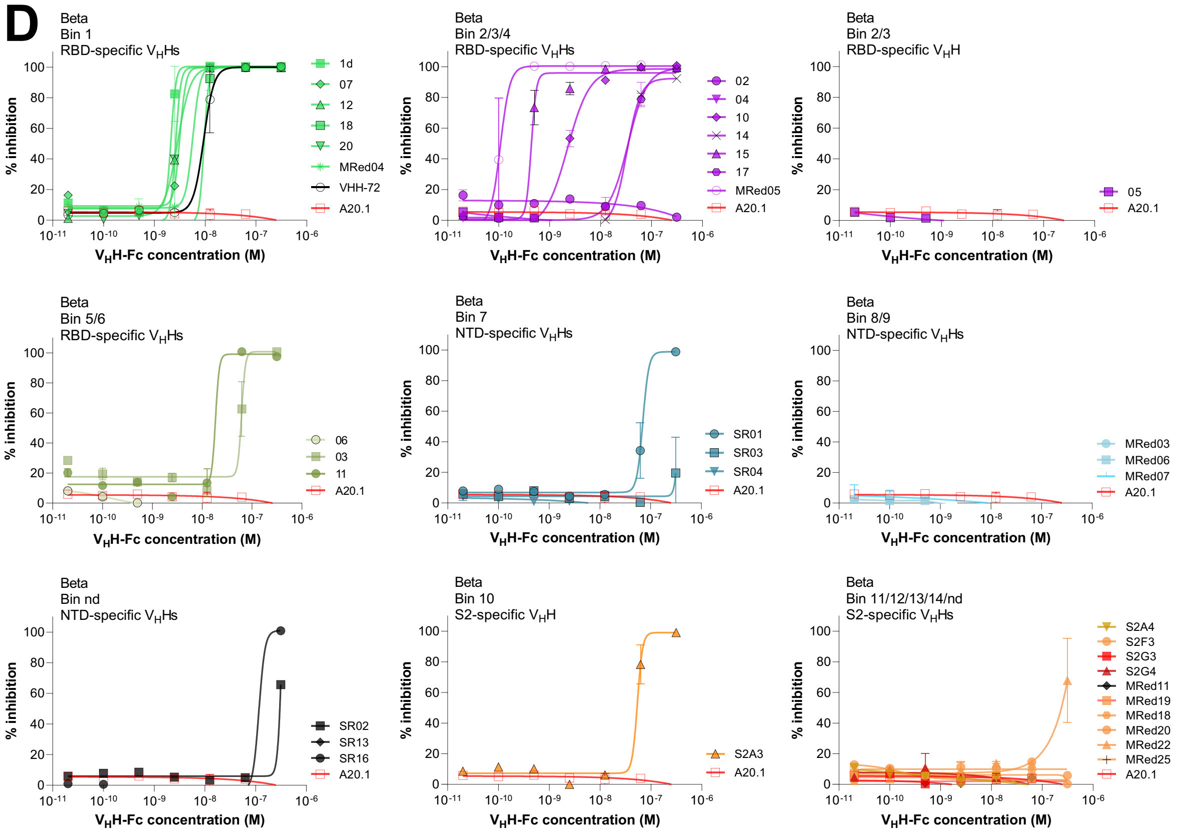


**Fig. S15. LVNAs assessing the ability of SARS-CoV-2 $V_H$H-Fcs in blocking the infection of ACE2-expressing Vero E6 cells by SARS-CoV-2 Alpha (A and C) and Beta (B and D) variants at fixed (A and B) or varying (C and D) $V_H$H-Fc concentrations. A and B)** Inhibition assays were performed at 312, 12.5 or 0.5 nM $V_H$H-Fc concentrations. $IC_{50}$s calculated from graphs in **C** and **D** are recorded in **Table 2**. VHH-72[1] and *C. difficile* toxin A-specific $V_H$H A20.1[2] are included as a benchmark and negative antibody control, respectively. $V_H$H-Fcs are color-coded according to their epitope bin designation (*see* **Fig. 5A**).

Figure S16

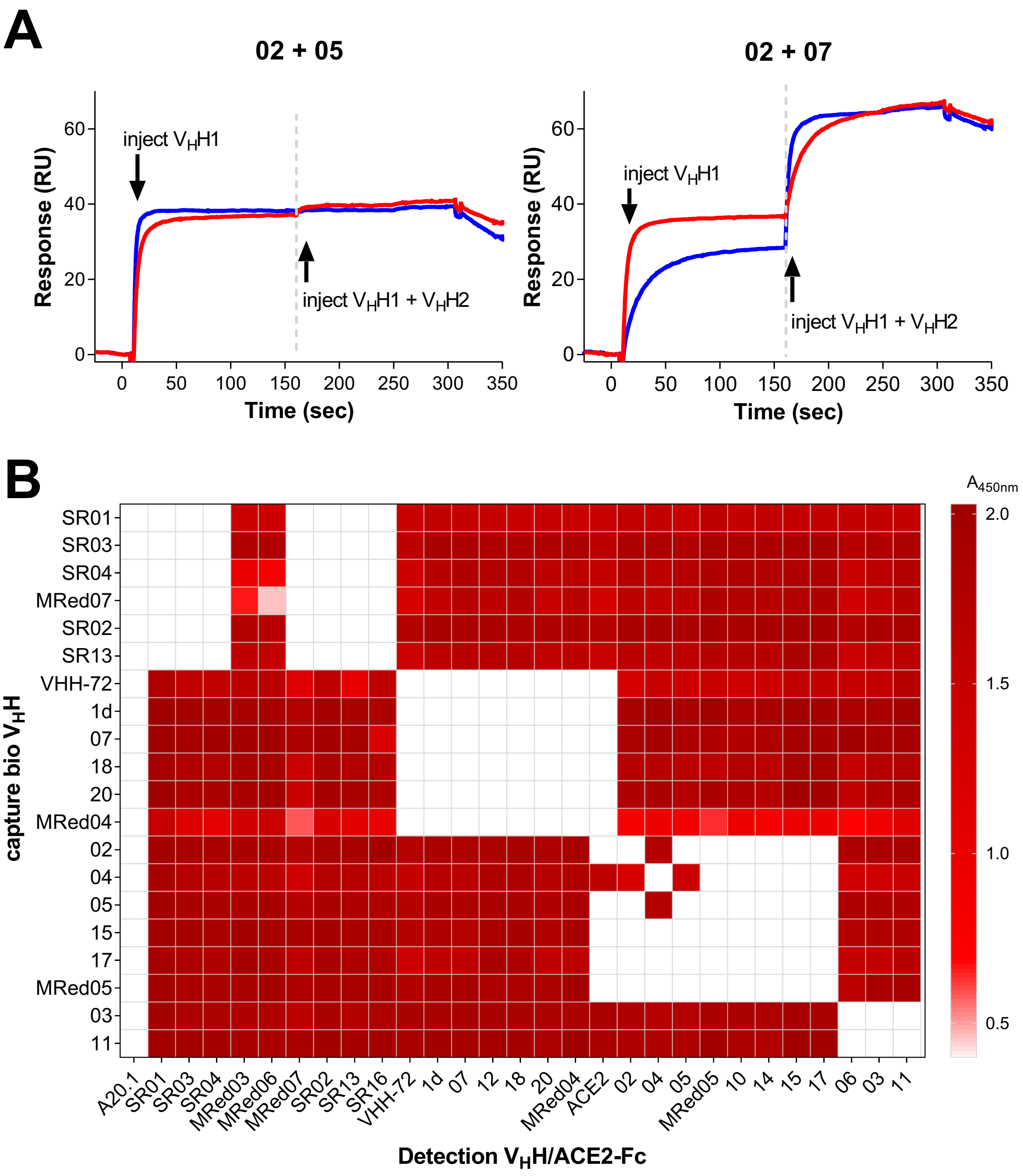
A
02 + 05
02 + 07
inject VHH1
inject VHH1 + VHH2
Response (RU)
Time (sec)
B
A450nm
capture bio VHH
Detection VHH/ACE2-Fc
SR01
SR03
SR04
MRed07
SR02
SR13
VHH-72
1d
07
18
20
MRed04
02
04
05
15
17
MRed05
03
11
A20.1
MRed03
MRed06
SR16
12
ACE2
10
14
06

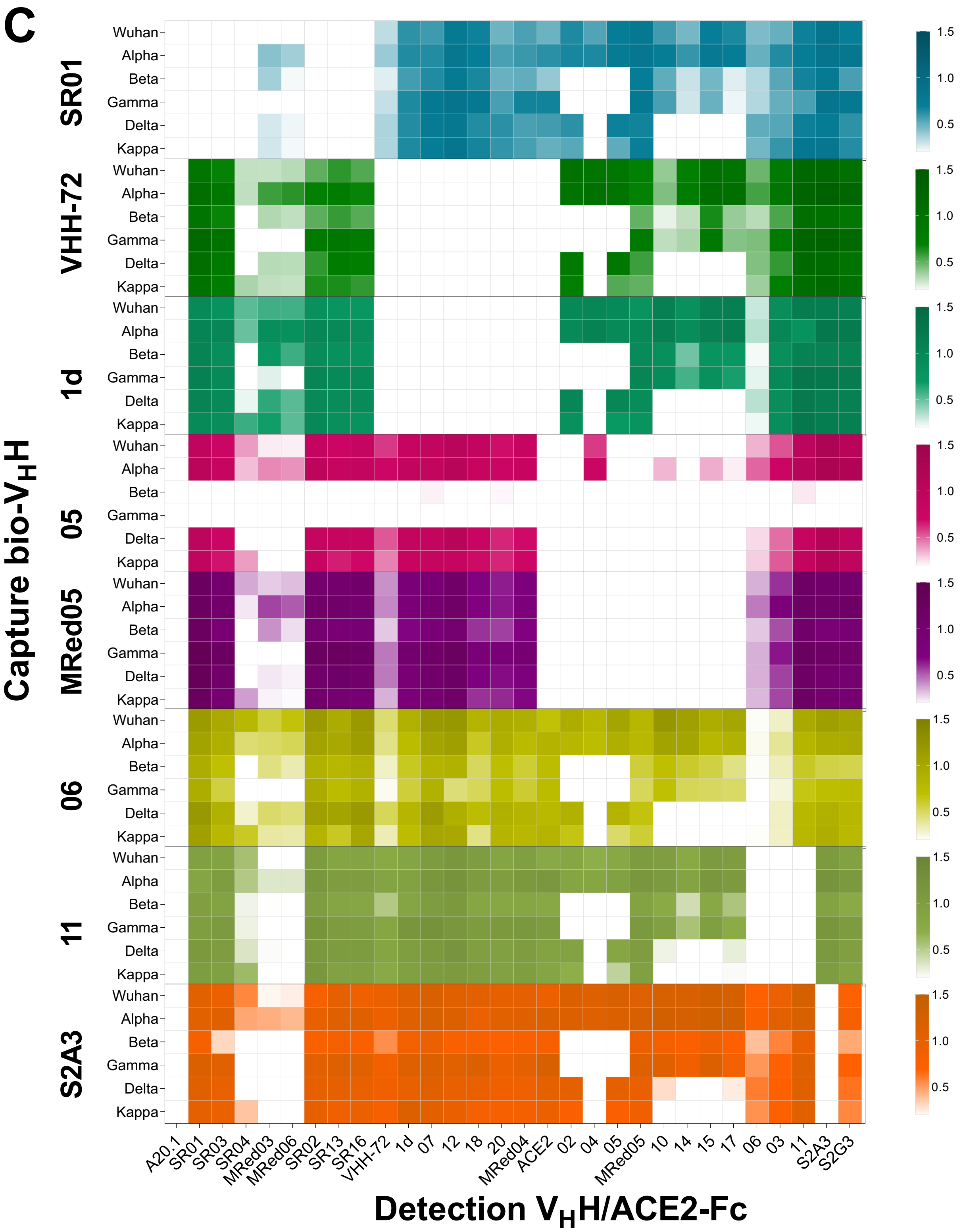


**Fig. S16. Epitope binning of SARS-CoV-2 $V_HHs$. A**) Epitope binning by SPR co-injections. Representative sensorgrams showing SPR epitope binning on SARS-CoV-2 Wuhan S-immobilized surface. For each test pair (02/05 or 02/07), the first $V_HH$ ($V_HH1$) was injected over immobilized S at saturating antigen binding concentration (50x $K_D$) followed by a second injection containing a mixture of the first $V_HH$ and a second $V_HH$ ($V_HH1$ + $V_HH2$), both at saturating concentrations (50x $K_D$). Assays were performed in two orientation formats: injecting $V_HH1$ first (orientation #1; blue profile) or $V_H$H2 first (orientation #2; red profile), followed by injecting $V_H$H1 + $V_H$H2 mix. The 02/05 profile represents $V_H$H pairs that are interpreted to bind to overlapping epitopes, hence belonging to the same epitope bin, since the injection of the second $V_H$H does not result in significant increase in binding (RU) over that already achieved by the injection of the first $V_H$H. Conversely, the 02/07 profile represents $V_H$H pairs that are interpreted to bind to non-overlapping epitopes and belong to the different epitope bins, since the addition of the second $V_H$H results in significant increase in binding over that achieved by the injection of the first $V_H$H. **B**) Epitope binning by competitive sandwich ELISA. Biotinylated $V_H$Hs (bio $V_H$Hs) were captured on streptavidin-coated wells, followed by the addition of SARS-CoV-2 Wuhan S1. The second, detecting $V_H$H sets were added as bivalent $V_H$H-Fcs, followed by the addition of HRP-conjugated anti-Fc antibody to probe $V_H$H-Fc binding to S1. ELISA binding results for pair-wise combinations of $V_H$Hs against S1 are presented as heat map. Binding pairs giving a binding signal (color) were considered as recognizing non-overlapping epitopes and belong to different epitope bins, while those giving no/weak binding signals (colorless/pale color) were considered recognizing overlapping epitopes and belong to the same epitope bin. **C**) Leads from various bins (SR01 [anti-NTD],1d, 05, MRed05, 06, 11 [anti-RBD], and S2A3 [anti-S2]) as well as VHH-72 benchmark were subjected to sandwich ELISA using S from Wuhan, Alpha, Beta, Gamma, Delta and Kappa variants. Epitope binning results are summarized in **Fig. 5A**.

**Figure S17**

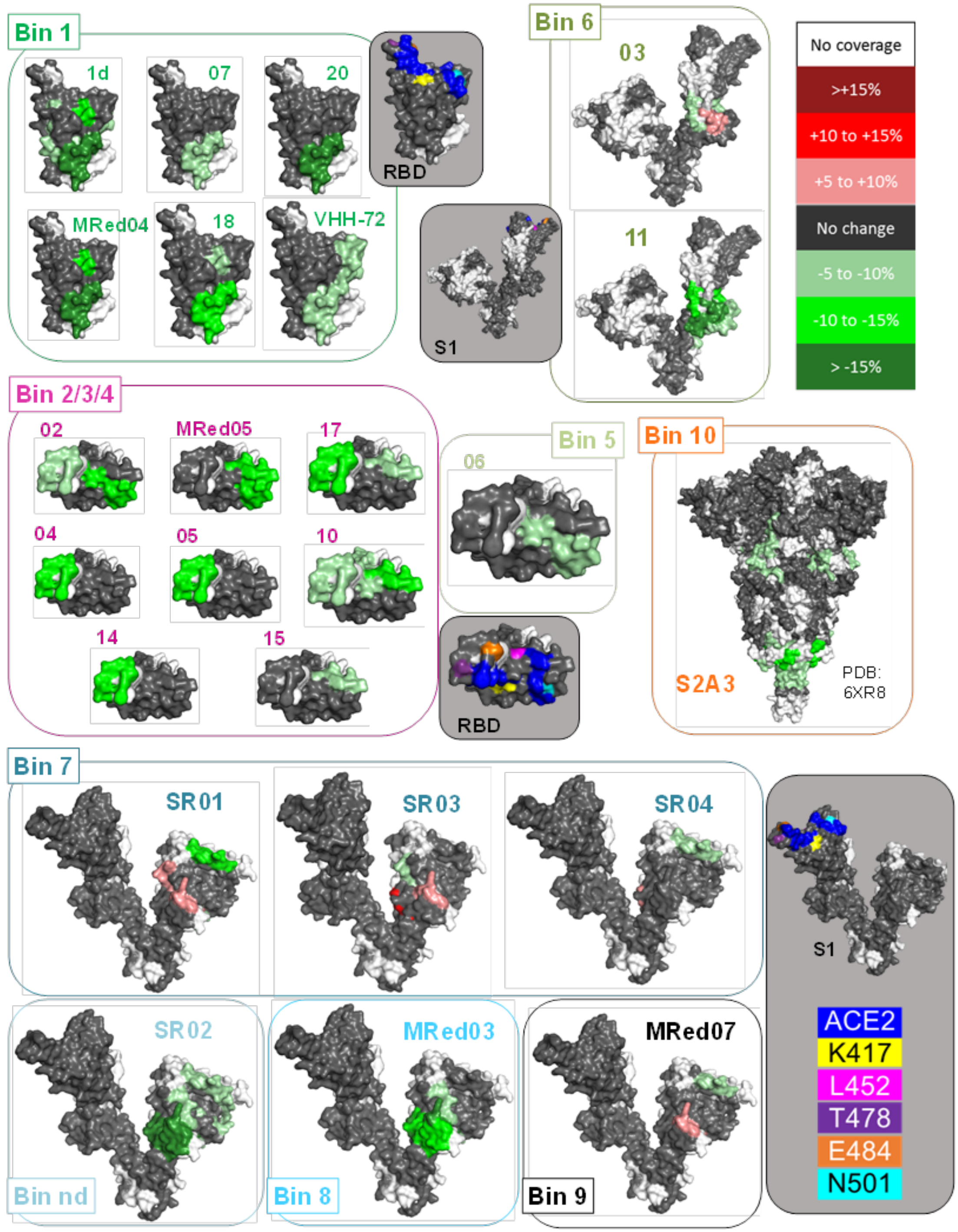


**Fig. S17. Visualization of differential HDX.** ΔD measurements ($D_{bound}$-$D_{control}$) projected onto three-dimensional structures (PDB 6XR8). Bins are colored based on **Fig 5A**. Stabilizations are shown in green, and destabilizations in red, while regions with no significant changes in deuteration are shown in grey and missing coverage in white. Key structural features are highlighted in shaded grey boxes. Here, the ACE2 binding site is shown in blue, and five mutations from VoCs over the RBD and S1 domain are included for reference (*see* legend).

**Figure S18**

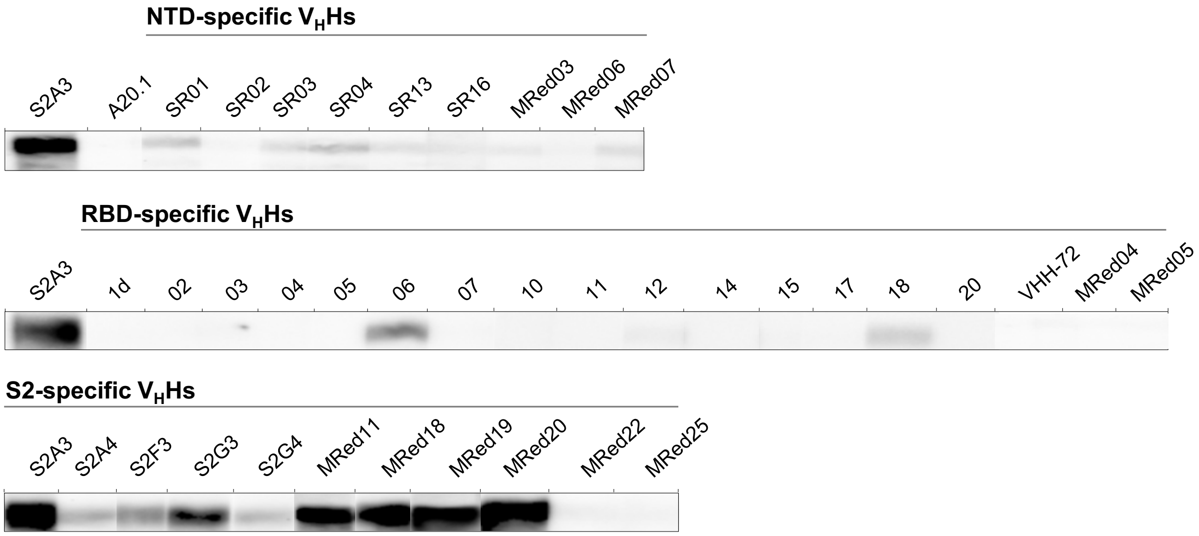


**Fig. S18. Epitope typing of SARS-CoV-2 $V_H$Hs by SDS-PAGE/western blot against denatured S**. Presence of blots (binding signals) indicate $V_H$H-Fcs recognizes a linear epitope. The absence of binding signals is an indirect indication of $V_H$H-Fcs recognizing conformational epitopes. A20.1[2] $V_H$H-Fc was included as isotype control. As expected, the conformational epitope-specific VHH-72[1] shows no binding towards denatured S. S2A3 was included in each row as reference. Epitope types are reported in **Table 1**.

**Figure S19**

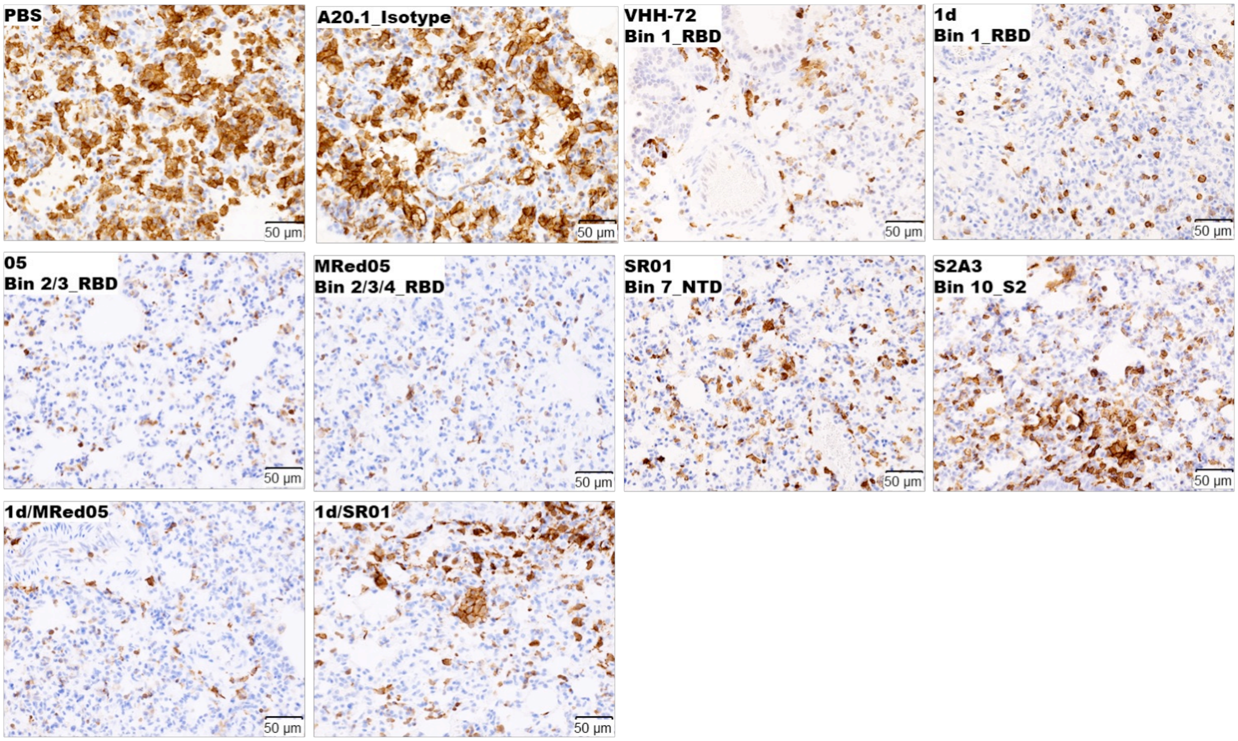


**Fig. S19. Immunohistochemical detection of infiltrating macrophages in the lungs of $V_H$H-Fc-treated animals.** Untreated (PBS) and A20.1[2] isotype-treated animals showed an intense immune reaction to anti-Iba-1 antibody and an increased number of Iba-1-positive macrophages in the consolidated areas. A substantial reduction in the number of Iba-1-positive macrophages was seen in the perivascular areas and pulmonary interstitium in the lungs of $V_H$H-Fc-treated animals. Representative images are shown from a single experiment.

**Figure S20**

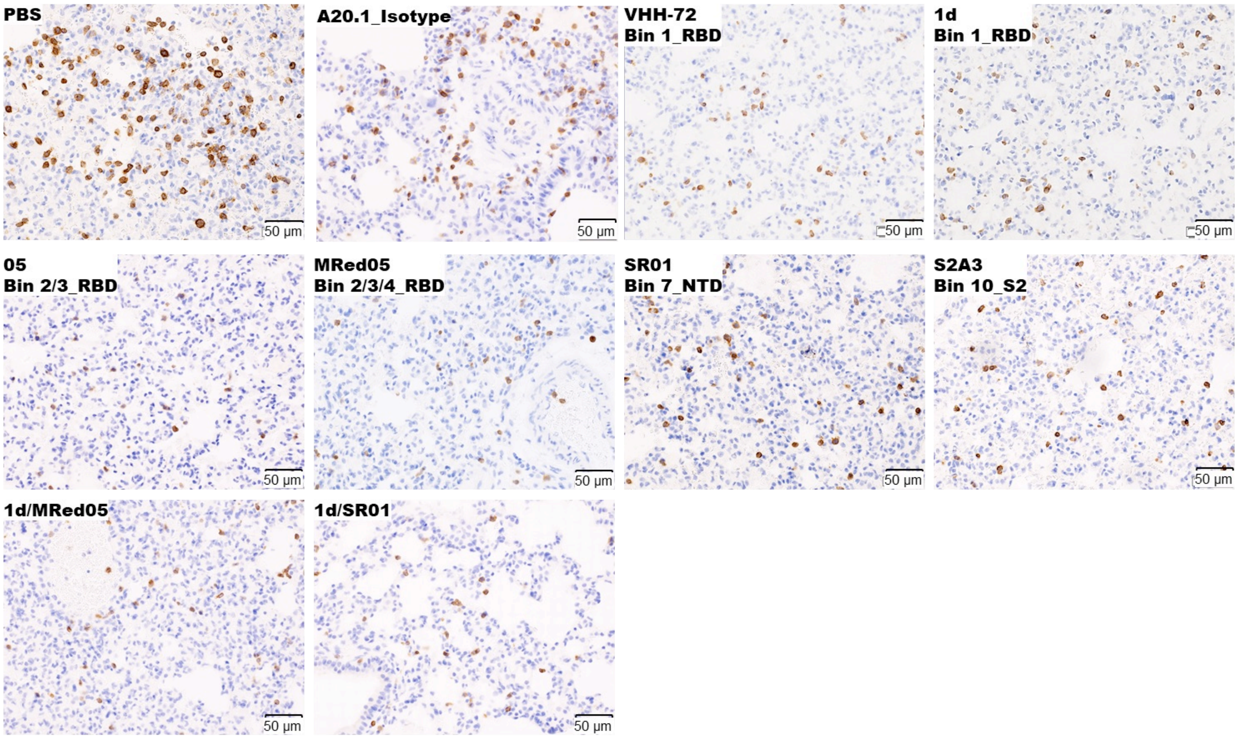


**Fig. S20. Immunohistochemical detection of T lymphocytes in the lungs of $V_H$H-Fc-treated animals.** Untreated (PBS) and A20.1[2] isotype-treated animals showed an increased number of T lymphocytes in the pulmonary interstitium. A dramatic decrease in the number of T lymphocytes was seen in the lungs of $V_H$H-Fc-treated animals. Representative images are shown from a single experiment.

**Figure S21**

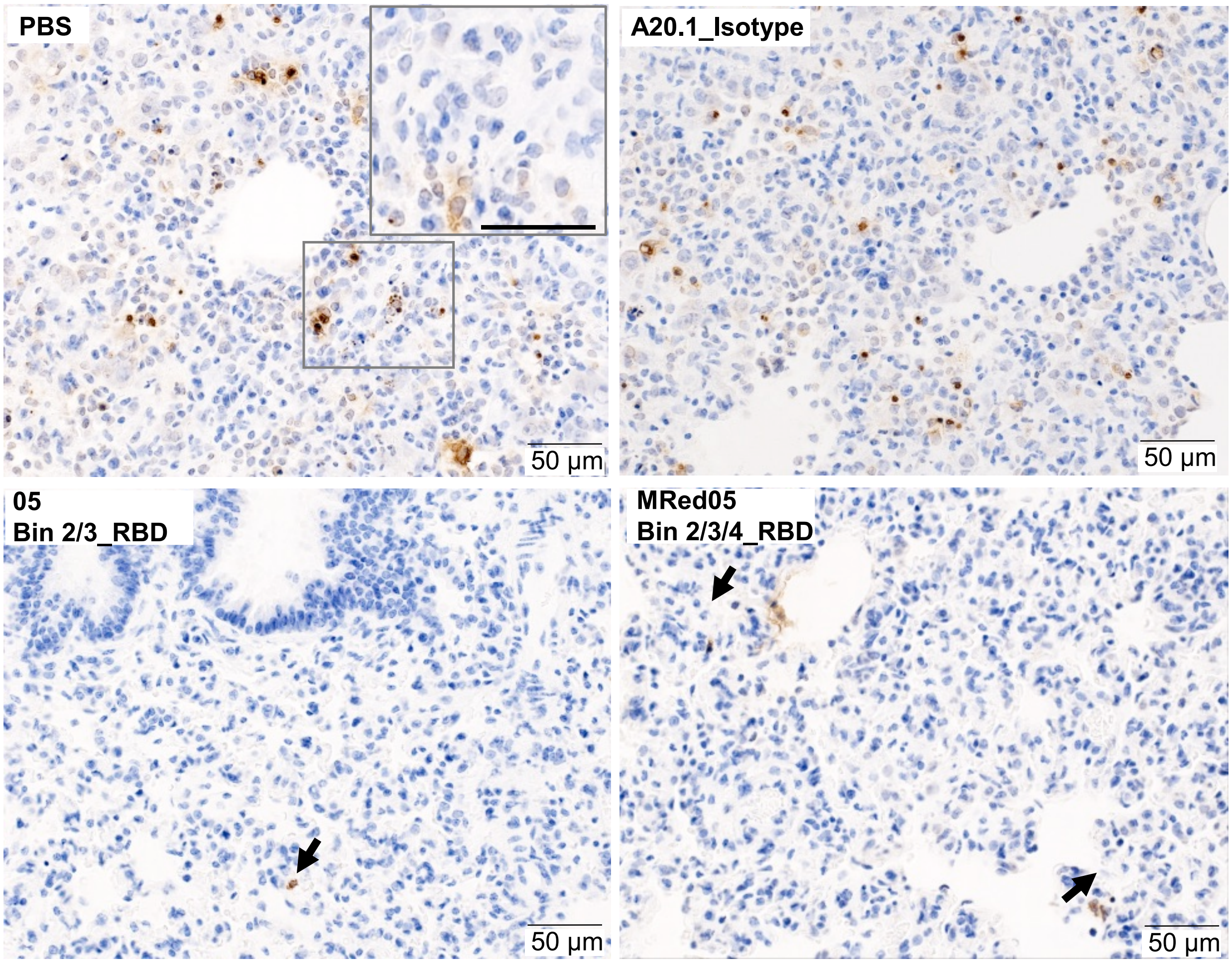


**Fig. S21. Immunohistochemical detection of apoptotic cells in the lungs of $V_H$H-Fc-treated animals.** Untreated (PBS) and A20.1[2] isotype-treated animals showed an increase in the number of TUNEL-positive cells with classical features of apoptotic cells in the pulmonary interstitium. The large grey frame in the corner of PBS panel shows the magnification of the region (small grey frame) in the lung parenchyma, scale bar = 50 µm. A marked reduction in the TUNEL-positive cells was seen in the lungs of 05- and MRed05-treated animals. Black arrows indicate occasional TUNEL-positive cells. Representative images are shown from a single experiment.
